# Macromolecular Diamidobenzimidazole Conjugates Activate STING

**DOI:** 10.1101/2025.01.21.634206

**Authors:** Karan Arora, Taylor L. Sheehy, Jacob A. Schulman, Blaise R. Kimmel, Caitlin McAtee, Vijaya Bharti, Alissa M. Weaver, John T. Wilson

## Abstract

Pharmacologic activation of the stimulator of interferon genes (STING) pathway has broad potential applications, including the treatment of cancer and viral infections, which has motivated the synthesis and testing of a diversity of STING agonists as next generation immunotherapeutics. A promising class of STING agonists are the non-nucleotide, small molecule, dimeric-amidobenzimidazoles (diABZI), which have been recently used in the synthesis of polymer- and antibody-drug conjugates to improve pharmacokinetics, modulate biodistribution, and to confer other favorable properties for specific disease applications. These approaches have leveraged diABZI variants functionalized with reactive handles and enzyme-cleavable linkers at the 7-position of the benzimidazole for conjugation to and tunable drug release from carriers. However, since this position does not interact with STING and is exposed from the binding pocket when bound in an “open lid” configuration, we sought to evaluate the activity of macromolecular diABZI conjugates that lack enzymatic release and are instead conjugated to polymers via a stable linker. By covalently ligating diABZI to 5 or 20 kDa mPEG chains via an amide bond, we surprisingly found that these conjugates could activate STING *in vitro.* To further evaluate this phenomenon, we designed a diABZI-functionalized RAFT chain transfer agent that provided an enabling tool for synthesis of large, hydrophilic, dimethylacrylamide (DMA) polymers directly from a single agonist and we found that these conjugates also elicited STING activation *in vitro* with similar kinetics to highly potent small molecule analogs. We further demonstrated the *in vivo* activity of these macromolecular diABZI platforms, which inhibited tumor growth to a similar extent as small molecule variants. Using flow cytometry and fluorescence microscopy to evaluate intracellular uptake and distribution of Cy5-labeled analogs, our data indicate that although diABZI-DMA conjugates enter cells via endocytosis, they can still colocalize with the ER, suggesting that intracellular trafficking processes can promote delivery of endocytosed macromolecular diABZI compounds to STING. In conclusion, we have described new chemical strategies for the synthesis of stable macromolecular diABZI conjugates with unexpectedly high immunostimulatory potency, findings with potential implications for the design of polymer-drug conjugates for STING agonist delivery that also further motivate investigation of endosomal and intracellular trafficking as an alternative route for achieving STING activation.

## Introduction

The stimulator of interferon genes (STING) pathway is an ancient and evolutionarily conserved innate immune sensing mechanism with critical roles in pathogen detection, tumor immune surveillance, and maintenance of tissue homeostasis.^1^ STING is a transmembrane protein predominantly localized on the endoplasmic reticulum (ER) and is activated upon the binding of several cyclic dinucleotides (CDNs), including 2’3’cGAMP, the endogenous STING ligand that is synthesized by the enzyme cGAS in response to detection of cytosolic double stranded DNA.^2^ ^3^ Activation of STING triggers the secretion of type-I interferons (IFNs) and other proinflammatory cytokines that promote immune responses against viruses, bacteria, and cancers.^4^ ^5^ Accordingly, STING agonists are an emerging and promising class of therapeutics with broad applications as antiviral agents, cancer immunotherapies, and vaccine adjuvants.^6^ ^7^

Although CDNs have been widely explored for diverse applications and continue to be optimized for clinical translation, their activity and efficacy is typically limited due to poor drug-like properties that results in rapid clearance, poor cellular uptake, and limited access to the cytosol for binding to STING. ^8^ ^9^ To address this challenge, several non-CDN small molecule STING agonists have recently been developed with improved membrane permeability and properties for systemic administration.^10^ ^11^ ^12^ One promising class of small molecule STING agonists are dimeric-amidobenzimidazoles (diABZI), first described by Ramanjulu et al. and under clinical investigation for immune-oncology (e.g., NCT03843359).^10^ While relatively large by conventional small molecule standards (e.g., >700 Da), diABZI compounds can diffuse across cell membranes, allowing for cytosolic access and binding to STING.^10^ Hence, diABZI compounds have increased potency compared to 2’3’ cGAMP and many other CDNs and pharmacological properties that confer improved therapeutic efficacy in mouse tumor models when administered systemically.^10^ ^13^ However, the reported serum half-life of diABZI is still relatively short (i.e., ∼90 min)^10^ and membrane permeability and lack of cell or organ (e.g., tumor) tropism results in indiscriminate STING activation with potential for inflammatory toxicities, such as cytokine release syndrome.

Akin to other classes of therapeutics (e.g., chemotherapy drugs), the development of carrier-drug conjugates has potential to overcome these pharmacological barriers.^14^ ^15^ Indeed, we and others have recently described the development of polymer- and protein-drug conjugates for improved delivery of diABZI compounds.^16^ ^17^ ^18^ ^19^ For example, we have recently described STING-activating polymer-drug conjugates (SAPCon) that are based on conjugation of diABZI to a hydrophilic polymer backbone through a cathepsin B-cleavable linker to enable intracellular diABZI release.^16^ Similarly, Nguyen et al. copolymerized diABZI-functionalized monomers into mannose-containing polymers to target macrophages, also using a cathepsin-cleavable linker.^19^ These reports exemplify the vast potential opportunities afforded through the design of diABZI compounds derivatized with reactive handles that can be ligated to diverse carriers, including not only polymers, but also proteins, lipids, peptides, and glycans, amongst others, to modulate pharmacological properties or enhance delivery to specific cells or tissues.

Central to this approach is the synthesis of diABZI variants with reactive handles installed at the 7-position of the benzimidazole which does not interact with STING and is exposed from the binding pocket when bound in an “open lid” configuration (**Figure 1A**).^20^ ^10^ By contrast, the binding of 2’3’-cGAMP and several synthetic CDNs to STING results in a “closed lid” conformation that wraps around the agonist and effectively buries it within the protein.^21^ A second important consideration is the choice of linker chemistry, which has potential to allow for diABZI release from carriers under specific intracellular or microenvironmental conditions, though, to date, only cathepsin-cleavable linkers for intracellular release have been utilized. While it is logical to integrate cleavable linkers that allow diABZI to be released from macromolecular carriers, because diABZI binds to STING in an “open lid” configuration it is also conceivable that diABZI compounds covalently linked to bulky macromolecules could still bind to and activate STING. This possibility is supported by evidence demonstrating that conjugation of diABZI to small molecules, such as fluorescent dyes and PET imaging probes can bind and activate STING.^22^ However, these molecules are still relatively small (<1000 Da) with the conjugated molecule assuming minimal steric bulk, and, therefore, it is unknown if larger, macromolecules can also be conjugated to diABZI while still allowing for STING binding and activation.

**Figure 1:**
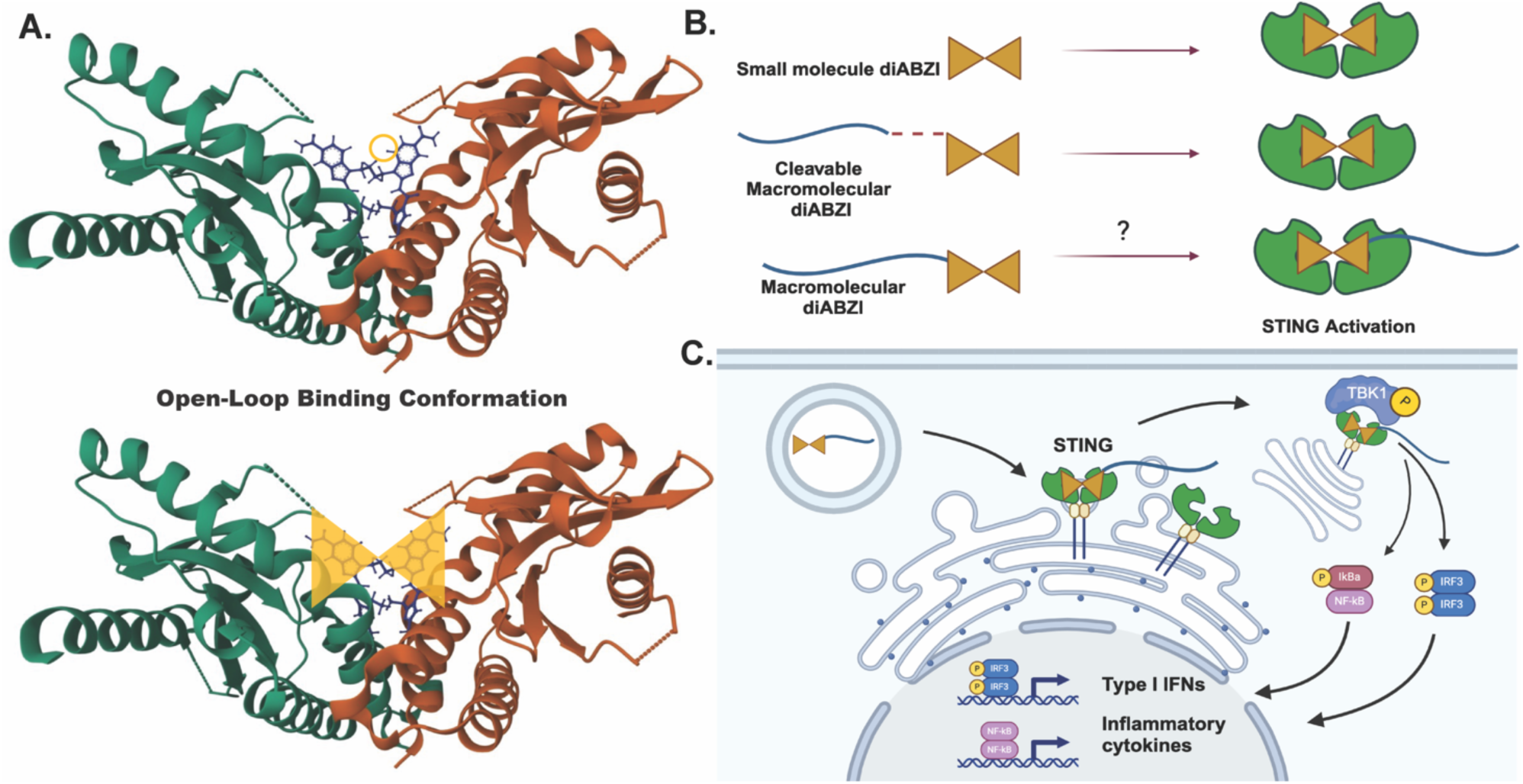
Dimeric-amidobenzimidazole (diABZI) compounds conjugated to hydrophilic macromolecules have potential to bind and activate STING. **(A)** Crystal structure of diABZI (compound 3) binding to STING demonstrating binding in an “open lid” conformation.^10^ ^20^ The 7-position of the benzimidazole (circled) extends from the binding pocket providing a potential site for covalent ligation of diABZI to large macromolecules while still allowing for STING binding and activation. **(B)** Schematic illustrating that small molecule diABZI compounds and diABZI linked to polymeric carriers via cleavable linkers can activate STING and the possibility that macromolecular diABZI conjugates may also act as STING agonists. **(C)** Schematic illustrating the concept that macromolecular diABZI conjugates can access, bind, and activate STING to stimulate type-I interferons (IFN) and proinflammatory cytokines.

Towards addressing this knowledge gap, here we asked if diABZI molecules conjugated to water soluble polymer chains via stable amide linkages could activate STING signaling (**Figure 1B-C**). To do so, we first covalently ligated diABZI to 5 or 20 kDa mPEG chains via an amide bond and, surprisingly, found that these conjugates could activate STING *in vitro*. To further explore this phenomenon, we developed a strategy for synthesis of amide-linked polymer-diABZI conjugates based on a diABZI-functionalized chain transfer agent (CTA) that allowed for growth of precisely-defined polymer chains from a single diABZI molecule via reversible addition-fragmentation chain-transfer (RAFT) polymerization. Intriguingly, we again found that chemically stable polymer-diABZI conjugates as large as 175 kDa could rapidly activate STING with similar kinetics to highly potent small molecule analogs. Additionally, these polymer-diABZI conjugates could activate STING *in vivo*, resulting in inhibition of tumor growth in mouse models. Through mechanisms that remain to be fully elucidated, we also observed that polymer-diABZI conjugates enter cells via endocytosis but can colocalize with the ER where STING is typically activated. These data suggest that diABZI covalently linked to large, hydrophilic, macromolecular carriers may access, bind, and activate STING and that this may be mediated by vesicular transport processes that allow endocytosed polymer-diABZI conjugates to be trafficked to ER. Therefore, our findings have important implications for the design of polymer-diABZI conjugates and may offer insight into how endocytic and intracellular trafficking may be harnessed to target diABZI to the ER for activation of STING.

## Results and Discussion

### Synthesis and characterization of mPEG-diABZI conjugates

We first sought to determine if covalent linkage of diABZI to a bulky, hydrophilic macromolecular carrier through a stable amide bond would allow for STING binding and *in vitro* activation. To enable these investigations, we synthesized an amine-functionalized diABZI **(1)** that provides a versatile precursor for further functionalization with other reactive handles and ligation to diverse macromolecules (**Figures S9, S25, and S31**).^16^ As a model polymeric carrier, we selected linear poly(ethylene glycol) (PEG) based on its common use as a biocompatible carrier for enhancing aqueous solubility and improving circulation half-life.^23^ ^24^ We synthesized diABZI-PEG conjugates by reacting diABZI-amine **(1)** with commercially available MeO-PEG_n_-NHS ester for nucleophilic substitution of NHS ester with diABZI-amine. This yielded PEGylated diABZI variants conjugated to PEG chains of 5 **(2)** or 20 kDa **(3)** via an amide linkage (**Scheme 1, Scheme S1**). diABZI-PEG conjugates were characterized by 1H NMR (**Figure S10-11**).

**Scheme 1:**
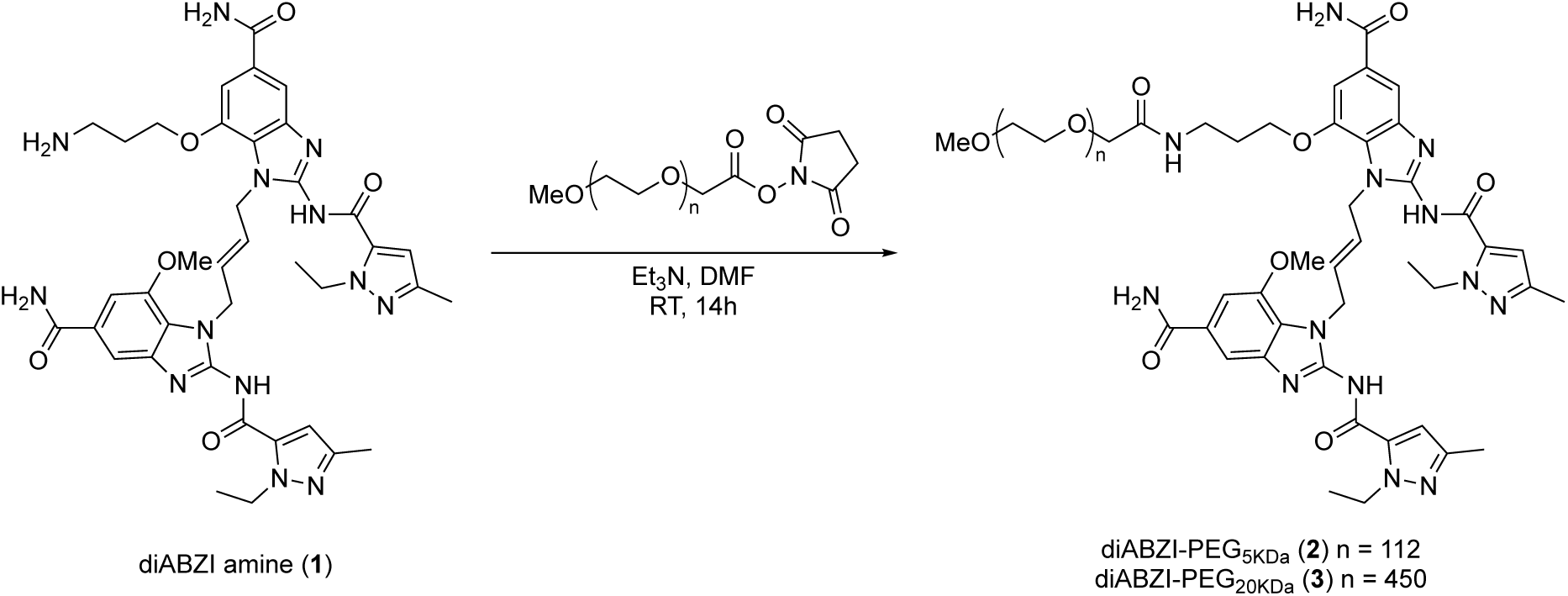
Synthesis of diABZI-PEG[5kDa] and diABZI-PEG[20kDa]

As controls, we also synthesized PEGylated analogs containing an established cathepsin-B cleavable valine-citrulline-PAB linker **(6 and 7),** a commonly used linker in antibody-drug conjugates that allows for regeneration of diABZI-amine **(1)** following capthesin-B cleavage within the endo-lysosome.^25^ ^26^ To do so, we first synthesized a maleimide functionalized diABZI with a valine-citrulline-PAB spacer (diABZI-V/C-Mal; **5**) for conjugation to thiol-functionalized carriers via thiol-maleimide Michael addition reaction (**Scheme S2)**. The compound and intermediates were characterized by NMR and ES-MS (**Figures S12, S26, and S32**). We validated the enzyme cleavability of diABZI-V/C-Mal **(5)** via incubation with cathepsin B for 48 hours followed by MALDI-MS to detect the regeneration of diABZI-amine (**1**) upon enzyme cleavage. Incubation of 50 μM diABZI-V/C-Mal (**5**) with 0.2 μM cathepsin B resulted in a shift in molecular weight from 1400.8 Da to 801.83 Da, consistent with the molecular weight of diABZI-amine **(1)** sodium adduct (**Figure S1**), whereas no change in molecular weight was observed under the same conditions for diABZI-Mal **(9)**, an analog that lacks an enzyme-cleavable spacer (**Scheme S3, Figures S15, S27, and S33**). To evaluate the activity of modified diABZ-amine compared to diABZI-V/C-Mal *in vitro*, we used human monocyte-derived THP1-Dual reporter cells that are engineered to secrete luciferase upon activation of interferon regulatory factor (IRF) signaling. Dose-response curves indicated relatively similar potency between the small molecule analogs diABZI-amine **(1)** and diABZI-V/C-Mal **(9)** with estimated EC_50_ values of 60.9 and 314 nM, respectively (**Figure 2A**). In primary murine splenocytes, which comprise a mixture of different immune cell populations, both compounds stimulated IFN-β production to a similar extent, with EC_50_ values for diABZI-amine (**1**) and diABZI-V/C-Mal (**9**) of approximately 2.24 and 3.38 μM, respectively (**Figure 2B**). The modest differences in EC_50_ values observed between the compounds in THP1 cells can likely be attributed to differences in cell membrane permeability as similar STING binding affinity was observed between the analogs. To synthesize the cathepsin B cleavable diABZI-PEG analogs, diABZI-V/C-PEG[5kDa, 20kDa], diABZI-V/C-Mal was reacted with MeO-PEG-SH under basic conditions where the terminal thiol attacks the maleimide group to form a thioether linkage (**Figure 2C**, **Scheme S2**). The PEGylated products were characterized by using 1H NMR (**Figure S13 and S14**).

**Figure 2:**
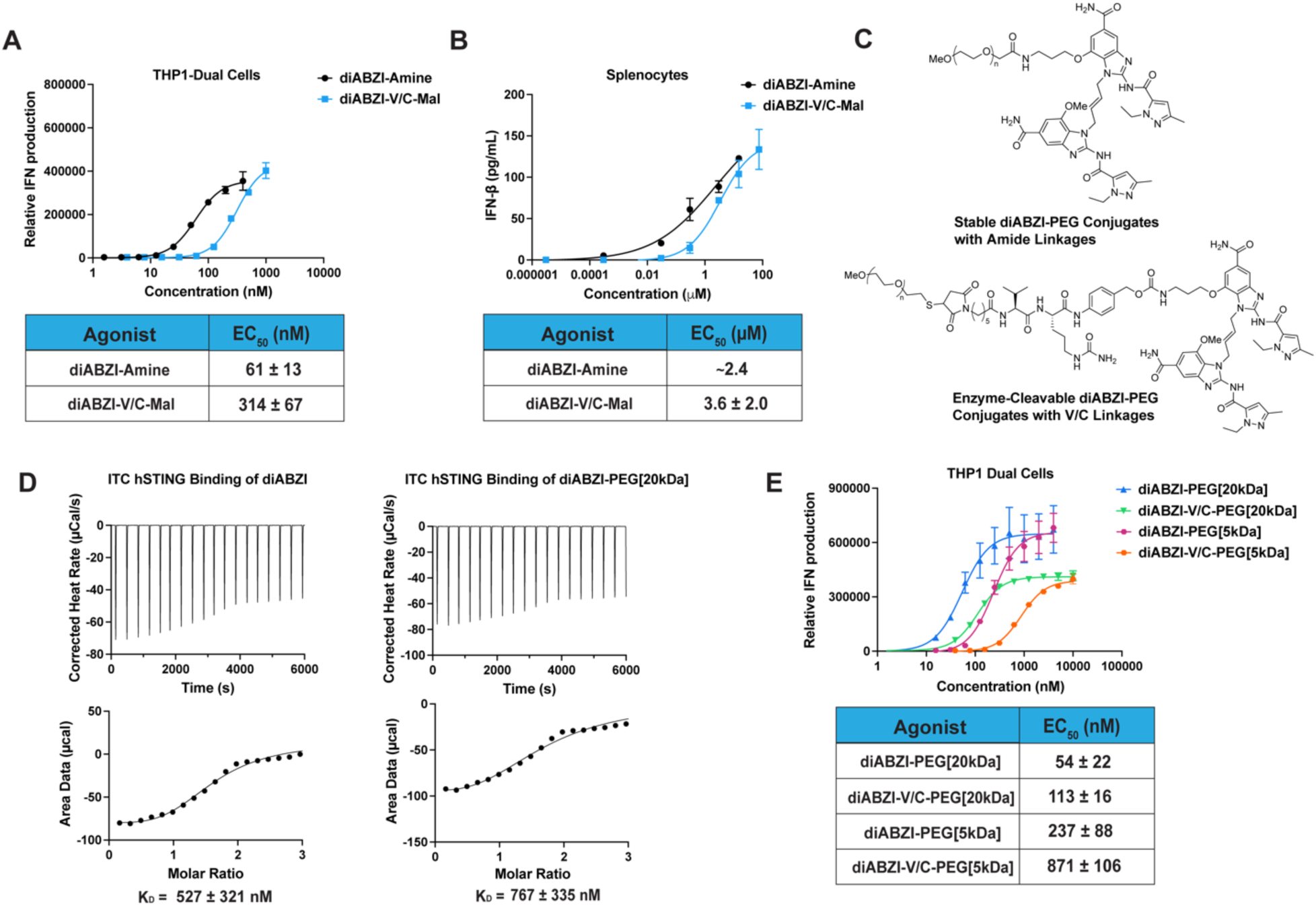
Chemical and biological characterization of polyethylene glycol (PEG) functionalized diAZBI compounds. **(A)** Dose-response curves for relative IFN-I production by THP1-Dual reporter cells treated with diABZI-Amine and diABZI-V/C-Mal (n=3). **(B)** Dose-response curves for IFN-b secretion by isolated murine splenocytes treated with diABZI-amine and diABZI-V/C-Mal (n=3). **(C)** Chemical structures of amide-linked diABZI-PEG (**2,3**) and cathepsin B cleavable diABZI-V/C-PEG (**5,6**) conjugates. **(D)** Isothermal calorimetry (ITC) traces (top) and binding isotherm (bottom) to determine a K_D_ for diABZI-PEG[20kDa] binding to human recombinant STING; commercially available diABZI (compound 3) was used as a control. **(E)** Dose-response curves for relative IFN-I production by THP1-Dual reporter cells treated with cleavable and non-cleavable diABZI-PEG conjugates (n=3). Dose-response curves were fit to a variable slope (four parameter) non-linear regression to estimate EC_50_ values.

### PEG-diABZI conjugates bind and activate STING

We next sought to understand the effects of these diABZI modifications on STING binding compared using isothermal calorimetry with recombinant human STING (STING isoform 1) to estimate binding affinity (**Figure 2D**). Using the commercially available diABZI molecule (compound 3) we determined a K_D_ of approximately 527 nM. Interestingly, when evaluating the binding of macromolecular diABZI-PEG[20kDa], we did not observe a significant differences in the K_D_ compared to the free diABZI compound, suggesting that the diABZI moiety can still access its STING binding domain even with a large PEG chain attached.

Having validated the activity of the functionalized small molecule diABZI variants, we next sought to understand the effect of diABZI PEGylation on STING activation *in vitro* by comparing the activity of diABZI-PEG conjugates with 5 and 20kDa PEGs linked via the stable or enzyme cleavable linker. Surprisingly, STING activation was similar between non-cleavable and enzyme cleavable PEG conjugates, with non-cleavable analogs exhibiting slightly higher activity (**Figure 2E**) and activity also increasing mildly when conjugated to the longer 20kDa PEG chain regardless of linker chemistry. In all cases, no activity was observed in STING knockout THP1 cells, confirming the dependence on STING for activation of IRF3 signaling (**Figure S2**). Together, these findings suggest that diABZI can access, bind, and activate STING even when conjugated to a 20kDa PEG chain via a stable amide linkage. This may be a distinctive feature of diABZI since other STING agonists, such as 2’3’-cGAMP or MSA-2, trigger a closed-lid conformation that may preclude the addition of steric bulk; additionally, these molecules lack sites for chemical modification that are not also essential for STING binding.^21^ Notably, PEG-diABZI conjugates were readily water soluble, which may afford a practical advantage for drug formulation and intravenous administration as solubilization of hydrophobic diABZI compounds requires the use of viscous and/or organic excipients that may not be suitable for clinical use and/or increase the risk of adverse effects (e.g., Cremophor®).^27^

### Synthesis and characterization of diABZI-functionalized polymers via RAFT polymerization

To further validate these findings and to determine if the observed phenomenon could be extrapolated to other types of polymers with increased molecular weight, we devised a strategy for synthesizing well-defined polymers containing a single diABZI molecule linked to the end of a polymer chain via a stable amide linkage. Central to our approach was the synthesis of diABZI-functionalized-ethylsulfanylthiocarbonyl-sulfanylvpentanoic acid (ECT) RAFT chain transfer agent, which we postulated would serve as a versatile tool for controlled free-radical polymerization of bespoke polymers of defined properties containing a single chain terminal diABZI molecule. To synthesize diABZI-ECT (**11**), the carboxylic acid of ECT was converted into an activated PNP-ester using DCC-DMAP coupling and treated with diABZI-amine to obtain the desired product as diABZI-ECT (**Scheme 2, Scheme S4)**. The diABZI-ECT compound and intermediates were characterized by NMR and ES-MS (**Figures S16, S28, and S34**).

We then used diABZI-ECT to synthesize a series of diABZI-functionalized-poly(dimethyl acrylamide) (DMA) polymers, a biocompatible, water soluble polymer that has been explored as a drug carriers and for production of hydrogels.^28^ ^29^ ^30^ A distinct advantage of a CTA-based approach is the ability to synthesize diABZI-functionalized polymers with a defined degree of polymerization over a large range of molecular weights from a single diABZI molecule. Here, we synthesized diABZI-poly(DMA) with molecular weights of ∼25 **(12)**, 50 **(13)**, and 175 kDa **(14)**. The polymerization was performed in N,N-Dimethylformamide (DMF) under inert atmosphere at 70 °C at a ratio of initial monomer : diABZI-ECT: initiator of n:1:0.2 (**Scheme 2, Scheme S5**). The resultant polymers were purified by extensive dialysis, lyophilized, and characterized by ^1^H NMR and GPC/LS (**Figures S17-19 and S38**). Monomer conversion for the 25 kDa, 50 kDa, and 175 kDa diABZI-poly(DMA) constructs were 90.1%, 98.1%, and 99.3%, respectively. GPC/LS analysis showed monodisperse diABZI-poly(DMA) constructs, with PDI values of 1.01 for the 25 kDa, 1.08 for the 50 kDa, and 1.17 for the 175 kDa variant. While herein we only leveraged diABZI-ECT for synthesis of poly(DMA) and employed a stable amide linkage, the functionalization of RAFT chain transfer agents with diABZI, or other STING agonists, provides a facile chemical tool for the synthesis of well-defined polymers for activation of STING signaling.

**Scheme 2:**
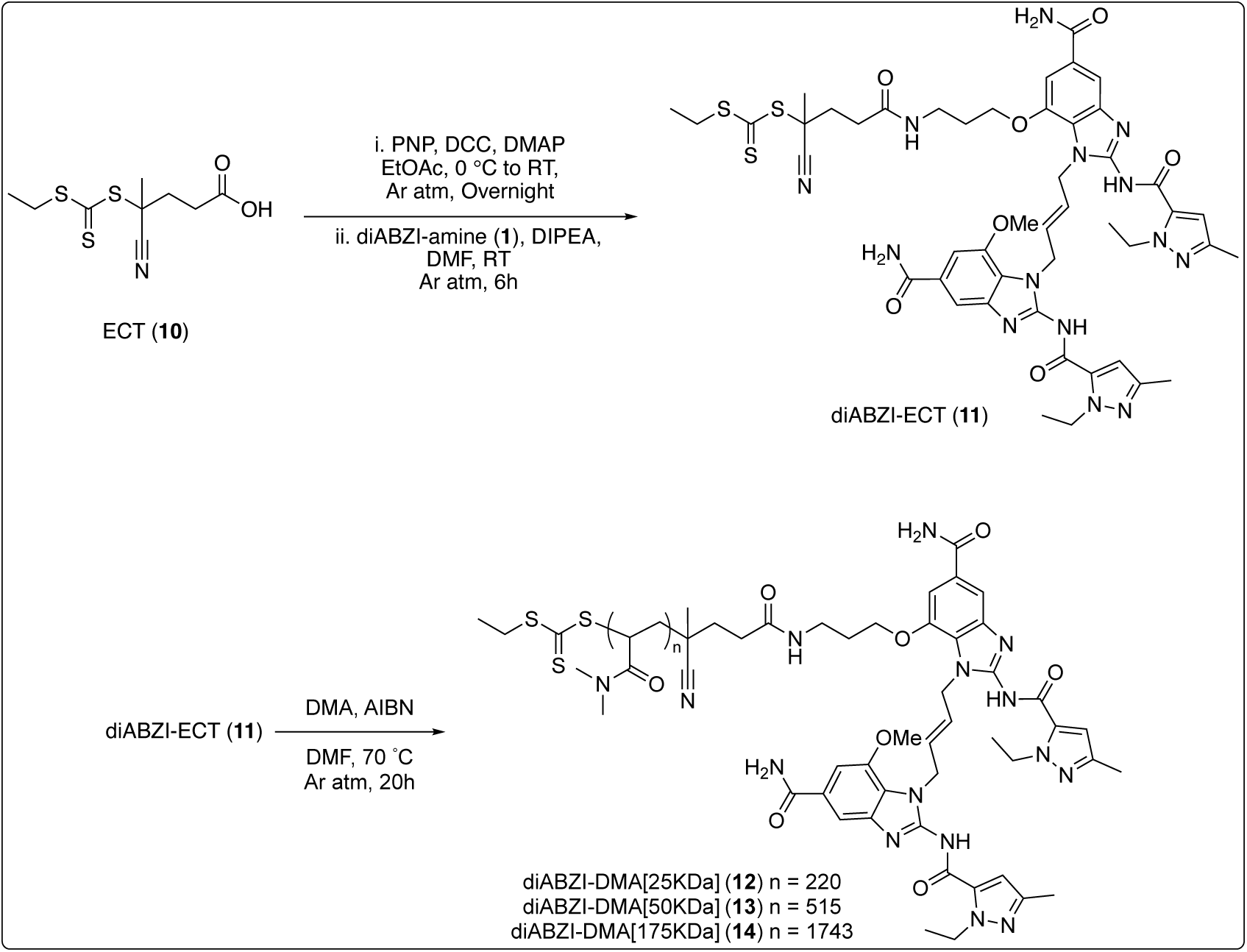
Synthesis of diABZI-ECT and diABZI-DMA[25kDa],[50kDa], and [175kDa]

### High molecular weight diABZI-DMA conjugates activate STING

We next evaluated the capacity of diABZI-DMA conjugates to activate STING in THP1-Dual reporter cells compared to the unpolymerized diABZI-ECT small molecule. Consistent with our findings with stable diABZI-PEG conjugates, we found that all stable diABZI-DMA constructs activated STING to a similar degree with EC_50_ values in the nanomolar range, regardless of polyDMA molecular weight (**Figure 3A**). As expected, we also found that the diABZI-ECT was significantly more potent than diABZI-amine, likely owing to differences in polarity and membrane permeability. Additionally, the ∼25-50-fold difference in potency between diABZI-ECT and diABZI-DMA suggests the potential to achieve prodrug-like and/or environmentally responsive STING activation through integration of a cleavable linker that liberates a potently active diABZI analog into the extracellular space. Since there were not significant differences between polymers of varying molecular weights, we selected the 50kDa diABZI-DMA for the remaining studies. Since STING activation also activates NF-κB signaling, we utilized the dual-reporter functionality of THP1-Dual reporter cells to evaluate the relative NF-κB response, finding that both diABZI-ECT and diABZI-DMA could activate this arm of the STING pathway with diABZI-ECT exhibiting higher potency as expected (**Figure S3A-B**). We also verified that responses to diABZI-ECT and diABZI-DMA were STING-dependent by evaluating activity in STING-knockout THP1 Dual reporter cells (**Fig S3C-D**). Collectively, these data further demonstrate that diABZI conjugated to bulky, water-soluble macromolecules via a stable amide linkage can still activate STING signaling and, remarkably, with a similar potency to the amine-functionalized parent compound, diABZI-amine, despite a ∼150x difference in molecular weight.

**Figure 3:**
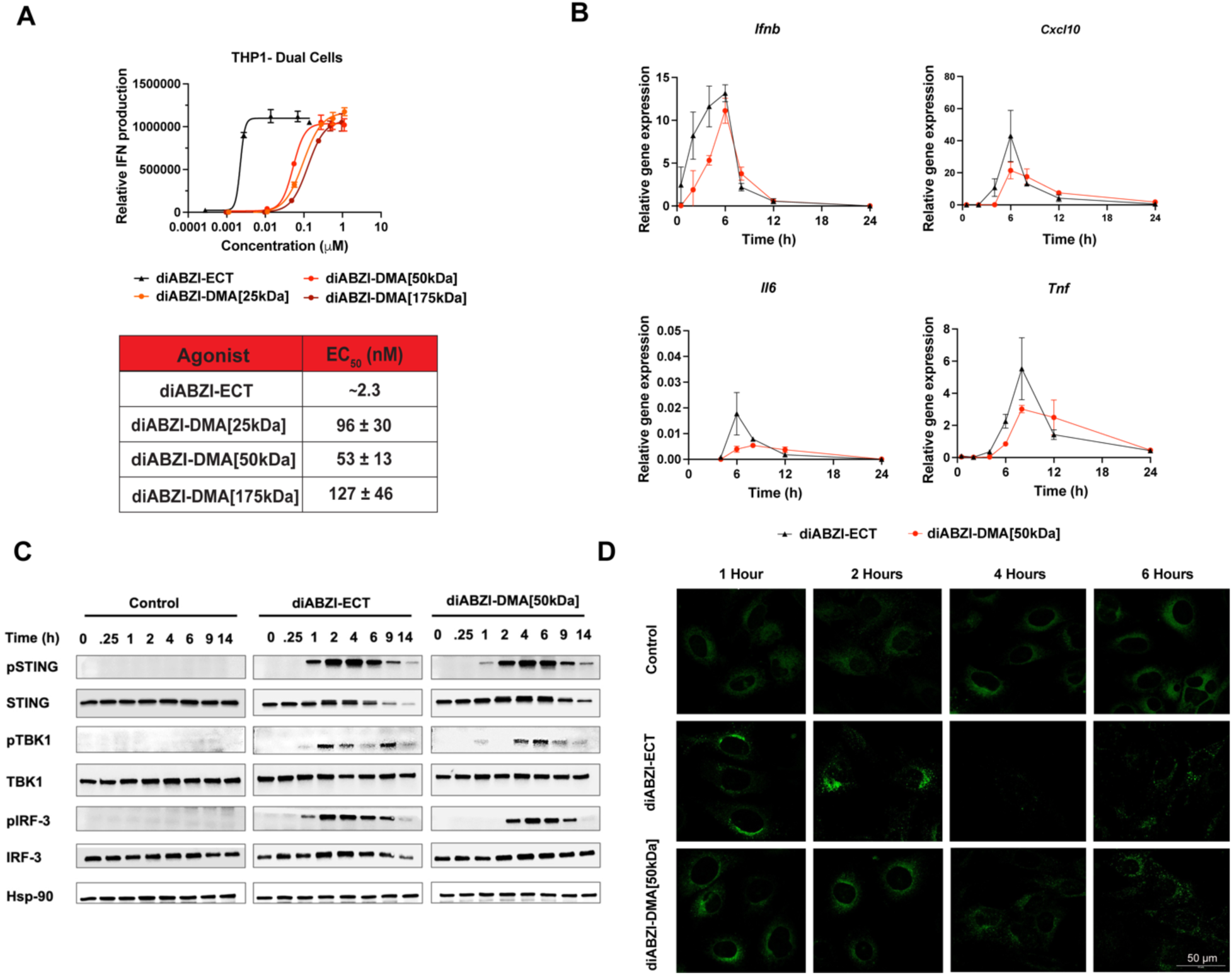
Evaluation of STING activation by diABZI-DMA conjugates. **(A)** Dose-response curves for relative IFN-I production by THP1-Dual reporter cells treated with diABZI-ECT and diABZI-DMA constructs of varying molecular weight. Dose-response curves were fit to a variable slope (four parameter) non-linear regression to estimate EC_50_ values. **(B)** qRT-PCR of STING-associated genes in response to treatment of THP1 Dual cells with diABZI-ECT or 50kDa diABZI-DMA as a function of incubation time. **(C)** Western blot analysis of STING pathway in response to treatment of THP1 Dual cells with diABZI-ECT or 50kDa diABZI-DMA as a function of incubation time. **(D)** Representative fluorescent micrographs of MEF cells expressing a STING-GFP fusion protein treated with diABZI-ECT or diABZI-DMA for indicated time. The formation of GFP puncta corresponds to multimerization of STING following ligand binding.

We next aimed to better understand STING activation kinetics between a small molecule and macromolecular diABZI variant, comparing diABZI-ECT and diABZI-DMA in THP1 Dual cells. qRT-PCR was utilized to measure the expression of STING associated genes over time by THP1 cells treated with 5 μM diABZI (**Figure 3B**). Interestingly, the magnitude and kinetics of gene expression was relatively similar between diABZI-ECT and diABZI-DMA, with gene expression of *Ifnb* and *Cxcl10* peaking around 6 h despite dramatic differences in molecular weight and postulated cellular uptake mechanisms. Treatment with diABZI-DMA resulted in slightly delayed gene expression with some prolongation of *Il6* and *Tnf* expression relative to diABZI-ECT, but inflammatory gene expression returned to baseline levels by 24 h in all cases. To further understand the kinetics of STING activation, we also performed western blot analysis for phosphorylated STING, TBK1, and IRF3 in THP1 Dual cells treated with 5 μM diABZI-ECT or diABZI-DMA. Following activation, STING traffics from the ER to the Golgi where it recruits TBK1 which phosphorylates itself, STING, and IRF3.^31^ Consistent with gene expression data, we found that diABZI-DMA results in slightly delayed STING activation as measured by levels pSTING, pTBK1, and pIRF3, which is evident within one hour for both treatments and peaks between 2-4 h for diABZI-ECT and between 4-6 h for diABZI-DMA (**Figure 3C**). Finally, to qualitatively assess intracellular STING dynamics in the initial activation period, we employed murine embryonic fibroblasts (MEF) expressing a STING-GFP fusion protein that has been used previously to evaluate the formation of fluorescent puncta associated with oligomerization of STING dimers on the ER following activation.^32^ After treating with 5 μM diABZI-ECT or diABZI-DMA, we used confocal microscopy to image relative puncta formation and GFP expression and degradation over time. Consistent with western and PCR analysis, diABZI-ECT triggered puncta formation within 1-2 h whereas this was slightly delayed with diABZI-DMA, which nonetheless activated STING within 2-4 h (**Figure 3D**). Collectively, these studies demonstrate that diABZI-DMA conjugates, linked via a stable amide bond, can activate STING with similar, though slightly delayed, kinetics relative to free small molecule diABZI compounds.

### Analysis of intracellular uptake and ER binding of diABZI-DMA conjugates

Intrigued by the findings that diABZI-DMA conjugates could rapidly and potently activate STING, we further evaluated their intracellular uptake and localization using sCy5 and Cy5-labeled variants of diABZI-Amine and diABZI-DMA **(16 and 18)** synthesized as described in **Scheme S5-6** and characterized in **Figures S20-21, S35, and S39**. The activity of Cy5-labeled variants in THP1 Dual reporter cells was confirmed in **Figure S4**. First, using flow cytometry we found that cellular uptake of diABZI-DMA-Cy5 is entirely inhibited by incubation at 4°C **(Figure 4A)**, indicating internalization via an active endocytic process, which was expected for a large hydrophilic macromolecule that cannot diffuse through the cell membrane. Uptake of diABZI-sCy5 was only partially inhibited at 4°C, which may reflect a decrease in membrane permeability at lower temperatures and/or labeling with sCy5 which reduces hydrophobicity and membrane permeability. This was further assessed by confocal microscopy of STING-GFP MEF cells treated with diABZI-DMA-Cy5 and diABZI-sCy5 and staining with LysoTracker^TM^ to identify late endosomes and lysosomes, where STING is ultimately trafficked to and degraded following activation.^31^ Colocalization of diABZI-DMA-Cy5 with LysoTracker^TM^ was significantly higher than diABZI-sCy5, further demonstrating that polymer-diABZI conjugates are internalized by endocytosis to a greater extent than free small molecule diABZI compounds, which can enter cells via passive diffusion across the cell membrane (**Figure 4B-D,S5**).

**Figure 4:**
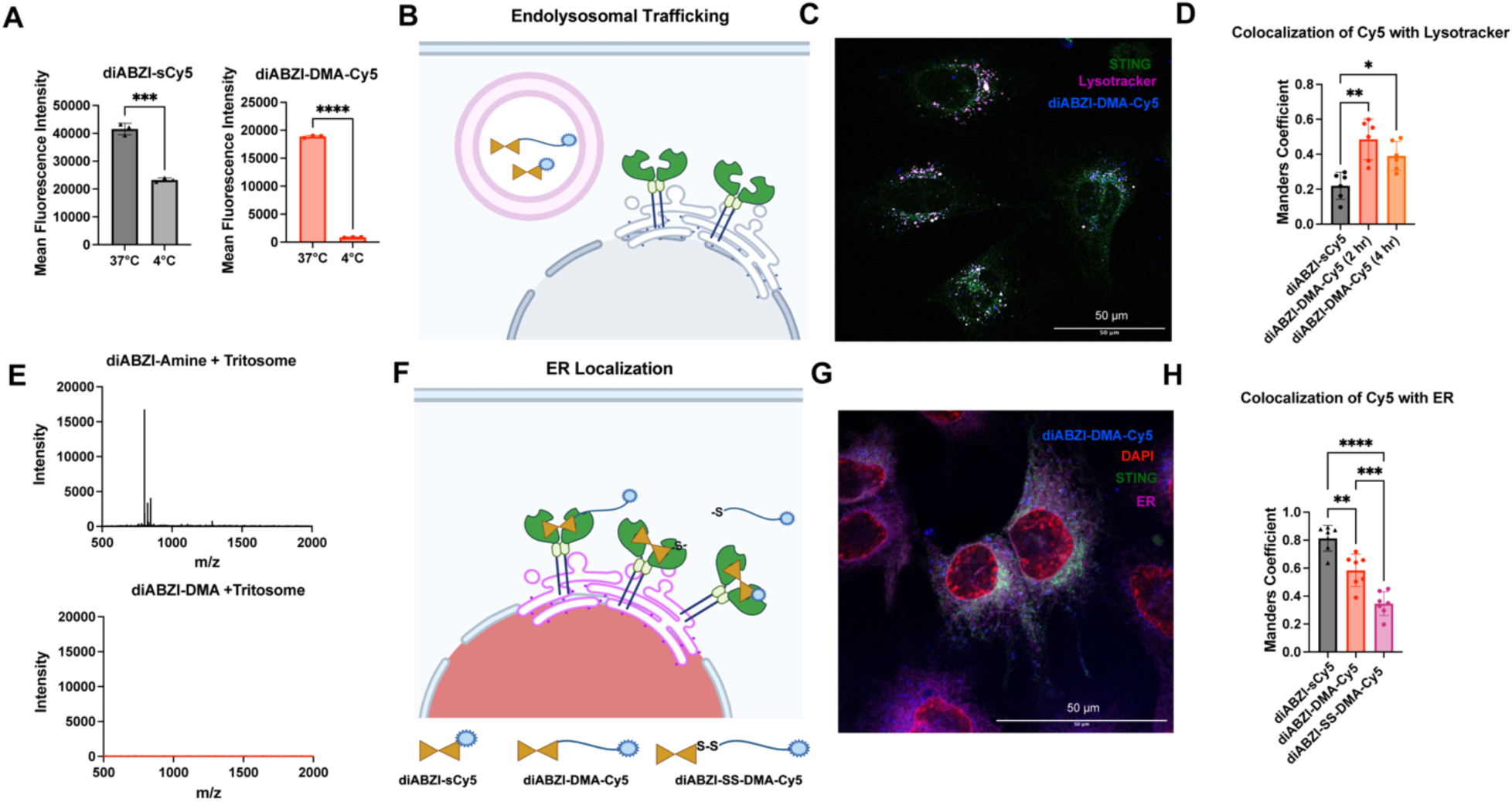
Intracellular uptake and ER localization of diABZI-DMA conjugates. **(A)** Flow cytometry analysis of THP1 Dual cell uptake of diABZI-sCy5 and diABZI-DMA-Cy5 at 37°C and 4°C to determine dependence on endocytosis (n=3). **(B)** Schematic depiction of diABZI-sCy5 and diABZI-DMA-Cy5 localized within an endolysomal compartment resulting in colocalization of Cy5 and Lysotracker in STING-GFP MEFs. **(C)** Representative confocal image of STING-GFP MEFs treated with diABZI-DMA-Cy5 for 2 hours and stained with Lysotracker Red; representative images of cells treated with diABZI-sCy5 are shown in **Figure S5**. **(D)** Quantification of colocalization between labeled diABZI variants and Lysotracker Red described by Manders coefficient (n=6). **(E)** MALDI-MS spectra (500-2000 m/z) of diABZI-DMA and diABZI-amine following incubation with rat liver tritosomes for 7h at 37°C demonstrating lysosomal stability of diABZI-DMA. **(F)** Schematic depiction of diABZI-sCy5 and diABZI-DMA-Cy5 localized with the ER membrane resulting in colocalization of diABZI-sCy5 and diABZI-DMA-Cy5 with ER Tracker Red in STING-GFP MEFs. An analog containing a cleavable disulfide linker, diABZI-SS-DMA-Cy5, was used as a control. Due to the reducing environment of the cytosol, diABZI is expected to be released from DMA-Cy5, resulting in reduced localization with the ER membrane. **(G)** Representative confocal image of STING-GFP MEFs treated with diABZI-DMA-Cy5 for 2 hours and stained for ER Red and DAPI (nuclear stain); representative images of cells treated with diABZI-sCy5 and diABZI-SS-DMA-Cy5 are shown in **Figure S7-8**. **(H)** Quantification of colocalization between Cy5-labeled diABZI variants and ER Red stain as determined using a Manders coefficient (n=7) calculated by JACoP ImageJ Plugin. All data were plotted using GraphPad 10 with multiple comparisons through one-way ANOVA. *P < 0.05, **P < 0.01, ***P < 0.001, ****P < 0.0001; one-way ANOVA with Tukey’s multiple comparisons test.

While amide bonds on polymer carriers are considered to be biologically stable,^33^ ^34^ particularly in the 1-4 hour time frame where diABZI-DMA activates STING, we evaluated the stability of diABZI-DMA in Tritosomes, lysosomes isolated from rat liver that are commonly used to investigate drug and conjugate stability and catabolism. We used MALDI mass spectroscopy to assess the emergence of diABZI-amine, which would be the predicted product if the amide bond linking diABZI to the polymer backbone was cleaved by lysosomal proteases. Even after incubation of diABZI-DMA with Tritosomes for 7 hours, we did not observe the formation of peaks that would correspond to liberated diABZI; however, diABZI-amine remains detectable using MALDI from 0-7 hours of incubation with Tritosomes at a matched molar concentration (**Figure 4E, Figure S6**). These data suggest that diABZI-DMA remains stable and is not rapidly degraded within endolysosomes, and, therefore, STING activation can be induced by diABZI-DMA as an intact macromolecule rather than by a released small molecule diABZI biproduct that can diffuse across biological membranes to activate STING.

STING primarily resides on the ER membrane where binding to diABZI or other agonists triggers its translocation to the Golgi apparatus, where TBK1 is recruited to phosphorylate STING and IRF3, and is subsequently trafficked on vesicles to late endosomes and lysosomes where STING is ultimately degraded. However, there is also evidence that STING does not remain stationary on the ER and instead moves continuously back and forth between the ER and the Golgi and may even require translocation to late endosomal membranes to activate IRF3.^31^ ^35^ While mechanisms of STING trafficking and signaling are not completely understood, given the complex dependence on vesicular transport processes it is conceivable that endocytosed STING agonists could ultimately reach the ER. Indeed, in STING-GFP MEF cells, treatment with diABZI-DMA results in the formation puncta spatially distributed near the nucleus, consistent with multimerization of STING on the ER membrane (**Figure 3D**). To further investigate this, we quantified the degree of colocalization of diABZI-sCy5 and diABZI-DMA-Cy5 with the ER in STING-GFP MEF cells using an ER Tracker Red stain. As a control, we synthesized another diABZI-functionalized chain transfer agent, diABZI-SS-ECT **(21)**, that includes a disulfide linker that would result in release of the Cy5-labeled polymer from diABZI under the reductive conditions of the cytosol. Using diABZI-SS-ECT as the CTA, we synthesized a diABZI-SS-DMA **(23)** analog that we also labeled with Cy5 (diABZI-SS-DMA-Cy5). The synthesis and chemical characterization of diABZI-SS-ECT and diABZI-SS-DMA-Cy5 is shown in **Schemes S7-8** and **Figures S23-24,S30,S37, and S39**. Therefore, if diABZI targets the Cy5-labeled DMA chain to the ER, decreased ER colocalization would be expected for the disulfide linked variant since bond reduction would liberate DMA-Cy5 from the ER membrane and into the cytosol (**Figure 4F**). As anticipated based on its more potent activity, we observed that diABZI-sCy5 colocalized with the ER to a greater degree than diABZI-DMA-Cy5. Nonetheless, diABZI-DMA-Cy5 colocalized with the ER to a significant degree that was higher than the diABZI-SS-DMA-Cy5 control (**Figure 4G-H, Figure S7-8**). Taken together, these findings suggest that at least a subset of endocytosed diABZI-DMA can access, bind to, and activate STING on the ER as an intact macromolecule. While the mechanisms through which this occurs are likely dynamic and complex, this suggests that vesicular transport of endocytosed macromolecular diABZI compounds may be harnessed for STING activation. This finding also opens the intriguing possibility that diABZI may be employed as an ER-targeting moiety with potential to localize covalently associated drug cargo to the ER membrane ^36^ and potentially synergize with STING activation; for example, ER-targeting of chemotherapeutics has been shown to increase tumor immunogenicity and targeting of antigens to the ER can improve responses to vaccines.^37^ ^38^ This also challenges the assumption that STING agonists must access the cytosol – either directly via passive membrane diffusion or escape from endosomal confinement – and may partially explain how delivery of poorly membrane permeable CDNs in liposomes or other carriers that do not promote cytosolic delivery are effective in enhancing CDN delivery and activity. ^39^ ^40^ ^41^

However, our studies cannot rule out other potential mechanisms of by which macromolecular diABZI constructs may activate STING and potentially in parallel. As noted above, STING is a unique pattern recognition receptor whose activation is partially regulated by its intracellular localization and, in addition to localization on the ER membrane, STING is found in various vesicular compartments such ERGIC, the Golgi, and endolysosomes,^31^ providing ample opportunity for endocytosed diABZI conjugates to encounter and bind STING and/or to harness endogenous vesicular transport mechanisms to reach the ER. Our findings set the stage for such future investigations to more fully dissect these possible novel mechanisms of STING activation, findings that will have important implications for the design of diABZI-based prodrugs and drug carrier platforms.

### Macromolecular diABZI conjugates are therapeutically active in mouse tumor models

Since STING agonists are being actively explored as cancer immunotherapeutics and diABZI compounds have entered clinical trials (NCT03843359), we next sought to validate the activity of mPEG-diABZI and diABZ-DMA constructs in mouse tumor models. First, we compared the efficacy of diABZ-PEG[20kDa], diABZI-V/C-PEG[20kDa], and diABZI-amine in a B16.F10 melanoma murine tumor model, a poorly immunogenic model commonly employed in preclinical immunotherapy development. After establishing ∼50 mm^3^ subcutaneous B16.F10 tumors, mice were intravenously injected with either PBS (vehicle), diABZI-amine, diABZI-20kPEG, or diABZI-V/C-20kPEG every 3 days for a total of 3 treatments at 0.035 μmol diABZI/mouse. Tumor volume was measured until day 18 when the first tumor reached 1500 mm^3^. Consistent with *in vitro* activity studies, there were no significant differences between treatments and all inhibited tumor growth to a comparable degree **(Figure 5A-C)**.

**Figure 5:**
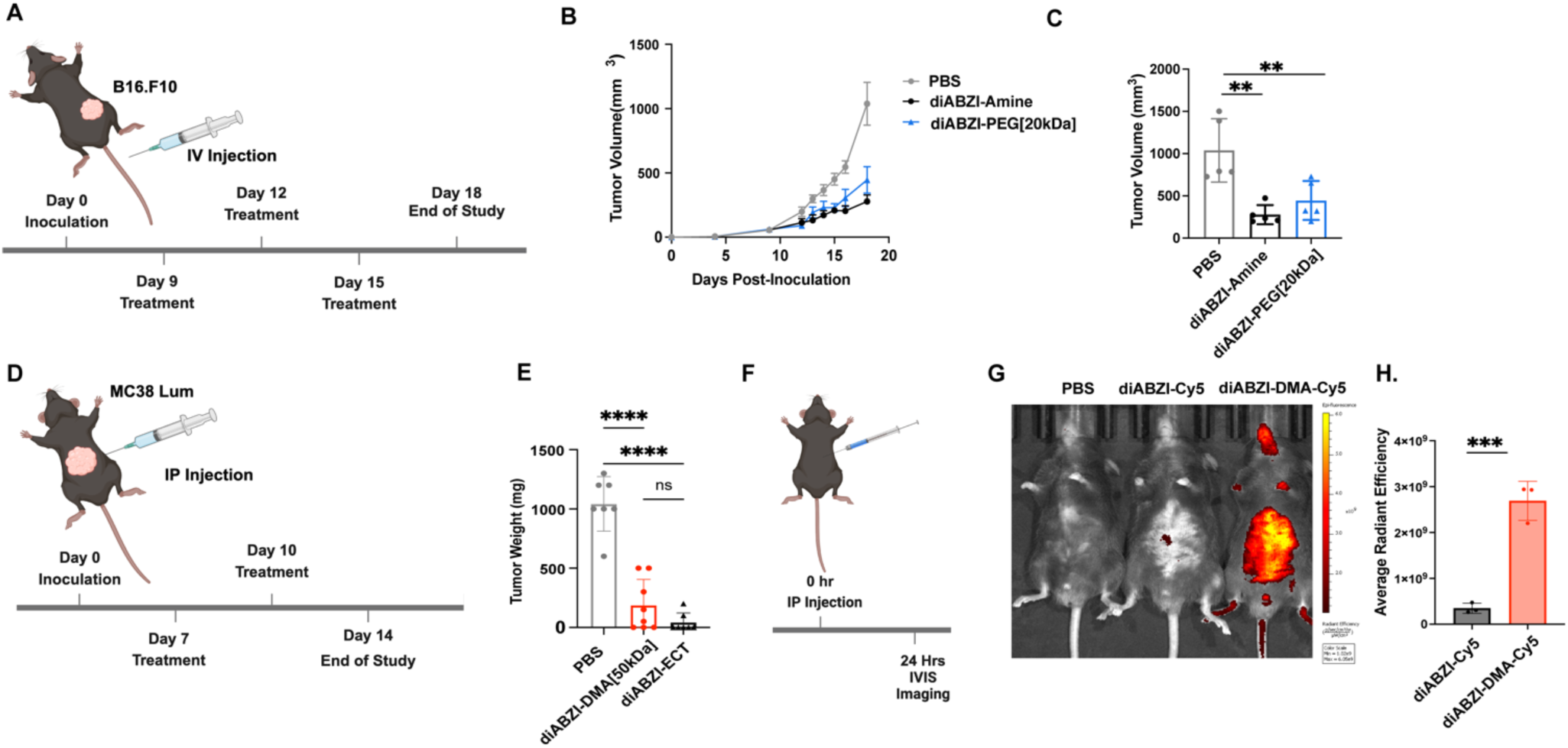
In vivo activity of macromolecular diABZI conjugates in tumor models. **(A)** Schematic of subcutaneous B16.F10 tumor inoculation, treatment schedule, and study endpoint for evaluation of therapeutic activity of diABZI-Amine, diABZI-PEG[20kDa], or diABZI-V/C-PEG[20kDa] relative to vehicle (PBS) treatment. **(B)** Average tumor volume growth curves for mice with B16.F10 melanoma tumors treated as indicated. **(C)** B16.F10 melanoma tumor volumes on day 18 (n=5). **(D)** Schematic of intraperitoneal MC38-Luc tumor inoculation, treatment schedule, and study endpoint for evaluation of therapeutic activity of diABZI-DMA and diABZI-ECT relative to vehicle (PBS) treatment. **(E)** Total weight of MC38-Luc tumors retrieved from peritoneal cavity on day 14 post-inoculation (n=7-8). **(F)** Schematic and study timeline for intraperitoneal administration of diABZI-sCy5 and diABZI-DMA-Cy5. **(G)** Representative IVIS images of mice 24 hours post-injection of either diABZI-sCy5 or diABZI-DMA-Cy5. **(H)** Quantitative analysis of Cy5 fluorescence 24 hours post-injection of diABZI-sCy5 and diABZI-DMA-Cy5 (n=3). All data were plotted using GraphPad 10 with multiple comparisons through one-way ANOVA. *P < 0.05, **P < 0.01, ***P < 0.001, ****P < 0.0001; one-way ANOVA with Tukey’s multiple comparisons test.

We evaluated the efficacy of diABZI-DMA in a model of peritoneal metastasis in which luciferase-expressing MC38 colon cancer cells are injected intraperitoneally to seed tumor nodules throughout the peritoneal cavity. Mice were administered diABZI-ECT or diABZI-DMA (0.007 μmol diABZI) intraperitoneally on days 7 and 10 following tumor inoculation and tumor burden was evaluated by quantification of excised tumor mass on day 14 **(Figure D-E)**. Both diABZI-ECT and diABZI-DMA were similarly effective in inhibiting tumor growth with diABZI-ECT exhibiting greater, but statistically insignificant, antitumor effects. Hence, these studies demonstrate that stable macromolecular diABZI-PEG and diABZI-DMA conjugates can activate STING both *in vitro* and *in vivo*.

The similar degree of efficacy observed between diABZI-ECT and diABZI-DMA may also reflect an interplay between potency and pharmacological properties that impact the magnitude and kinetics of STING activation. Notably, diABZI-ECT is more potent than diABZI-DMA (**Figure 3A**) owing to its high membrane permeability while the higher molecular weight of diABZI-DMA may allow for increased half-life, modulated clearance rates and mechanisms, and/or cellular and tissue distribution profiles; for example, we found that intraperitoneally administered diABZI-DMA-Cy5 had a longer intraperitoneal residence time than diABZI-sCy5 (**Figure F-H**). The objective of these studies was only to validate the *in vivo* activity of stable macromolecular diABZI agonists rather than to engineer constructs for maximal antitumor efficacy or to investigate the impact of STING activation on antitumor immunity or the tumor microenvironment, which we and others have described previously across a range of tumor types.^17^ ^42^ ^43^ ^44^ However, these findings motivate future work focused on leveraging both stable and cleavable conjugates to optimally balance diABZI potency and pharmacological properties for specific disease applications.

## Conclusion

Dimeric amidobenzimidazole (diABZI) STING agonists hold promise as cancer immunotherapeutics, vaccine adjuvants, and antiviral agents, motivating the recent exploration of diABZI-polymer conjugates to modulate their pharmacological properties to improve efficacy and/or safety. Central to this effort is the design and synthesis of diABZI analogs with reactive handles for covalent ligation to carriers and drug linkers to modulate drug release, stability, and/or bioavailability. Here report the surprising finding diABZI compounds linked to multiple highly-water soluble and large (up to 175 kDa) polymers via a stable amide bond can rapidly bind and activate STING both *in vitro* and *in vivo*. We demonstrate that while such macromolecular diABZI constructs enter cells through endocytosis they can localize to the ER to bind STING and that this does not appear to be dependent on drug release and subsequent delivery to the cytosol. While the biological mechanisms through which this occurs remain to be elucidated, our studies support an alternative pathway whereby intracellular trafficking of diABZI-polymer conjugates may be harnessed to access and activate STING. This finding has potentially important implications for the continued design of polymer-drug conjugates, linkers, and prodrugs for STING activation and serve to motivate the exploration of diABZI-carrier conjugates with stable linkers for certain therapeutic applications.

## Methods

### Molecule Synthesis (additional details provided in Supplemental Information)

We synthesized diABZI-amine with a reactive primary amine handle, enabling it to be tethered to the macromolecule through stable bonds that do not release small molecules. To achieve the desired macromolecules, we employed two distinct strategies. In the first strategy, we synthesized stable analogs of diABZI-amine by reacting it with activated N-hydroxysuccinimide (NHS) esters of MeO-PEGn-NHS, producing two analogs: diABZI-PEG[5KDa] (2) and diABZI-PEG[20KDa] (3). The primary amine group of diABZI-amine attacks the carbonyl group of the activated ester, forming an addition product with a tetrahedral intermediate. Subsequently, the more labile leaving group is released, resulting in the formation of a stable secondary amide through nucleophilic substitution. In the second strategy, we reacted diABZI-amine with the activated PNP ester of a chain transfer agent (10), forming a diABZI-based chain transfer agent analog (11). This analog was used to synthesize the macromolecules 12-14 via RAFT polymerization in a controlled manner.

### Cathepsin B Activity Assay

Recombinant Mouse Cathepsin B (RND Systems) was prepared at 50 µM in MES buffer (pH 5.0) upon opening and kept in −80 °C conditions when not in use. To activate the enzyme, Cathepsin B was diluted to 0.2 µM in MES buffer (pH 5.0) containing 1 mM EDTA and 2 mM DTT and placed at 37 °C for 15 minutes. After activation, Cathepsin B was combined with 50 µM of substrate at 37 °C for 48 hours at a total volume of 100 µL. Activity was determined by observing molecular weight shifts in the substrate using matrix assisted laser desorption and ionization mass spectrometry (MALDI-MS).

### MALDI-MS

4 µL of Matrix (20 mg mL^-1^ THAP and 20 mg mL^-1^ CHCA in dry acetone) was combined with 1 µL of sample from the Cathepsin B activity assay and spotted on a stainless steel MALDI-MS plate (Bruker). After evaporation of matrix, three technical replicates were collected for each spot using FlexControl software (Bruker Daltonics) on a Bruker AutoFlex MALDI-TOF. The laser pulse rate was 1000 Hz and spectra were obtained with a mass window of 600-5000 *m/z* at high resolution (4.00 GS/s). FlexAnalysis software (Bruker Daltonics) was used to perform unbiased smoothing and obtain baseline spectra for all samples. Spectra were exported to Microsoft Excel and spectra were plotted using MATLAB or GraphPad Prism.

### THP1-Dual Reporter Cell Assay

THP1-Dual cells and STING-Knockout THP1 Duals (InvivoGen) were cultured in Roswell Park Memorial Institute (RPMI) 1640 Medium (Gibco) supplemented with 2 mM L-glutamine, 25 mM HEPES, 10% heat-inactivated fetal bovine serum (HI-FBS; Gibco), 100 U ml^−1^ penicillin/100 μg ml^−1^ streptomycin (Gibco), and 100 µg/mL Normocin. Cells were subjected to 10 µg/mL Blasticidin and 100 µg/mL Zeocin for continual selection after every cell passage. 96-well plates (REF 655180; Greiner Bio-One) were used for screening agonist activity. Reporter cells were seeded at 25,000 cells/well in 100 µL media and treatments were administered in 100 µL of medium. Results were collected 24 hours after treatment using a Quanti-Luc™ or Quanti-Blue™ (InvivoGen) assay on cell supernatants following manufacturer’s instructions. Luminescence was quantified using a plate reader (Synergy H1 Multi-Mode Microplate Reader; Biotek) after supernatants were transferred to opaque-bottom 96-well plates (REF 655073; Greiner Bio-One).

### Splenocyte Isolation and IFN-β Reporter ELISA

Spleens were harvested from Female C57BL/6 mice (8 weeks old), mechanically disrupted into single-cell suspensions through a 70 μm cell strainer (FisherbrandTM; Thermo Fisher Scientific), and suspended in complete RPMI 1640 medium (Gibco) supplemented with 10% FBS, 10% HI-FBS (Gibco), 100 U ml^−1^ penicillin/100 μg ml^−1^ streptomycin (Gibco), 50 μM 2-mercaptoethanol, and 2 mM L-glutamine. The cells were centrifuged for 5 min at 1500 rpm and resuspended in ACK lysis buffer (KD Medical) for 5 minutes. Cells were centrifuged and resuspended in fresh media at a concentration of 3 million cells per mL. Cells were seeded in a 96 well round bottom plate at 100 μL per well and treatments were administered in 100 μL of medium. Results were collected 24 hours after treatment using a mouse IFN-β solid-phase sandwich ELISA kit (Invivogen Cat#42400-1) on cell supernatants following manufacturer’s instructions. Luminescence was quantified using a plate reader (Synergy H1 Multi-Mode Microplate Reader; Biotek).

### Isothermal Calorimetry (ITC)

Recombinant hSTING was synthesized in Escherichia coli (New England Biolabs, T7 SHuffle Express line) and purified by affinity chromatography. Buffer exchange was performed prior to ITC (pH 7.5: PBST, 150 mM NaCl, 3 mM EDTA, 0.05% Tween 20), using Amicon Ultra 4 mL centrifugal filters (Millipore, Etobicoke, Canada). ITC experiments were performed on a TA Instruments Affinity ITC instrument. 24 total injections were performed using the following instrument settings: cell temperature 25°C, reference power 10 µCal/second, initial delay 240 seconds, stirring speed 75 rpm, feedback mode/gain high, and injection volume 2 µL for 10 seconds spaced at 120 second intervals with a filter period of 10 seconds. hSTING was set in the cell at a concentration of 10 µM and a volume of 350 µL. Agonists were prepared at a stock concentration of 20 mM in DMSO and diluted using pH 7.5 PBST to 150 µM for titration by the syringe (120 µL). Commerical diABZI molecule compound 3 was purchased through Selleck Chem (Catalog No.S8796). Data were analyzed using TA Instruments NanoAnalyze Software.

### RAFT Polymerization of diABZI-DMA Polymers (12-14)

Reversible addition-fragmentation chain transfer (RAFT) was used to synthesize analogs of polymers with distinct molar masses. For synthesis dimethyl acrylamide (DMA) monomer was filtered over activated alumina and allowed to react under inert atmosphere in DMF (30 wt% monomer) at 70 °C for 24 h in an oil bath. The initial monomer ([M]_o_) to diABZI-CTA compound **5** ([CTA]_o_) to initiator ([I]_o_) ratio was n:1:0.2. The resultant desired polymers DMA(nk)-diABZI-CTA was isolated by dialysis against pure acetone (2×), pure deionized water (2×). Following dialysis, the purified compound was frozen at −80 °C for 8 h and then lyophilized for 3 days. Final polymers were characterized by 1H NMR and GPC (TOSOH) in HPLC-grade DMF with 0.01 M LiBr via the LenS3 Multi-Angle Light Scattering Detector (TOSOH).

### THP1-Dual qRT-PCR

THP1-Dual cells (InvivoGen) were cultured in Roswell Park Memorial Institute (RPMI) 1640 Medium (Gibco) supplemented with 2 mM L-glutamine, 25 mM HEPES, 10% heat-inactivated fetal bovine serum (HI-FBS; Gibco), 100 U ml^−1^ penicillin/100 μg ml^−1^ streptomycin (Gibco), and 100 µg/mL Normocin. Cells were subjected to 10 µg/mL Blasticidin and 100 µg/mL Zeocin for continual selection after every cell passage. 96-well plates (REF 655180; Greiner Bio-One) were used for screening agonist activity. Reporter cells were seeded at 300,000 cells/well and treated with 5 μM drug. Cell pellets and RNA was extracted via Qiagen RNeasy Plus Mini Kit. An iScript cDNA synthesis kit (Bio-Rad) was used to synthesize cDNA per manufacturer’s protocol. RT-qPCR on the cDNA was run using TaqMan gene expression kits (primer and master mix) and run on the Bio-Rad CFX Connect Real-time System, with a threshold cycle number determination made by the Bio-Rad CFX manager software V.3.0. Primers: mouse Ifnb1 (Mm00439552_s1), mouse Tnf (Mm00443258_m1), mouse Cxcl10 (Mm00445235_m1),, mouse IL-6 (mouse IL-6) and mouse Hmbs (Mm01143545_m1). Gene expression relative to Hmbs was calculated using 2^-(Cq-CqHmbs)^.

### Western Blot Analysis

Cells were lysed with 50 μl RIPA buffer (Sigma-Aldrich) supplemented with protease inhibitors (Sigma-Aldrich). Protein concentration was measured using a BCA protein assay kit (Thermo Scientific). Equal amount of protein (10 −30μg) was loaded to SDS/PAGE and transferred onto nitrocellulose membranes using the semi-dry transfer protocol (Bio-Rad Laboratories). After transfer, membranes were probed with each respective primary antibody (anti-Sting, anti-p-Sting, anti-TBK1, anti-p-TBK1, anti-IRF3, anti-p-IRF3, and anti-Hsp-90) overnight at 4°C. All antibodies were purchased from Cell signaling. Following incubation, the membranes were probed with HRP-conjugated secondary antibodies. All antibodies were purchased from Cell signaling. Protein bands were visualized using an ECL western blotting substrate (Thermo Scientific). Images of immunoblots were obtained using a LI-COR Odyssey Imaging System.

### Confocal Microscopy for STING Kinetics

STING-GFP Murine Fibroblasts (provided by N. Yan, UT Southwestern) were grown in complete DMEM medium supplemented with fetal bovine serum (HI-FBS; Gibco), and 100 U ml^−1^ penicillin/100 μg ml^−1^ streptomycin (Gibco). These were seeded at 1×10^5^ cells per slip on fibronectin-coated coverslips and incubated overnight. Cells were washed with PBS and dosed with 5 μM drug in complete DMEM (no phenol red, Gibco). Confocal Images were taken over time using a Nikon A1R-HD25 confocal microscope with motorized stage, Tokai Hit stage top incubation system with built-in digital gas mixer for 100% CO2 use, equipped with Apo TIRF 60x/1.49 oil immersion lens. check it). Images were acquired using the Nikon NIS-Elements software and analyzed through ImageJ.

### RAFT polymerization for diABZI-DMA-AzPMAm and diABZI-DMA-AzPMAm (17 and 22)

For synthesis, DMA was filtered over activated alumina, combined with a solution of diABZI-ECT, AIBN, and AzPMAm, and allowed to react under inert atmosphere in DMF (30 wt% monomer) at 70 °C for 24 h in an oil bath. The initial DMA to AzPMAm to diABZI-ECT to AIBN ratio was 496:11:1:0.2 for (17) and 490:15:1:0.2 (22). The resultant desired polymer, was isolated by dialysis against pure acetone (2×), 1:1 pure acetone and deionized water (1x), and then pure deionized water (2×). Following dialysis, the purified compounds wwere frozen at −80 °C for 8 h and then lyophilized for 72 hours, which was further characterized by 1H NMR and GPC (TOSOH) in HPLC-grade DMF with 0.01 M LiBr via the LenS3 Multi-Angle Light Scattering Detector (TOSOH).

### Flow Cytometry for Cell Uptake

THP1-Duals were plated in FBS containing RPMI media at 100,000 cells per well in a 96 well plate at 180 µL and incubated at either 4 °C diluted to 5 µM in FBS containing RPMI media for a final volume of 200 µL and incubated at either 4 °C or 37 °C for 2 hours. After incubation, the plates were centrifuged at 500 x g for 5 minutes at 4 °C, decanted, and washed three times with refrigerated 1% BSA containing PBS. A 20,000 x 1 dilution of DAPI was used to prepare a DAPI containing 1% BSA in PBS solution for a final cell suspension, at 100 µL per well. Data were collected and analyzed for cell uptake on a CellStream Flow Cytometer (Luminex) equipped with SSC, FFC, 405 (DAPI), and 642 (Cy5) nm lasers.

### Confocal Microscopy for Intracellular Colocalization

STING-GFP Murine Fibroblasts (provided by N. Yan, UT Southwestern) were grown in complete DMEM medium supplemented with fetal bovine serum (HI-FBS; Gibco), and 100 U ml^−1^ penicillin/100 μg ml^−1^ streptomycin (Gibco). These were seeded at 1×10^5^ cells per slip on fibronectin-coated coverslips and incubated overnight. Cells were washed with PBS and dosed with 5 μM drug in complete DMEM (no phenol red, Gibco) for 2 or 4 hours. For lysotracker assays, 75nM Lysotracker Red DND-99 (Thermofisher) was added 1 hour before imaging and cells were washed with complete DMEM (no phenol red, Gibco) immediately before imaging. For ER colocalization assays, 1μM ER Tracker Red (Thermofisher) was added 15 minutes before imaging and cells were washed and incubated with complete DMEM (no phenol red, Gibco) containing 1:50,000 DAPI nuclear stain immediately before imaging. Confocal Images were taken over time using a Nikon A1R-HD25 confocal microscope with motorized stage, Tokai Hit stage top incubation system with built-in digital gas mixer for 100% CO2 use, equipped with Apo TIRF 60x/1.49 oil immersion lens. check it). Images were acquired using the Nikon NIS-Elements software and analyzed through ImageJ. Manders coefficients were calculated using JACoP ImageJ plugin.

### Tritosome Assay

Isolated rat lysosomes (tritosomes) and lysosome/tritosome buffer were purchased from BioIVT. diABZI variants were incubated at 100 μM in sterile water complete with 10% tritosome and 10% buffer for 7 hours at 37°C in a total of 50μL reaction volume. 20μL samples were taken at 0 and 7 hours and frozen a −80°C until MALDI-MS analysis was performed as described above. Data were plotted via GraphPad Prism.

### B16 Melanoma Tumor Model

6-8 week-old C57BL/6 mice were inoculated with B16.F10 tumors by subcutaneously injecting 1 x 10^6^ cells suspended in 100 µL of PBS into the rear right flank. When tumors were ∼ 50 mm^3^, the mice were given a total of 3, 100 µL intravenous injections administered every 3 days containing either PBS, diABZI-Amine, diABZI-PEG20k at 0.035 μmol/mouse. Tumor volume and murine weight were measured for the duration of the study. The study endpoint for maximum tumor volume was 1500 mm^3^.

### MC38-Luc Tumor Model

6-8 week-old C57BL/6 mice were inoculated with MC38 Lum tumors by injecting 1 x 10^6^ cells suspended in 100 µL of PBS into the IP space. Tumors were qualitatively measured using IVIS Imaging and randomized on day 6 post inoculation. Mice were treated on day 7 and day 10 post-inoculation with either PBS, diABZI-ECT, of diABZI-DMA[50kDa] at 0.007 μmol/mouse. Mice were sacrificed on day 14 post-inoculation and tumor weights were reported.

### IP Retention Study

7 week-old C57BL/6 mice were injected (IP) with PBS, diABZI-sCy5, or diABZI-DMA-Cy5 (0.006 μmol/mouse) at 100μL injection volume and imaged under isoflurane 24 hours post-injection using an IVIS Lumina III (PerkinElmer). Fluorescence (radiant efficiency) was measured and average radiant efficiency values (per cm2) were calculated using the Living Image software (version 4.5).

### Statistical analysis

Significance for each experiment was determined as indicated in the corresponding figure captions. All analyses were done using GraphPad Prism software, version 9. Plotted values represent experimental means, and error bars represent SD unless otherwise noted in the figure captions. **** P < 0.0001, *** P < 0.005, **P < 0.01, * P < 0.05.

## Acknowledgements

The authors thank C. Duvall (Vanderbilt University) for use of IVIS Imaging System, N. Yan (UT Southwestern) for providing MEF STING-GFP cells, and the Vanderbilt University Small Molecular NMR Facility. This research was supported by grants from Susan G. Komen (CCR19609205 to J.T.W.), the National Institutes of Health (R01 CA245134, R01 CA266767, and R01 CA274675 to J.T.W.), a Vanderbilt Ingram Cancer Center (VICC) Ambassador Discovery Grant (J.T.W.), VICC Support Grant P30 CA68485, and funds provided by the Vanderbilt University School of Engineering (J.T.W.). TLS acknowledges funding support through the National Science Foundation Graduate Research Fellowship Program under grant number 193793. BRK acknowledges postdoctoral funding support from the PhRMA Foundation Postdoctoral Fellowship in Drug Delivery. Schematics were made using Biorender.com.

## Notes

### Competing Interest Statement

J.T.W, K.A., T.L.S., and J.A.S are co-inventors on a patent application (PCT/US2023/076732) on polymer-drug conjugates for STING pathway activation, including the compounds described in this work.

