## Supporting Information for "Macromolecular Diamidobenzimidazole Conjugates Activate STING"

**(9 Pages)**

**Part A.** General Considerations

**Part B.** Synthesis — Scheme S1-S8

**Part C.** Supplemental Data — Figures S1-S8

**Part E.** [1]H NMR spectra for new compounds. — Figures S9-S24

**Part F.** [13]C NMR spectra for new compounds. — Figures S25-S30

**Part G.** HRMS spectra for new compounds. — Figures S31-S37

**Part H.** GPC Analysis for polymers. — Figures S31-S37

**Part I.** References

**Part A: General Considerations**

All reagents were purchased from commercial suppliers and used as received. NMR spectra were recorded on a Bruker – 400 MHz or 600 MHz Spectrometer. Mass spectra were recorded on a Thermo-Fisher Scientific LTQ-Orbitrap XL Mass Spectrometer. All reactions were performed under ambient atmosphere unless otherwise noted. Anaerobic reactions were performed by purging the reaction solutions with argon or nitrogen. $PEG_{5KDa}$-NHS ester, $PEG_{20Kda}$-NHS ester, Methoxy-$PEG_{5Kda}$-SH and Methoxy-$PEG_{20Kda}$-SH were purchased from JenKem Technology USA. Compounds **1**,[1] **4**,[2] **8**,[3] **10**,[4] **15**,[5] **19**,[6] were synthesized as per the literature protocol.

**Part B: Synthesis**

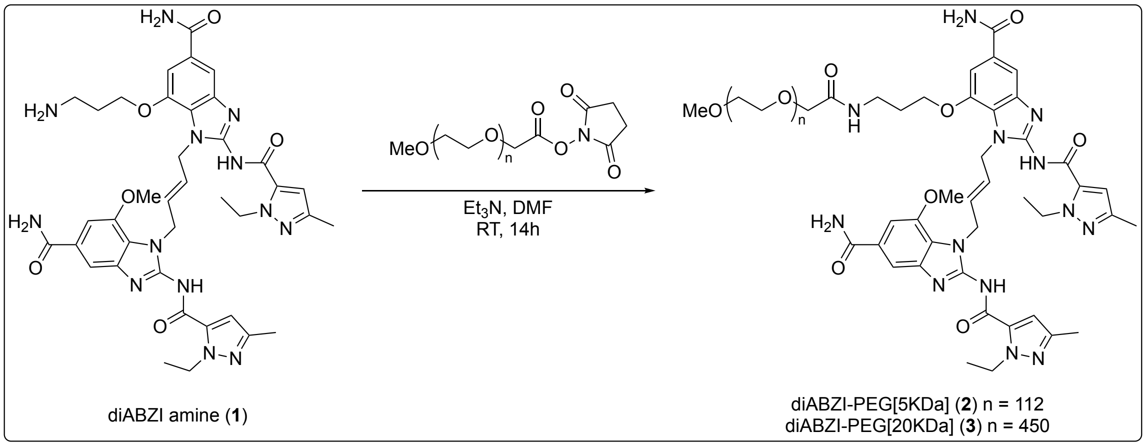


**Scheme S1:** Synthesis of diABZI-PEG[5Kda, 20Kda]

**diABZI-PEG$_{nKDa}$ (2,3).** To a stirred solution of diABZI amine (**1**) (1.2 eq) and $Et_3N$ (5 eq) in DMF add solution of Methoxy-PEG(nk)-NHS ester (1.0 eq) in DMF:DCM. The reaction mixture was stirred overnight and purified by dialysis (3kDa MWCO) against (1:1) DCM:MeOH (2 ×), Acetone (2 ×) followed by distilled water (2 ×). Following dialysis, the purified compound was frozen at -80 ºC for 8 h and then lyophilized for 3 days to obtain the desired purified product and characterized by $^1$H NMR.

**diABZI-PEG[5KDa] (2)**: To a stirred solution of diABZI amine (**1**) (81 mg, 72 µmol, 1.2 eq) and $Et_3N$ (42 µL, 300 µmol, 5 eq) in DMF (2 mL) add solution of Methoxy-PEG(5k)-NHS ester (300 mg, 60 µmol, 1 eq) in 1:1 DCM:DMF 4 mL and stir overnight. The desired compound **6** was purified by dialysis (3kDa MWCO) against (1:1) DCM:MeOH (2 ×), acetone (2 ×) followed by distilled water (2 ×). The desired purified product (180 mg, 31.2 µmol, 52%) was lyophilized and characterized by $^1$H NMR. $^1$H NMR (400 MHz, DMSO) δ 7.95 (broad s, 2H), 7.94 (broad s, 3H), 7.73 (t, 1H), 7.64 (s, 2H), 7.31 – 7.28 (m, 5H), 6.5 (d, 2H), 5.87 – 5.82 (m, 2H), 4.96 – 4.89 (m, 4H), 4.53 – 4.50 (m, 4H), 3.97 (t, $J$ = 6.1 Hz, 2H), 3.81 (s, 3H, terminal methoxy of PEG), 3.72 (s, 3H), 3.51 (broad s, 462H, -$CH_2$-$CH_2$- of PEGMA), 2.10 (s, 3H), 2.09 (s, 3H), 1.70 (p, $J$ = 6.1 Hz, 2H), 1.28 – 1.24 (m, 6H).

**diABZI-PEG[20KDa] (3)**: To a stirred solution of diABZI amine (**1**) (34 mg, 30 µmol, 1.2 eq) and $Et_3N$ (17.5 µL, 125 µmol, 5 eq) in DMF (1.5 mL) add solution of Methoxy-PEG(20k)-NHS ester (500 mg, 25 µmol, 1 eq) in 1:1 DCM:DMF 5 mL and stir overnight. The desired compound **7** was purified by dialysis (3kDa MWCO) against (1:1) DCM:MeOH (2 ×), acetone (2 ×) followed by distilled water (2 ×). The desired purified product was lyophilized to obtain white fluffy powder (393 mg, 19 µmol, 76%).

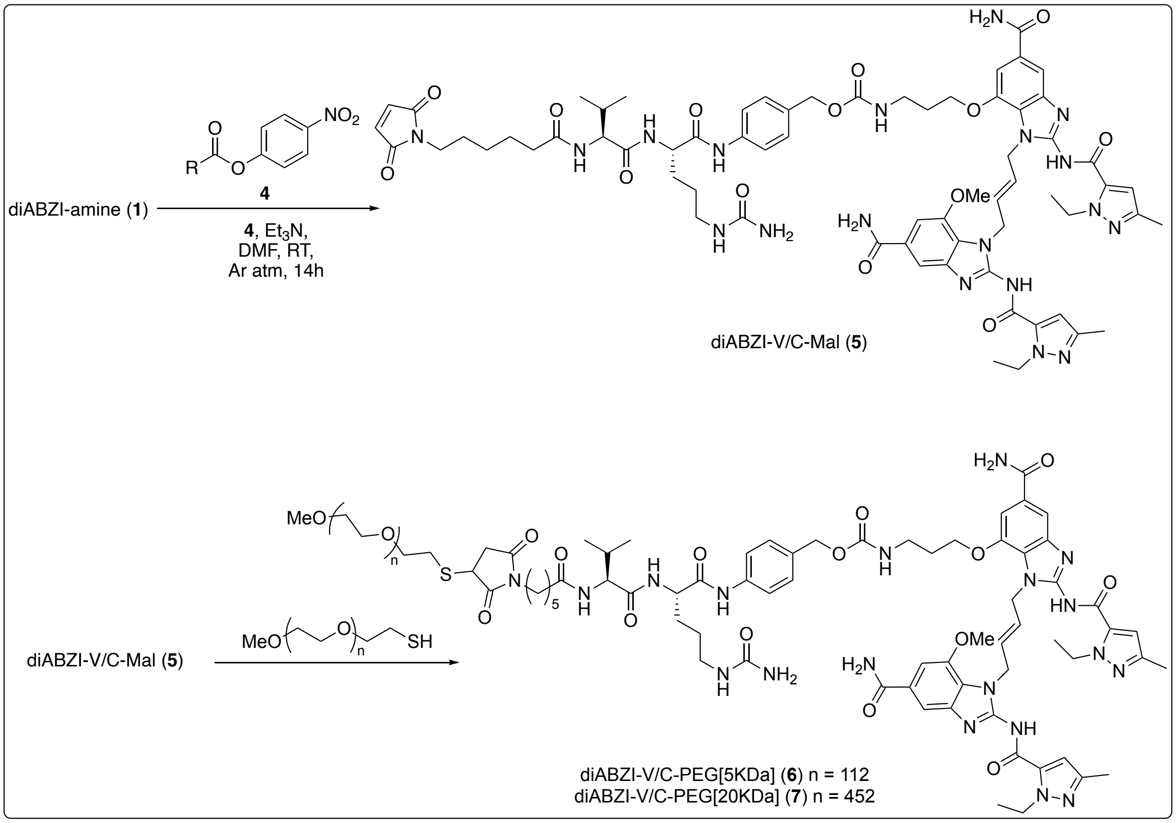


**Scheme S2:** Synthesis of diABZI-V/C-Mal (**5**), diABZI-V/C-PEG[5KDa, 20KDa] (**6**, **7**)

**4-((*S*)-2-((*S*)-2-(6-(2,5-dioxo-2,5-dihydro-1*H*-pyrrol-1-yl)hexanamido)-3-methylbutanamido)-5-ureidopentanamido)benzyl (3-((5-carbamoyl-1-((*E*)-4-(5-carbamoyl-2-(1-ethyl-3-methyl-1*H*-pyrazole-5-carboxamido)-7-methoxy-1*H*-benzo[*d*]imidazol-1-yl)but-2-en-1-yl)-2-(1-ethyl-3-methyl-1*H*-pyrazole-5-carboxamido)-1*H*-benzo[*d*]imidazol-7-yl)oxy)propyl)carbamate (5).** To a stirred solution of (*E*)-7-(3-aminopropoxy)-1-(4-(5-carbamoyl-2-(1-ethyl-3-methyl-1*H*-pyrazole-5-carboxamido)-7-methoxy-1*H*-benzo[*d*]imidazol-1-yl)but-2-en-1-yl)-2-(1-ethyl-3-methyl-1*H*-pyrazole-5-carboxamido)-1*H*-benzo[*d*]imidazole-5-carboxamide tris trifluoracetic acid **1** (350 mg, 0.3 mmol, 1eq) and hunig's base (0.27 mL, 1.56 mmol, 5 eq) in DMF (5 mL) dropwise added solution of 4-((*S*)-2-((*S*)-2-(6-(2,5-dioxo-2,5-dihydro-

1*H*-pyrrol-1-yl)hexanamido)-3-methylbutanamido)-5-ureidopentanamido)benzyl (4-nitrophenyl) carbonate **9**[2] (253 mg, 0.34 mmol, 1.1 eq) in DMF (3 mL) under inert atmosphere which turned reaction mixture into a bright yellow color that indicate releasing of para nitrophenol. The reaction mixture stirred overnight at room temperature. The diethyl ether added to precipitate out the crude solid desired product. The solid was filtered over Büchner funnel and resuspended in 1:1 (DCM:MeOH) 5 mL and stirred for 2 hr. The solid was filtered over Büchner funnel to obtain the pure desired product **10** as a light pink solid (413 mg, 0.29 mmol, 96%). $^1$H NMR (400 MHz, DMSO) δ 9.95 (s, 1H), 8.06 (d, *J* = 7.5 Hz, 1H), 7.95 (broad s, 2H), 7.79 (d, *J* = 8.5 Hz, 1H), 7.63 (d, *J* = 6.7 Hz, 2H), 7.56 (d, *J* = 8.5 Hz, 2H), 7.34 – 7.23 (m, 7H), 6.99 (s, 2H), 6.5 (d, *J* = 5.6 Hz, 2H), 5.96 (t, *J* = 5.0 Hz, 1H), 5.88 – 5.79 (m, 2H), 5.40 (s, 2H), 4.93 – 4.87 (m, 6H), 4.53 – 4.49 (m, 4H), 4.38 – 4.35 (m, 1H), 4.19 – 4.16 (m, 1H), 3.98 (t, *J* = 5.0 Hz, 2H), 3.71 (s, 3H), 3.40 – 3.34 (m, 2H), 3.08 – 2.90 (m, 4H), 2.20 – 1.92 (m, 9H), 1.73 – 1.54 (m, 4H), 1.52 – 1.14 (m, 16H), 0.84 (d, *J* = 6.7 Hz, 3H), 0.80 (d, *J* = 6.7 Hz, 3H). $^{13}$C NMR (151 MHz, DMSO) δ 172.7, 171.8, 171.5, 171.0, 168.1, 167.3, 159.4, 156.6, 145.5, 145.3, 140.4, 139.0, 134.9, 132.2, 130.5, 129.1, 119.4, 109.7, 65.5, 58.0, 56.4, 53.6, 46.0, 37.5, 35.4, 30.8, 29.8, 28.2, 27.3, 26.2, 25.4, 19.7, 18.7, 16.6, 13.6. HRMS (ESMS) Calculated for $C_{67}H_{83}N_{19}O_{14}$ $[M+H]^+$: 1378.6440, found 1378.6415.

**Methoxy-PEG$_n$-Mal-Val-Cit-PAMB-diABZI (6, 7).**

**General method for Thia-Michael addition:** To a stirred solution of compound **10** (1.1 eq) and mPEG(nk)-SH (1.0 eq) in DMF was added solution of N-methyl morpholine (NMM, 4.0 eq) in DMF. The reaction mixture was stirred at room temperature for 2 h and purified by dialysis (3kDa MWCO) against (1:1) DCM:MeOH (2 ×), acetone (2 ×) followed by distilled water (2 ×). Following dialysis, the purified compound was frozen at -80 ºC for 8 h and then lyophilized for 3 days to obtain the desired purified product and characterized by $^1$H NMR.

**diABZI-V/C-PEG[5KDa]** (**6**)**:** To a stirred solution of compound **10** (30 mg, 21.7 µmol, 1.1 eq) and mPEG(5k)-SH (100 mg, 19.8 µmol, 1.0 eq) in DMF (2.0 mL) was added solution of N-methyl morpholine (NMM, 9 µL, 79.1 µmol, 4.0 eq) in DMF (0.5 mL). The desired purified product was lyophilized to obtain white fluffy powder (84 mg, 13.2 µmol, 66%). $^1$H NMR (400 MHz, DMSO) δ 9.96 (s, 1H), 8.08 (s, 1H), 7.95 (broad s, 2H), 7.79 (d, *J* = 8.5 Hz, 1H), 7.64 (d, *J* = 6.7 Hz, 2H), 7.57 (d, *J* = 8.5 Hz, 2H), 7.34 – 7.23 (m, 6H), 6.5 (s, 1H), 5.96 (t, *J* = 5.0 Hz, 1H), 5.89 – 5.79 (m,

2H), 5.40 (s, 2H), 4.94 – 4.88 (m, 6H), 4.53 – 4.49 (m, 4H), 4.39 – 4.35 (m, 1H), 4.20 – 4.18 (m, 1H), 3.98 (t, *J* = 5.0 Hz, 2H), 3.71 (s, 3H), 3.50 (broad s, 395H, -$CH_2$-$CH_2$- of PEGMA), 3.19 – 2.79 (m, 6H), 2.20 – 1.92 (m, 9H), 1.73 – 1.54 (m, 4H), 1.52 – 1.14 (m, 14H), 0.85 (d, *J* = 6.7 Hz, 3H), 0.82 (d, *J* = 6.7 Hz, 3H

**diABZI-V/C-PEG[20KDa] (7):** To a stirred solution of compound **10** (30 mg, 21.7 µmol, 1.1 eq) and mPEG(20k)-SH (396 mg, 19.8 µmol, 1.0 eq) in DMF (2.0 mL) was added solution of N-methyl morpholine (NMM, 9 µL, 79.1 µmol, 4.0 eq) in DMF (0.5 mL). The desired purified product was lyophilized to obtain white fluffy powder (271 mg, 12.7 µmol, 64%).

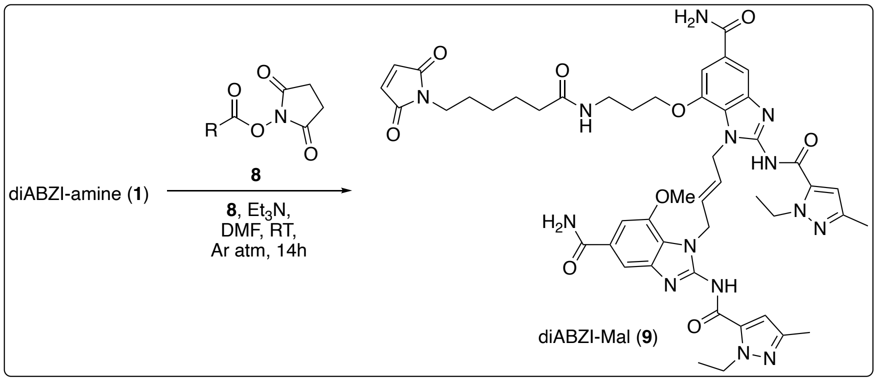


**Scheme S3:** Synthesis of diABZI-Mal (**9**)

**(*E*)-1-(4-(5-carbamoyl-2-(1-ethyl-3-methyl-1*H*-pyrazole-5-carboxamido)-7-methoxy-1*H*-benzo[*d*]imidazol-1-yl)but-2-en-1-yl)-7-(3-(6-(2,5-dioxo-2,5-dihydro-1*H*-pyrrol-1-yl)hexanamido)propoxy)-2-(1-ethyl-3-methyl-1*H*-pyrazole-5-carboxamido)-1*H*-benzo[*d*]imidazole-5-carboxamide (9).** To a stirred solution of (*E*)-7-(3-aminopropoxy)-1-(4-(5-carbamoyl-2-(1-ethyl-3-methyl-1*H*-pyrazole-5-carboxamido)-7-methoxy-1*H*-benzo[*d*]imidazol-1-yl)but-2-en-1-yl)-2-(1-ethyl-3-methyl-1*H*-pyrazole-5-carboxamido)-1*H*-benzo[*d*]imidazole-5-carboxamide tris trifluoracetic acid **1** (200 mg, 0.18 mmol, 1eq) and hunig's base (0.12 mL, 0.89 mmol, 5 eq) in DMF (3 mL) dropwise added solution 2,5-dioxopyrrolidin-1-yl 6-(2,5-dioxo-2,5-dihydro-1*H*-pyrrol-1-yl)hexanoate **8**[3] (66 mg, 0.2 mmol, 1.2 eq) in DMF (2 mL) under inert atmosphere and stirred overnight at room temperature. The diethyl ether added to precipitate out the crude solid desired product. The crude solid was purified over silica gel chromatography (DCM:MeOH 0-20%) to obtain the desired product **9** as an off white solid (80 mg, 0.08 mmol, 46%). $^1$H NMR (400 MHz, DMSO) δ 7.95 (broad s, 2H), 7.76 (t, *J* = 5.5 Hz, 1H), 7.63 (s, 2H),

7.32 (braod s, 2H), 7.29 (d, $J = 6.2$ Hz, 2H), 6.97 (s, 2H), 6.5 (d, $J = 7.6$ Hz, 2H), 5.89 – 5.77 (m, 2H), 4.95 – 4.88 (m, 4H), 4.54 – 4.48 (m, 4H), 3.96 (t, $J = 6.1$ Hz, 2H), 3.71 (s, 3H), 3.09 (dt, $J = 6.2$, 5.9 Hz, 2H), 2.10 (s, 3H), 2.09 (s, 3H), 1.98 (t, $J = 7.2$ Hz, 2H), 1.67 (p, $J = 6.1$ Hz, 2H), 1.47 – 1.40 (m, 4H), 1.25 (dt, J = 7.1, 3.2 Hz, 6H), 1.18 – 1.10 (m, 2H). $^{13}$C NMR (151 MHz, DMSO) δ 172.4, 171.5, 168.1, 167.3, 152.5, 145.5, 145.3, 144.7, 140.4, 140.3, 134.9, 130.5, 130.5, 128.7, 128.2, 120.1, 109.7, 56.4, 46.0, 37.4, 35.6, 29.2, 28.2, 26.3, 25.2, 16.6, 13.6. HRMS (ESMS) Calculated for $C_{48}H_{56}N_{14}O_9$ $[M+H]^+$: 973.4427, found 973.4417.

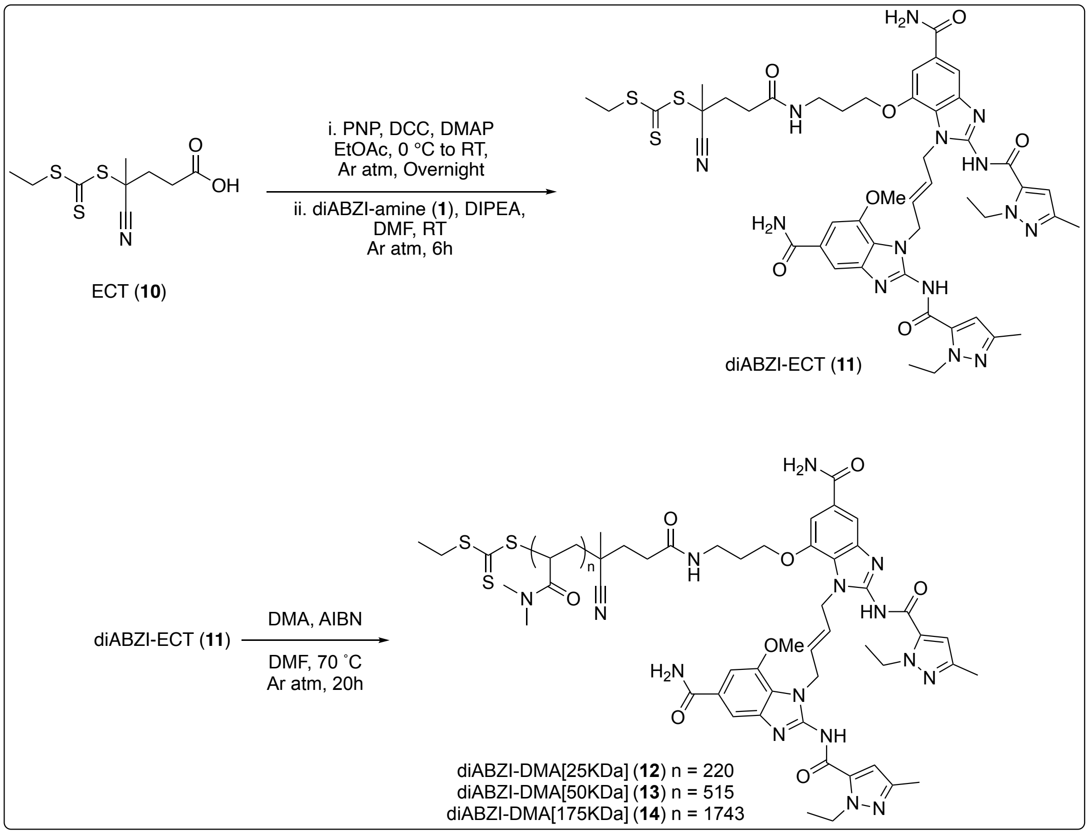


**Scheme S4:** Synthesis of diABZI-ECT, diABZI-DMA[25KDa, 50KDa, 175KDa]

**(*E*)-5-((3-((5-carbamoyl-1-(4-(5-carbamoyl-2-(1-ethyl-3-methyl-1*H*-pyrazole-5-carboxamido)-7-methoxy-1*H*-benzo[*d*]imidazol-1-yl)but-2-en-1-yl)-2-(1-ethyl-3-methyl-1*H*-pyrazole-5-carboxamido)-1*H*-benzo[*d*]imidazol-7-yl)oxy)propyl)amino)-2-cyano-5-oxopentan-2-yl ethyl carbonotrithioate** (**11**). A solution of 4-cyano-4-(((ethylthio)carbonothioyl)thio)pentanoic acid **10**[4] (100 mg, 0.38 mmol, 1 eq), *p*-nitrophenol (53 mg, 0.38 mmol, 1 eq) and DMAP (5 mg, 0.038 mmol, 0.1 eq) in EtOAc (4 mL) was maintained at 0 °C under argon atmosphere. A solution of DCC (82 mg, 0.40 mmol, 1.05 eq) in EtOAc (4 mL) was added dropwise. The reaction mixture was allowed to warm to rt and stirred overnight for 18 h. After consumption of the starting material, as judged by the TLC analyses, the reaction mixture was filtered through celite. The solvent was evaporated *in vacuo* to obtain the activated ester as a crude product. The crude product was analyzed by 1H NMR spectroscopy and was used without further purification. $^{1}$H NMR (400 MHz, Chloroform-*d*) δ 8.28 (d, *J* = 9.2 Hz, 2H), 7.31 (d, *J* = 9.2 Hz, 2H), 3.36 (q, *J* = 7.4 Hz, 2H), 2.95 (t, *J* = 7.9 Hz, 2H), 2.70 – 2.46 (m, 2H), 1.95 (s, 3H), 1.37 (t, *J* = 7.4 Hz, 3H).

A solution of diABZI amine (**1**) (300 mg, 0.27 mmol, 1 eq) and hunig's base (0.23 mL, 1.35 mmol, 5 eq) in DMF (5 mL) stirred under argon atmosphere at room temperature. A solution of crude *p*-nitrophenol ester (144 mg, 0.37 mmol, 1.4 eq) in DMF (4 mL) was added dropwise to the reaction mixture. The reaction mixture was stirred for 6 h. The solvent DMF was evaporated *in vacuo* to obtain the crude desired product. The crude solid was purified over silica gel chromatography (DCM:MeOH 0-25%) to obtain the desired product **11** as an light yellow solid (200 mg, 0.19 mmol, 73%). $^{1}$H NMR (400 MHz, DMSO) δ 8.04 (t, *J* = 5.2 Hz, 1H), 7.98 (broad s, 2H), 7.56 (broad s, 2H), 7.35 (broad s, 2H), 7.31 (d, *J* = 5.9 Hz, 2H), 6.56 (d, *J* = 8.5 Hz, 2H), 4.58 – 4.52 (m, 4H), 4.38 – 4.34 (m, 4H), 4.10 (t, *J* = 5.90 Hz, 2H), 3.85 (s, 3H), 3.19 – 3.14 (m, 2H), 2.35 – 2.20 (m, 4H), 2.09 (s, 6H), 1.86 (broad s, 6H), 1.80 (s, 3H), 1.29 (t, *J* = 7.0 Hz, 6H), 1.25 (t, *J* = 7.3 Hz, 3H). HRMS (ESMS) Calculated for $C_{47}H_{56}N_{14}O_{7}S_{3}$ $[M+H]^{+}$: 1025.3691, found 1025.3685.

**General method of RAFT Polymerization:** Reversible addition-fragmentation chain transfer (RAFT) was used to synthesize three analogs of polymers with distinct molar masses (25k, 50k, 175k). For synthesis dimethyl acrylamide (DMA) monomer was filtered over activated alumina and allowed to react under inert atmosphere in DMF (30 wt% monomer) at 70 ºC for 24 h in an oil bath. The initial monomer ($[M]_o$) to diABZI-CTA compound **11** ($[CTA]_o$) to initiator ($[I]_o$) ratio was n:1:0.2. The resultant desired polymers DMA(nk)-diABZI-CTA was isolated by dialysis against pure acetone (2×), pure deionized water (2×). Following dialysis, the purified compound was frozen at -80 ºC for 8 h and then lyophilized for 3 days.

diABZI-DMA[25KDa] (**12**)**:** The initial monomer ($[M]_o$) to diABZI-CTA compound **11** ($[CTA]_o$) to initiator ($[I]_o$) ratio was 242:1:0.2. The desired polymer (**12**) was obtained after dialysis as white solid (255 mg, 84%) which was further characterized by $^1$H NMR. $^1$H NMR (400 MHz, DMSO) δ 3.06 – 2.64 (dimethyl group PDMA), 2.62 – 2.11 (-CH- backbone of PDMA), 1.67 – 1.00 (-$CH_2$- backbone of PDMA),

diABZI-DMA[50KDa] (**13**)**:** The initial monomer ($[M]_o$) to diABZI-CTA compound **11** ($[CTA]_o$) to initiator ($[I]_o$) ratio was 525:1:0.2. The desired polymer (**13**) was obtained after dialysis as white solid (365 mg, 72%) which was further characterized by $^1$H NMR.

diABZI-DMA[175KDa] (**14**)**:** The initial monomer ($[M]_o$) to diABZI-CTA compound **11** ($[CTA]_o$) to initiator ($[I]_o$) ratio was 1755:1:0.2. The desired polymer (**14**) was obtained after dialysis as white solid (663 mg, 65%) which was further characterized by $^1$H NMR.

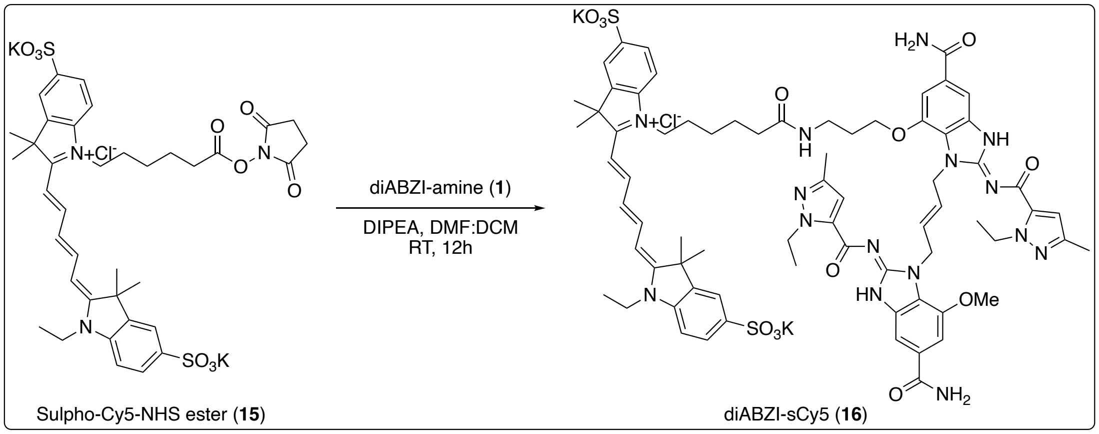


**Scheme S5:** Synthesis of diABZI-sCY5 (**16**)

**potassium 1-(6-((3-(((*E*)-6-carbamoyl-3-((*E*)-4-((*E*)-5-carbamoyl-2-((1-ethyl-3-methyl-1*H*-pyrazole-5-carbonyl)imino)-7-methoxy-2,3-dihydro-1*H*-benzo[*d*]imidazol-1-yl)but-2-en-1-yl)-2-((1-ethyl-3-methyl-1*H*-pyrazole-5-carbonyl)imino)-2,3-dihydro-1*H*-benzo[*d*]imidazol-4-yl)oxy)propyl)amino)-6-oxohexyl)-2-((1*E*,3*E*)-5-((*E*)-1-ethyl-3,3-dimethyl-5-sulfonatoindolin-2-ylidene)penta-1,3-dien-1-yl)-3,3-dimethyl-3*H*-indol-1-ium-5-sulfonate chloride (16).** To a stirred solution of compound **1** (18 mg, 16 μmol, 1.0 eq) and hunig's base (8.4

μL, 48 μmol, 3 eq) in DMF (1.5 mL) dropwise add solution of activated NHS ester compound **15**[5] in DMF (1.0 mL). The reaction mixture stirred overnight and precipitated by adding diethyl ether to obtain the desired product diABZI-sCy5 (**16**) as a blue solid (17.2 mg, 11.2 μmol, 70%). $^1H$ NMR (400 MHz, DMSO) δ 8.33 (t, *J* = 13.0 Hz, 2H), 7.99 (broad s, 2H), 7.82 – 7.79 (m, 3H), 7.66 – 7.61 (m, 4H), 7.35 – 7.28 (m, 8H), 6.61 – 6.44 (m, 3H), 6.35 – 6.21 (m, 2H), 5.88 – 5.70 (m, 2H), 5.00 – 4.83 (m, 4H), 4.52 – 4.45 (m, 4H), 4.14 – 3.93 (m, 6H), 3.72 (s, 3H), 3.11 – 3.05 (m, 2H), 2.09 (s, 3H), 2.08 (s, 3H), 2.00 (t, *J* = 7.4 Hz, 2H), 1.69 – 1.65 (m, 13H), 1.51 – 1.46 (m, 2H), 1.29 – 1.20 (m, 14H). HRMS (ESMS) Calculated for $C_{71}H_{84}N_{15}O_{13}S_2$ $[M+H]^{2+}/2$ = 709.7941, found 709.7946.

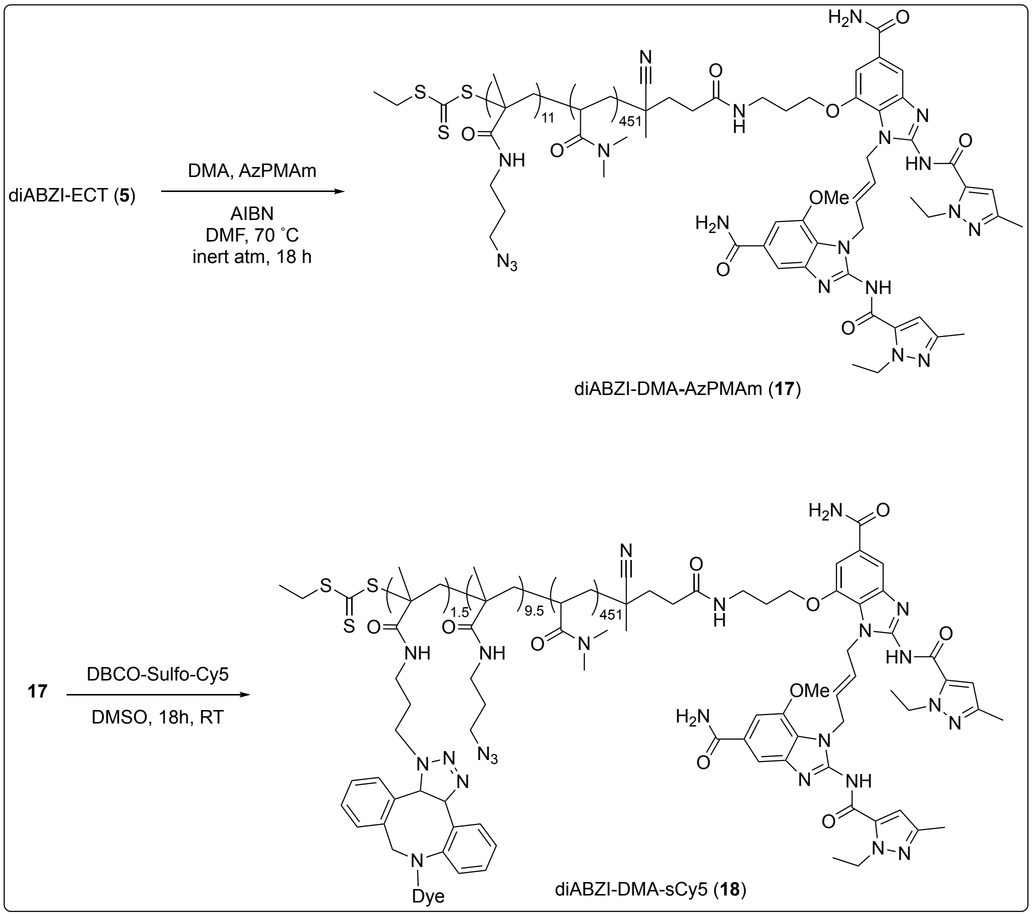


**Scheme S6:** Synthesis of diABZI-DMA-sCy5 (**18**)

RAFT polymerization was used to synthesize Compound **17**. For synthesis, DMA was filtered over activated alumina, combined with a solution of diABZI-ECT, AIBN, and AzPMAm, and allowed to react under inert atmosphere in DMF (30 wt% monomer) at 70 ºC for 24 h in an oil bath. The initial DMA to AzPMAm to diABZI-ECT to AIBN ratio was 496:11:1:0.2. The resultant desired polymer, diABZI-DMA-AzPMAm, was isolated by dialysis against pure acetone (2×), 1:1 pure acetone and deionized water (1x), and then pure deionized water (2×). Following dialysis, the purified compound was frozen at -80 ºC for 8 h and then lyophilized for 72 hours (494 mg, 91%), which was further characterized by $^1$H NMR and GPC in HPLC-grade DMF with 0.01 M LiBr via Light Scattering (TOSOH). GPC analysis showed a monodisperse polymer (PDI = 1.10) at a 69.0 kDa molecular weight.

Copper-free click chemistry was utilized to conjugate DBCO-Sulfo-Cy5 to Compound **17**, forming Compound **18**. Compounds **17** and DBCO-Sulfo-Cy5 were dissolved separately at 20 mg/mL in DMSO. A molar excess of 2.5:1 DBCO-Sulfo-Cy5 to polymer was utilized for the reaction. Briefly, ~63 µL of DBCO-Sulfo-Cy5 solution was added to 1200 µL of Compound **17** solution. The reaction was continuously stirred at room temperature in darkness for 24 hours. Compound **18** was purified via dialysis using 3.5 kDa Snakeskin cellulose tubing (Thermo) against pure acetone (2×), 1:1 pure acetone and deionized water (1x), pure deionized water (2×). Compound 18 was then lyophilized for 72 hours and stored at -20 ºC until use (~15 mg, ~53%). sCy5 concentration was determined using absorbance at 650 nm and diABZI concentration was determined at 325 nm using the NanoDrop UV-Vis Spectrophotometer (Thermo).

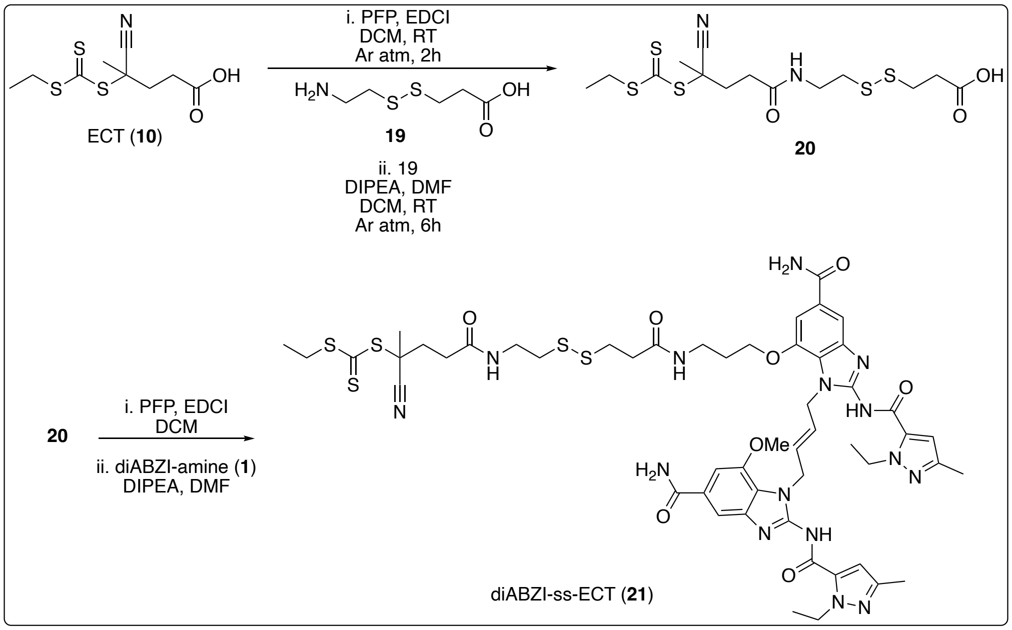


**Scheme S7:** Synthesis of diABZI-ss-ECT (**21**)

**3-((2-aminoethyl)disulfaneyl)propanoic acid--4-cyano-4-(((ethylthio)carbonothioyl)thio) pentanoic acid--6-cyano-6-methyl-9-oxo-4-thioxo-3,5,13,14-tetrathia-10-azaheptadecan-17-oic acid (1/1/1) (20).** To a stirred solution of 4-cyano-4-(((ethylthio)carbonothioyl)thio)pentanoic acid **10**[4] (2.10 g, 7.97 mmol, 1 eq) and 3-(((ethylimino)methylene)amino)-*N*,*N*-dimethylpropan-1-amine hydrogen chloride (1.68 g, 8.77 mmol, 1.1 eq) in 25 mL DCM dropwise added solution of 2,3,4,5,6-pentafluorophenol ( 1.46g, 7.97 mmol, 1 eq) in 5 mL DCM under inert atmosphere. The reaction mixture was stirred for 4 hours. After consumption of the starting material, as judged by the TLC analyses. The solution of 3-((2-aminoethyl)disulfaneyl)propanoic acid (**19**)[6] (2.25 g, 10.36 mmol, 1.3 eq) and triethyl amine (4.45 mL, 3.19 mmol, 4 eq) in 10 mL DCM added to the reaction mixture and stirred overnight. The solvent was evaporated *in vacuo* to obtain the crude desired product. The crude product was purified over silica gel chromatography (DCM:MeOH 0-5%) to obtain the desired product **20** as an pale yellow oil (2.1 g, 4.89 mmol, 61%). $^1$H NMR (600 MHz, $CDCl_3$) δ 6.35 (t, *J* = 5.3 Hz, 1H), 3.60 (q, *J* = 6.0 Hz, 2H), 3.35 (q, *J* = 7.4 Hz, 2H), 2.97 (t, *J* = 6.6 Hz, 2H), 2.84 (t, *J* = 6.3 Hz, 2H), 2.80 (t, *J* = 6.8 Hz, 2H) , 2.59 – 2.50 (m, 3H) , 2.44 – 2.31 (m, 1H), 1.90 (s, 3H), 1.36 (t, *J* = 7.44 Hz, 3H). $^{13}$C NMR (151 MHz, CDCl3) δ 217.1, 175.8, 171.1, 119.3, 46.7, 38.4, 37.5, 34.5, 34.0, 33.0, 31.8, 31.4, 24.9, 12.8. HRMS (ESMS) Calculated for $C_{14}H_{22}N_2O_3S_5$ $[M+H]^+$ = 427.0307, found 427.0308.

**(*E*)-5-((2-((3-((3-((5-carbamoyl-1-(4-(5-carbamoyl-2-(1-ethyl-3-methyl-1*H*-pyrazole-5-carboxamido)-7-methoxy-1*H*-benzo[*d*]imidazol-1-yl)but-2-en-1-yl)-2-(1-ethyl-3-methyl-1*H*-pyrazole-5-carboxamido)-1*H*-benzo[*d*]imidazol-7-yl)oxy)propyl)amino)-3-oxopropyl)disulfaneyl)ethyl)amino)-2-cyano-5-oxopentan-2-yl ethyl carbonotrithioate (21).** To a stirred solution of compound (**20**) (80 mg, 187.5 µmol, 1eq) and 3-(((ethylimino)methylene)amino)-*N*,*N*-dimethylpropan-1-amine hydrogen chloride (43 mg, 225 µmol, 1.1eq) in DCM 2 mL dropwise added solution of 2,3,4,5,6-pentafluorophenol ( 34 mg, 187 µmol, 1eq) in 1 mL DCM under inert atmosphere. The reaction mixture was stirred for additional 4 hours. After consumption of the starting material, as judged by the TLC analyses. The solution of diABZI-amine (**1**) (189 mg, 168 µmol, 0.9 eq) and hunig's base (163 µL, 937 µmol, 5eq) in 4 mL DMF added to the reaction mixture and stirred overnight. The diethyl ether added to precipitate out the crude solid desired product. The crude solid was purified over silica gel chromatography (DCM:MeOH 0-25%) to obtain the desired product **21** as an light yellow solid (122 mg, 102 µmol, 55%). $^{1}$H NMR (400 MHz, DMSO) δ 8.22 (t, *J* = 5.6 Hz, 1H), 8.00 – 7.96 (m, 3H), 7.64 (d, *J* = 4.5 Hz, 2H), 7.36 – 7.28 (m, 4H), 6.50 (d, *J* = 9.6 Hz, 2H), 5.90 – 5.78 (m, 2H), 4.94 – 4.89 (m, 4H), 4.53 – 4.49 (m, 4H), 3.98 (t, *J* = 5.8 Hz, 2H), 3.72 (s, 3H), 3.14 – 3.11 (m, 2H), 2.85 (t, *J* = 7.1 Hz, 2H), 2.75 (t, *J* = 6.6 Hz, 2H), 2.43 (t, *J* = 7.2 Hz, 2H), 2.40 – 2.26 (m, 4H), 2.10 (s, 3H), 2.09 (s, 3H), 1.82 (s, 3H), 1.72 – 1.67 (m, 2H), 1.27 – 1.24 (m, 9H). $^{13}$C NMR (151 MHz, DMSO) δ 218.7, 170.5, 170.4, 168.1, 168.0, 145.5, 145.3, 144.7, 130.4, 128.6, 128.2, 119.7, 109.7, 66.5, 56.4, 47.5, 46.1, 38.4, 37.5, 35.8, 35.5, 34.3, 34.2, 31.5, 31.0, 29.1, 24.3, 16.6, 13.6, 13.2. HRMS (ESMS) Calculated for $C_{52}H_{65}N_{15}O_8S_5$ $[M+H]^+$ = 1188.3817, found 1188.3824.

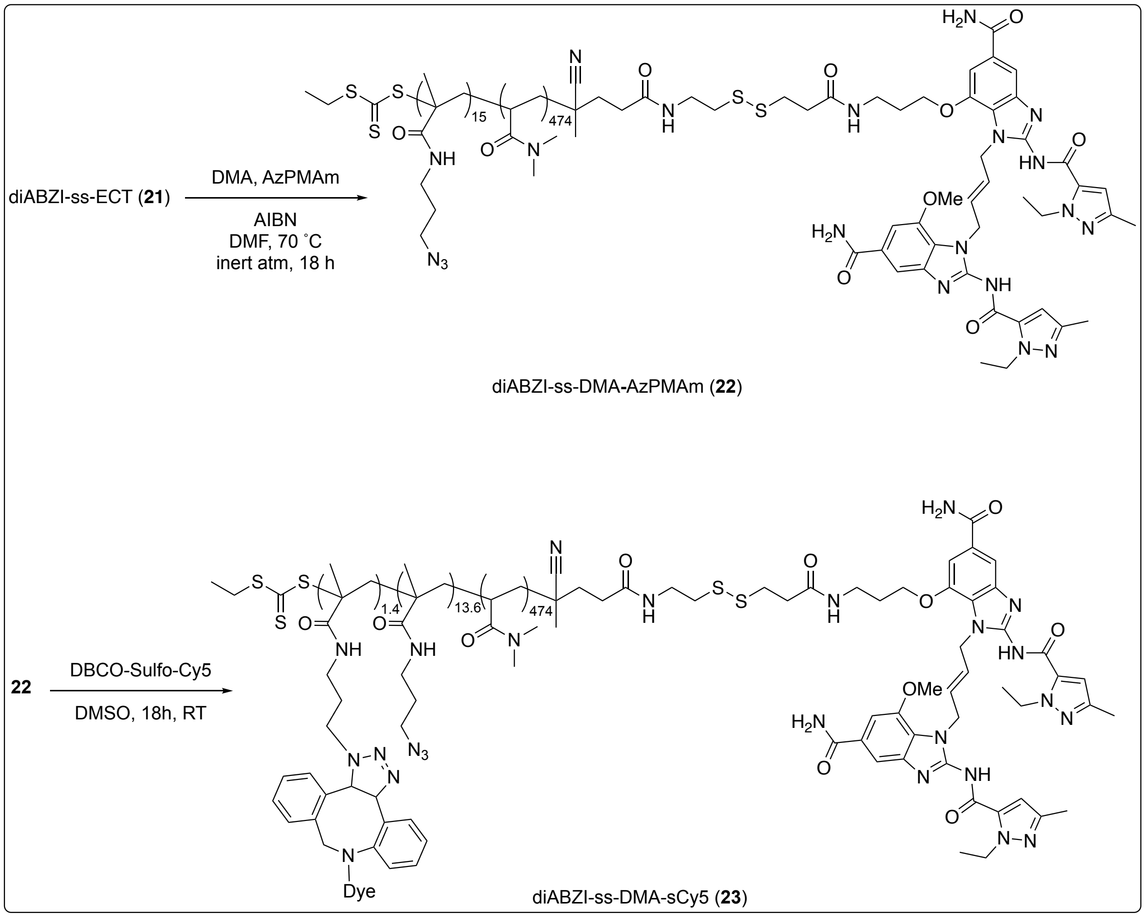


**Scheme S8:** Synthesis of diABZI-ss-DMA-sCy5 (**23**)

RAFT polymerization was used to synthesize Compound **22**. For synthesis, dimethyl acrylamide (DMA) monomer was filtered over activated alumina, combined with a solution of diABZI-ss-ECT, AIBN, and AzPMAm, and allowed to react under inert atmosphere in DMF (30 wt% monomer) at 70 ºC for 24 h in an oil bath. The initial DMA to AzPMAm to diABZI-SS-ECT to AIBN ratio was 490:15:1:0.2. The resultant desired polymer diABZI-ss-DMA-AzPMAm was isolated by dialysis against pure acetone (2×), 1:1 pure acetone and deionized water (1x), pure deionized water (2×). Following dialysis, the purified compound was frozen at -80 ºC for 8 h and then lyophilized for 3 days (154 mg, 76%) which was further characterized by $^{1}H$ NMR and GPC in HPLC-grade DMF with 0.01 M LiBr via Light Scattering (TOSOH). GPC analysis showed a monodisperse polymer (PDI = 1.05) at a 60.3 kDa molecular weight.

Copper-free click chemistry was utilized to conjugate DBCO-Sulfo-Cy5 to Compound **22**, forming Compound **23**. Compounds **22** and DBCO-Sulfo-Cy5 were dissolved separately at 20 mg/mL in DMSO. A molar excess of 2.5:1 DBCO-Sulfo-Cy5 to polymer was utilized for the reaction. Briefly, ~63 uL of DBCO-Sulfo-Cy5 solution was added to 1200 uL of Compound 22 solution. The reaction was continuously stirred at room temperature in darkness for 24 hours. Compound 23 was purified via dialysis using 3.5 kDa Snakeskin cellulose tubing (Thermo) against pure acetone (2×), 1:1 pure acetone and deionized water (1x), pure deionized water (2×). Compound 23 was then lyophilized for 72 hours and stored at -20 ºC until use (~15 mg, ~53%). sCy5 concentration was determined using absorbance at 650 nm and diABZI concentration was determined at 325 nm using the NanoDrop UV-Vis Spectrophotometer (Thermo).

**Part C: Supplementary Data**

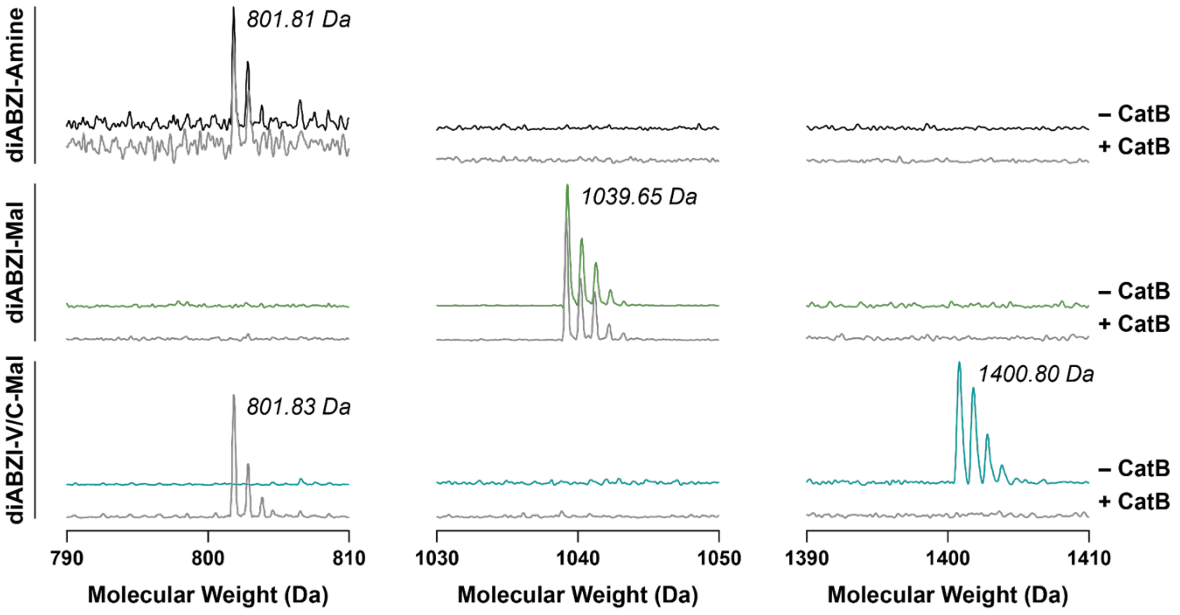


**Figure S1.** Analysis of cathepsin B-mediated linker cleavage of diABZI-V/C-Mal with diABZI-Mal and diABZI-Amine as controls. Molecules were incubated at 50 µM diABZI in a solution of cathepsin B at 37 ºC for 48 hours and MALDI-MS used to detect starting materials and expected cleavage products.

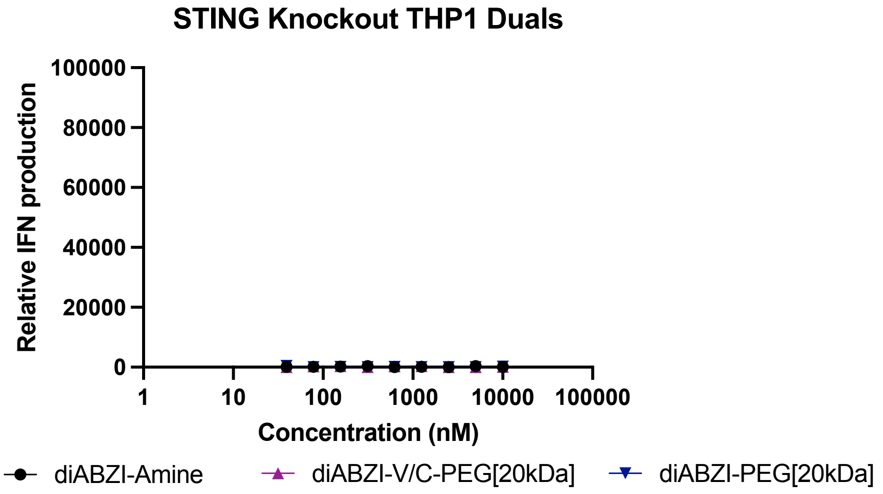


**Figure S2:** Dose-response curve for relative IFN-I production by STING knockout THP1-Dual reporter cells.

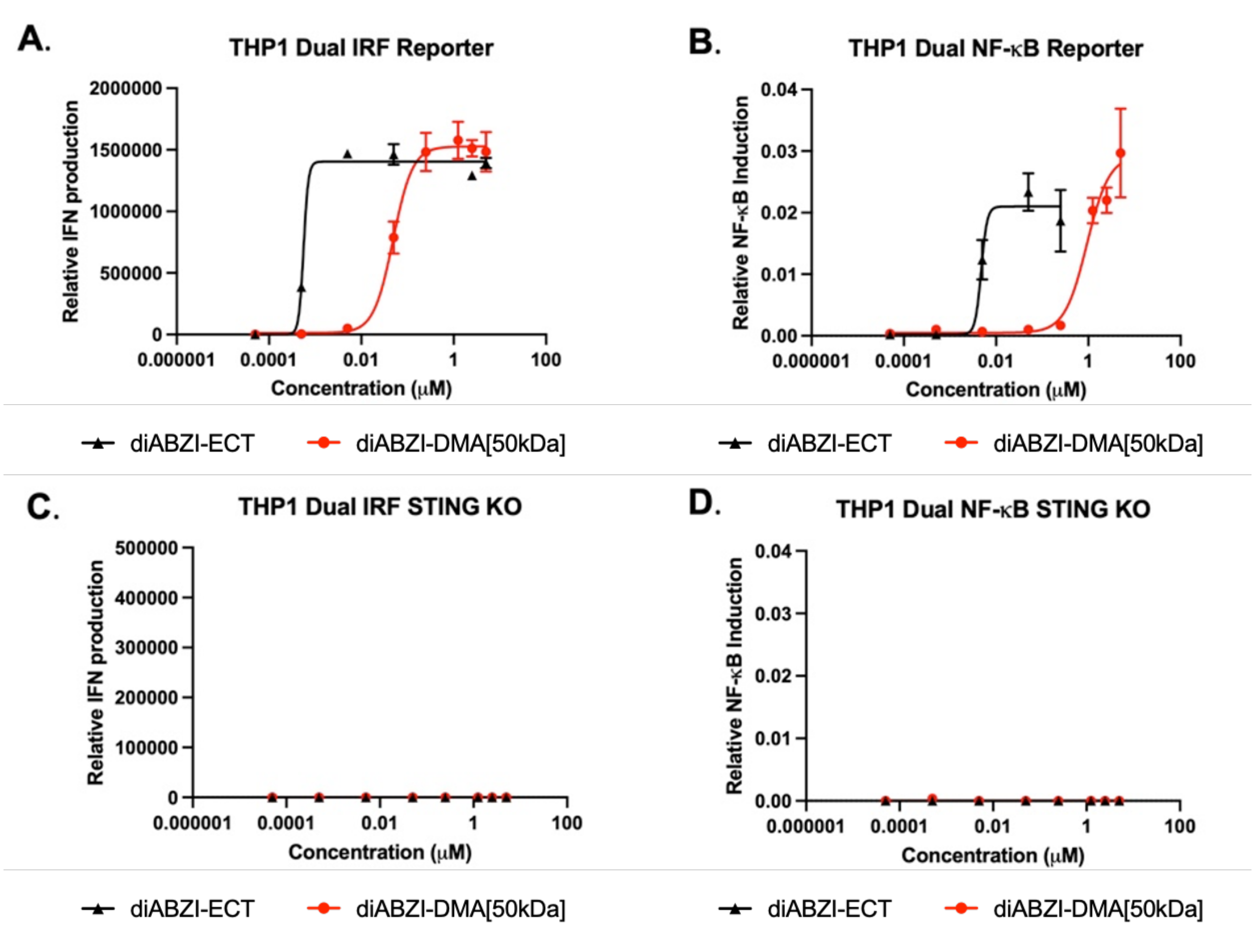


**Figure S3:** STING activation assays comparing the activity of diABZI-ECT and diABZI-DMA. **(A)** Dose-response curves for relative IFN-I production by THP1-Dual reporter cells treated with diABZI-ECT and diABZI-DMA[50kDa]. **(B)** Dose-response curves for relative NF-κB production by THP1-Dual reporter cells treated with diABZI-ECT and diABZI-DMA[50kDa]. **(C)** Dose-response curve for relative IFN-I production by STING knockout THP1-Dual reporter cells. **(D)** Dose-response curve for relative NF-κB production by STING knockout THP1-Dual reporter cells. Dose-response curves were fit to a variable slope (four parameter) non-linear regression to estimate EC50 values.

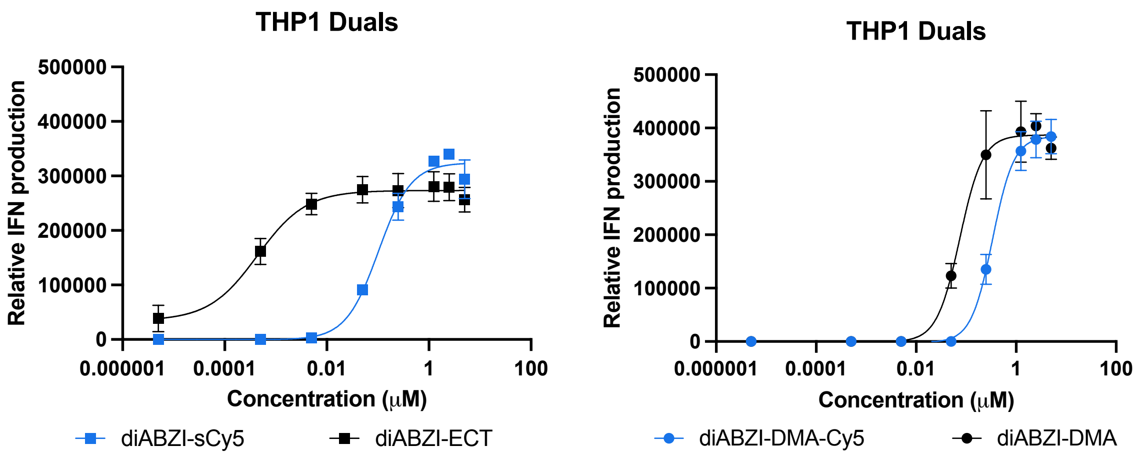


**Figure S4:** Dose-response curves for relative IFN-I production by THP1-Dual reporter cells treated with diABZI-ECT, diABZI-sCy5, diABZI-DMA-Cy5, and diABZI-DMA. Dose-response curves were fit to a variable slope (four parameter) non-linear regression to estimate EC50 values.

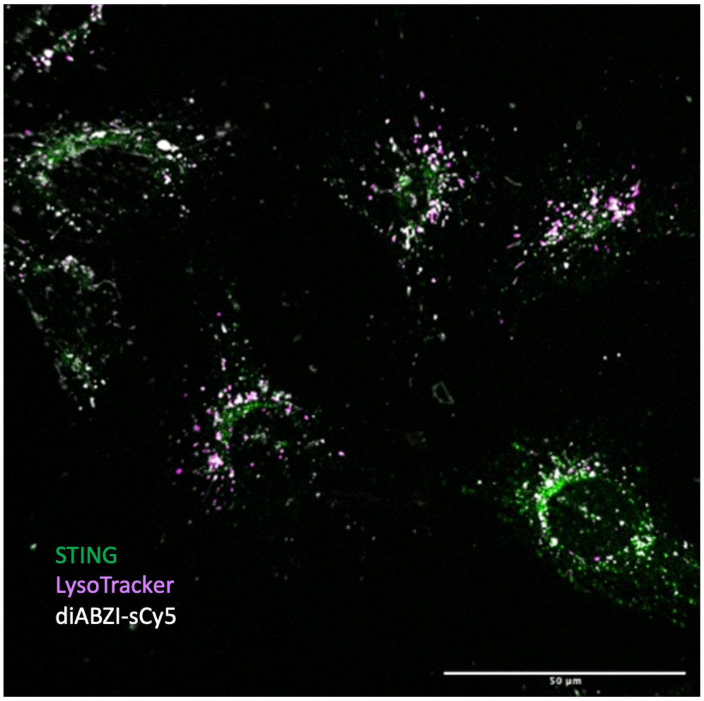


**Figure S5:** Representative image of lysotracker assay for diABZI-sCy5

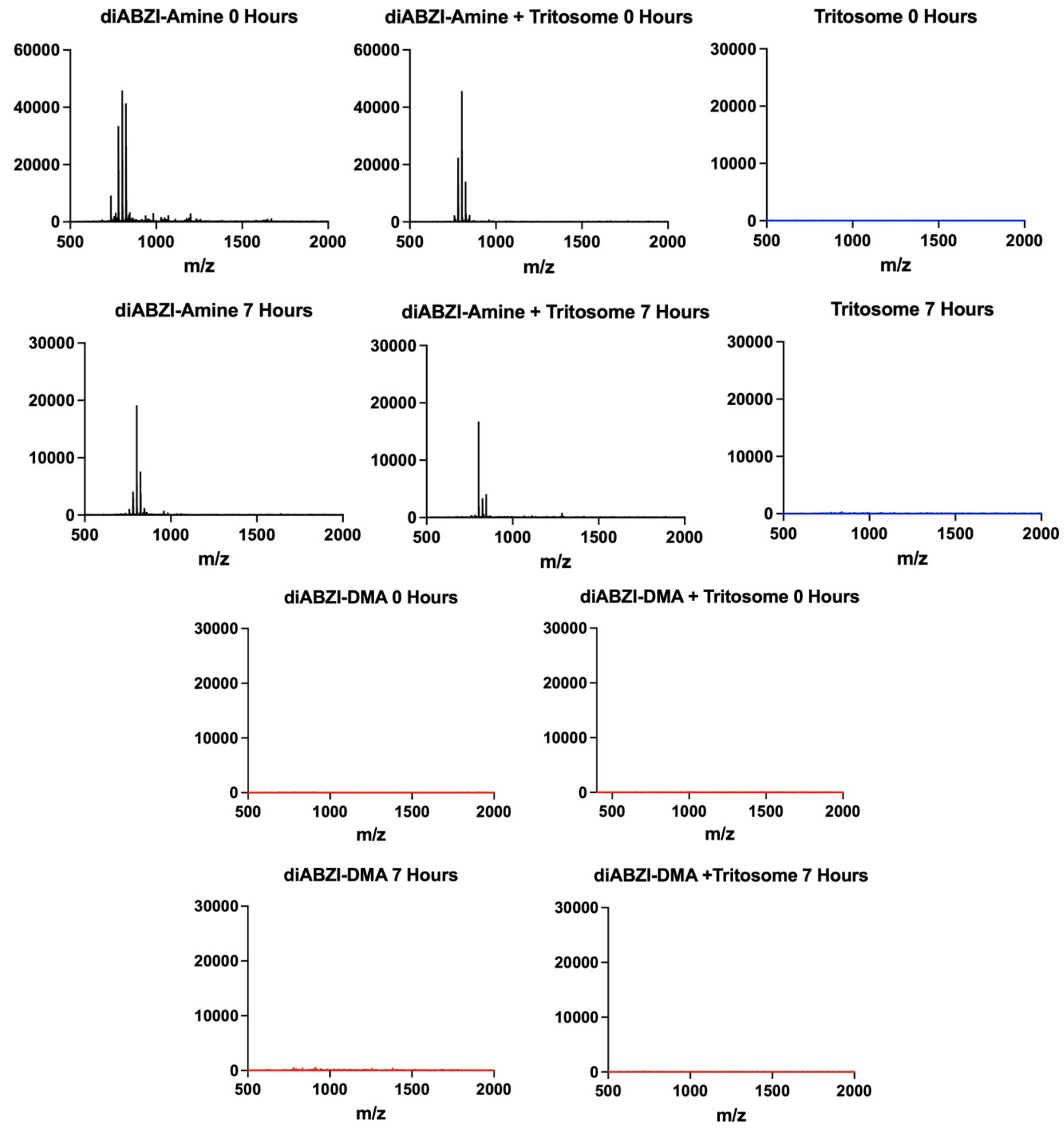


**Figure S6:** Analysis of diABZI-Amine and diABZI-DMA stability of in rat liver tritosomes measured using MALDI-MS. DiABZI-Amine or diABZI-DMA[50kDa] (100 μM) were incubated at 37°C for 7 hours in either water or 10% isolated rat lysosomes (tritosomes) and MALDI-MS was performed on the samples to detect diABZI amine or other related variants with m/z between 500-2000. After incubation in tritosomes for 7h, diABZI-amine remains stable and cannot be detected in diABZI-DMA[50kDa] sample, indicating lysosomal stability of diABZI-DMA.

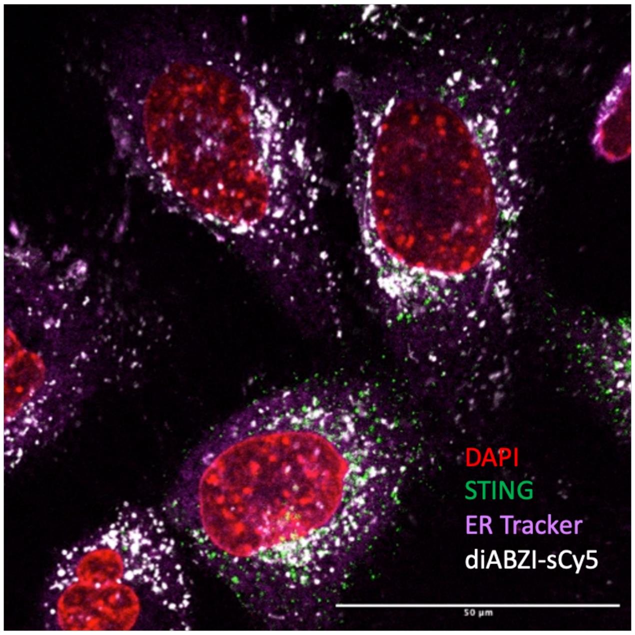


**Figure S7:** Representative image for ER Tracker assay for diABZI-sCy5

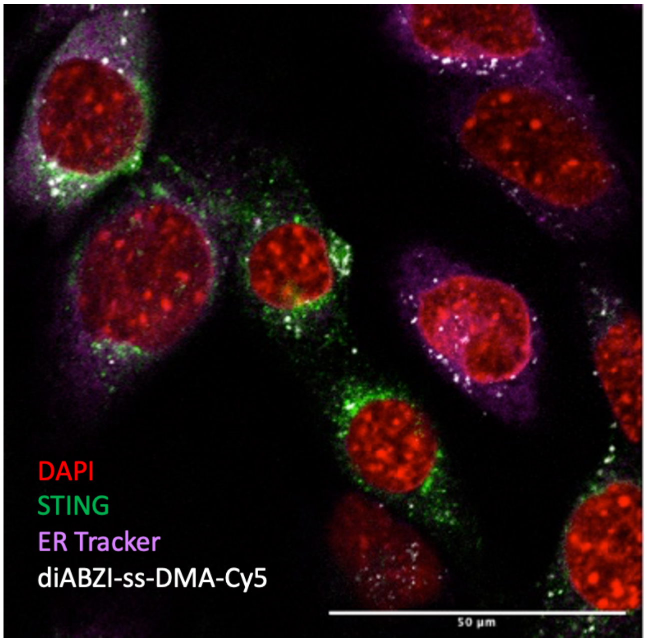


**Figure S8:** Representative image for ER tracker assay for diABZI-sCy5

**Part E: $^{1}$H NMR spectra for new compounds**

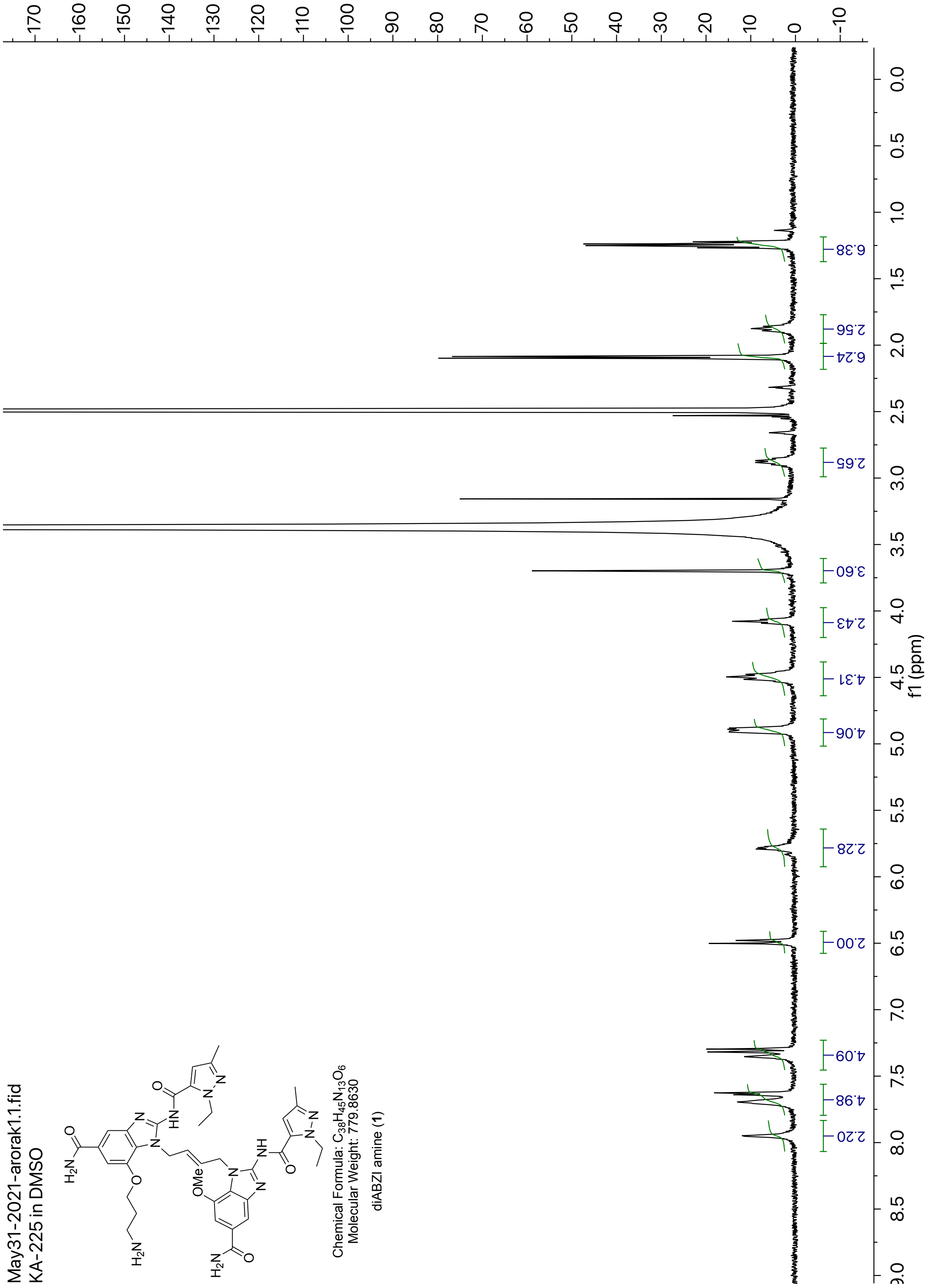


**Figure S9**: $^{1}$H NMR of diABZI amine (**1**) in DMSO

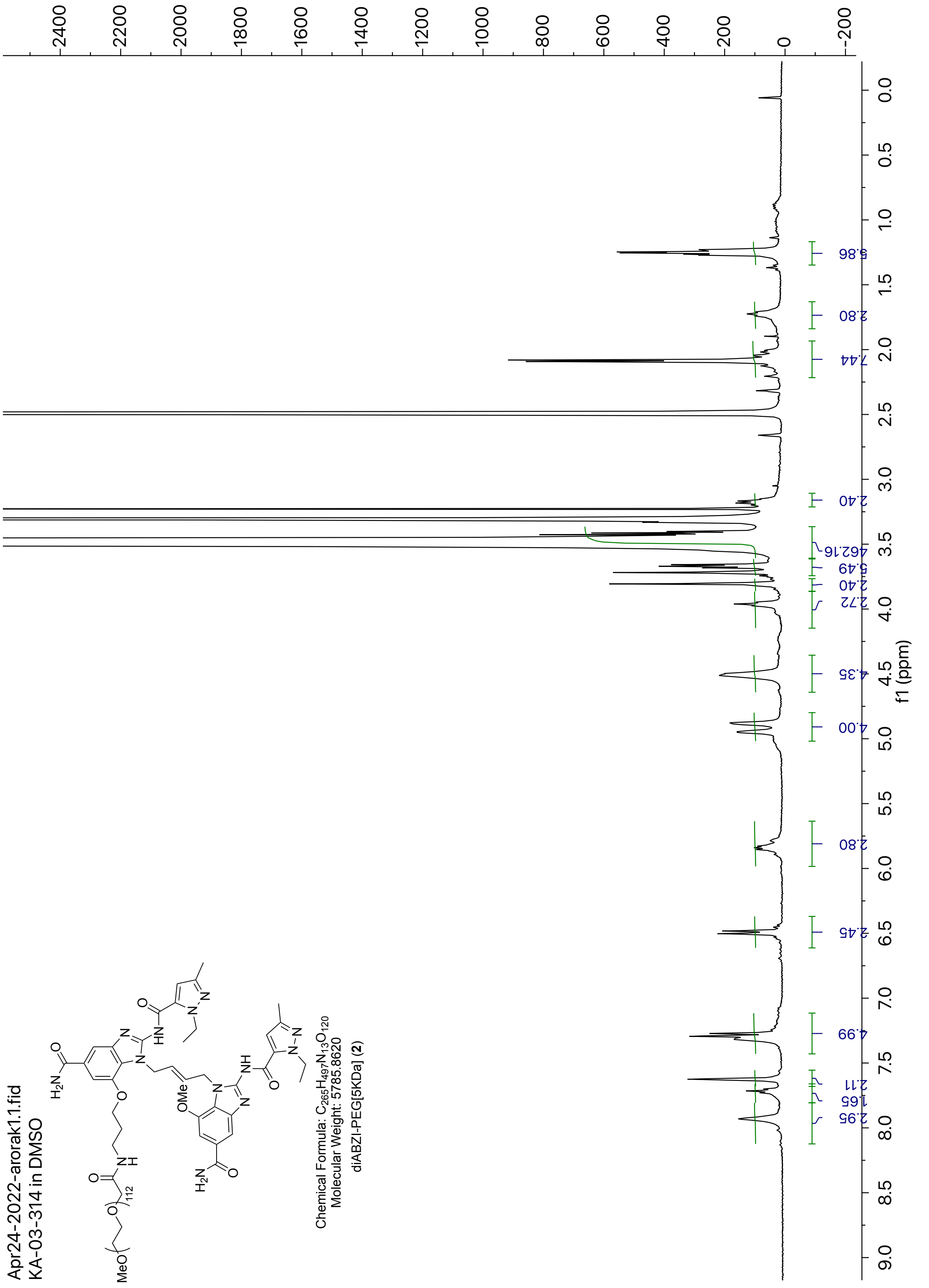


**Figure S10**: $^{1}$H NMR of diABZI-PEG[5KDa] (**2**) in DMSO

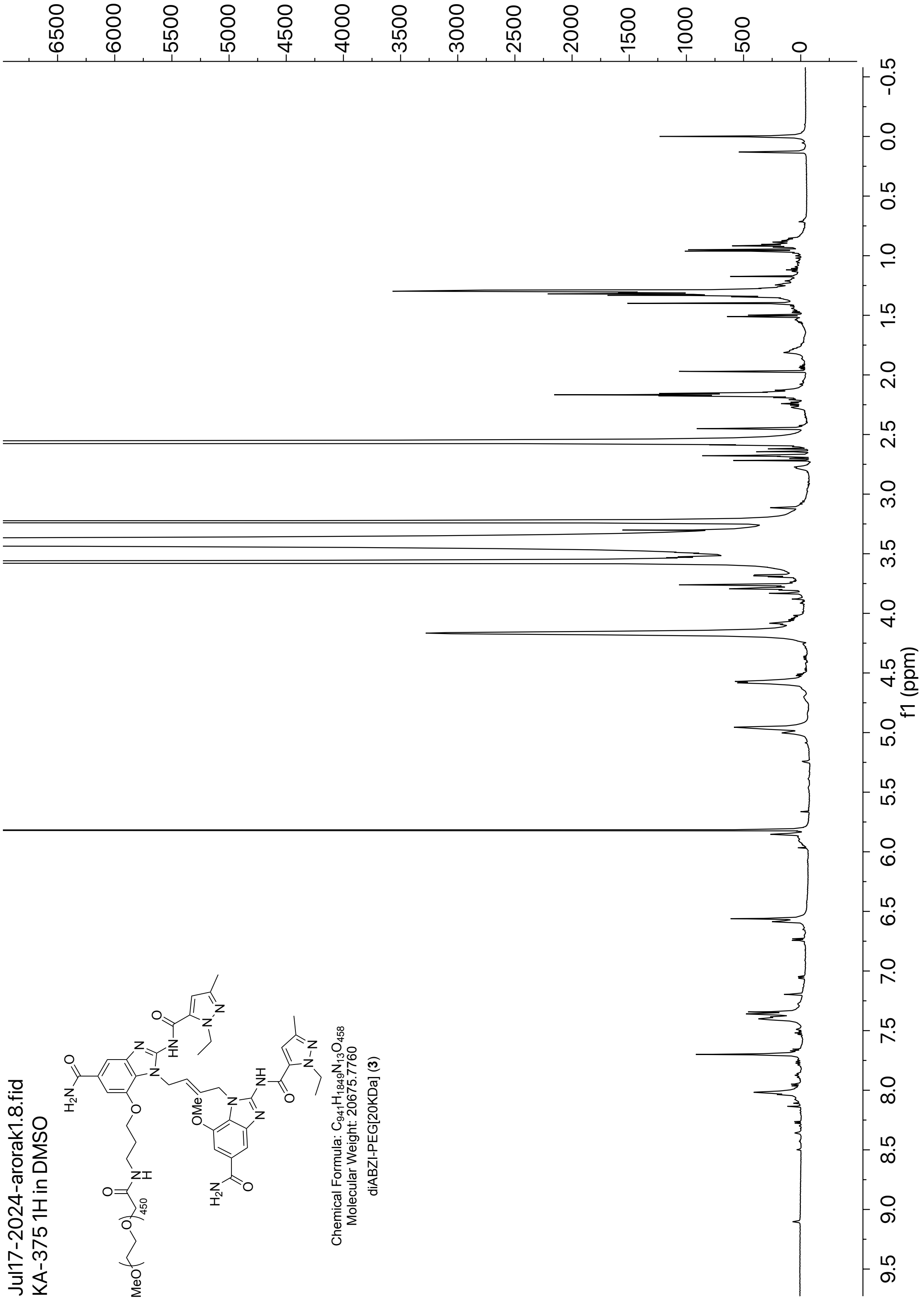


**Figure S11**: [1]H NMR of diABZI-PEG[20KDa] (**3**) in DMSO

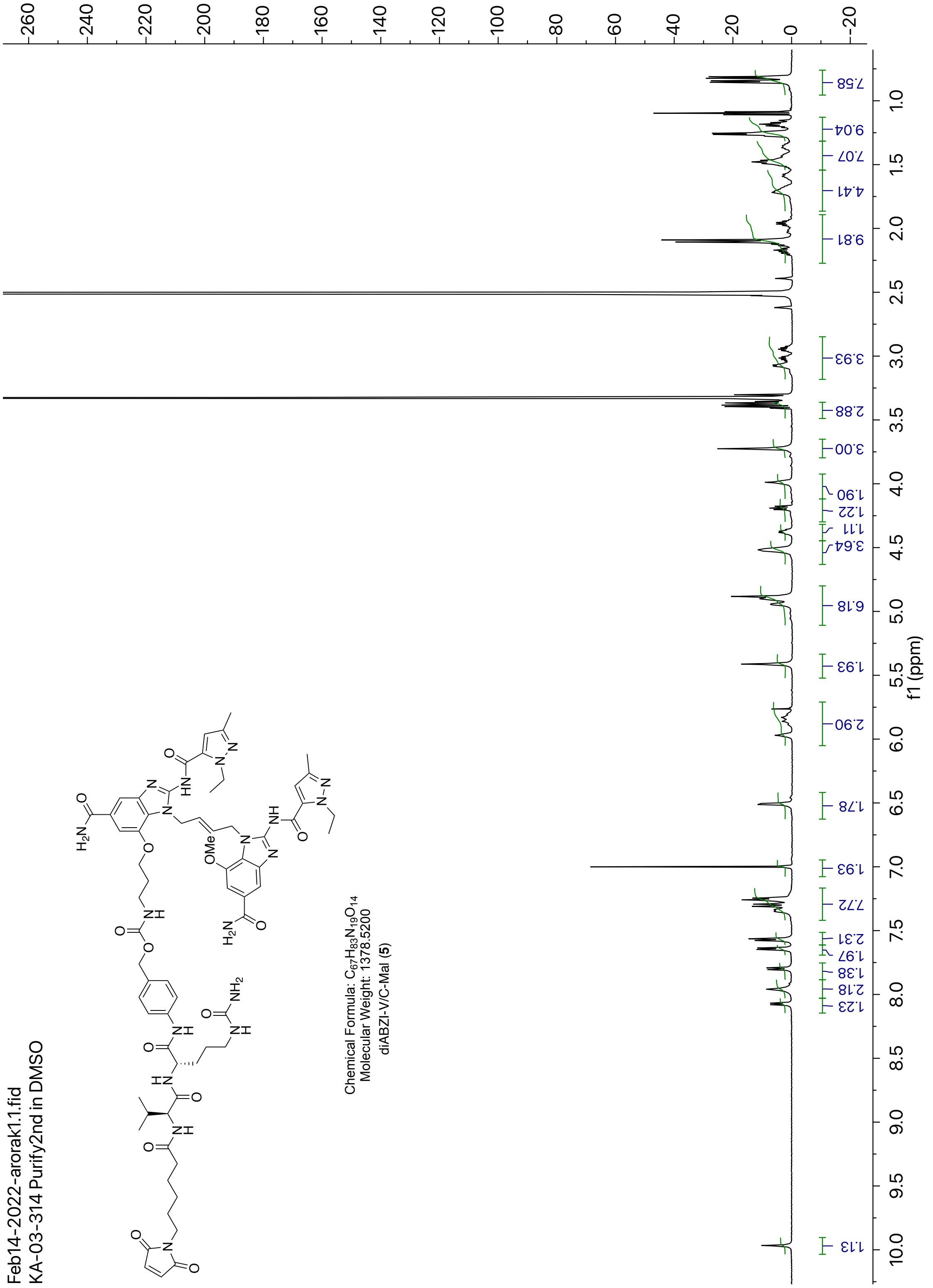


**Figure S12**: $^{1}$H NMR of diABZI-V/C-Mal (**5**) in DMSO

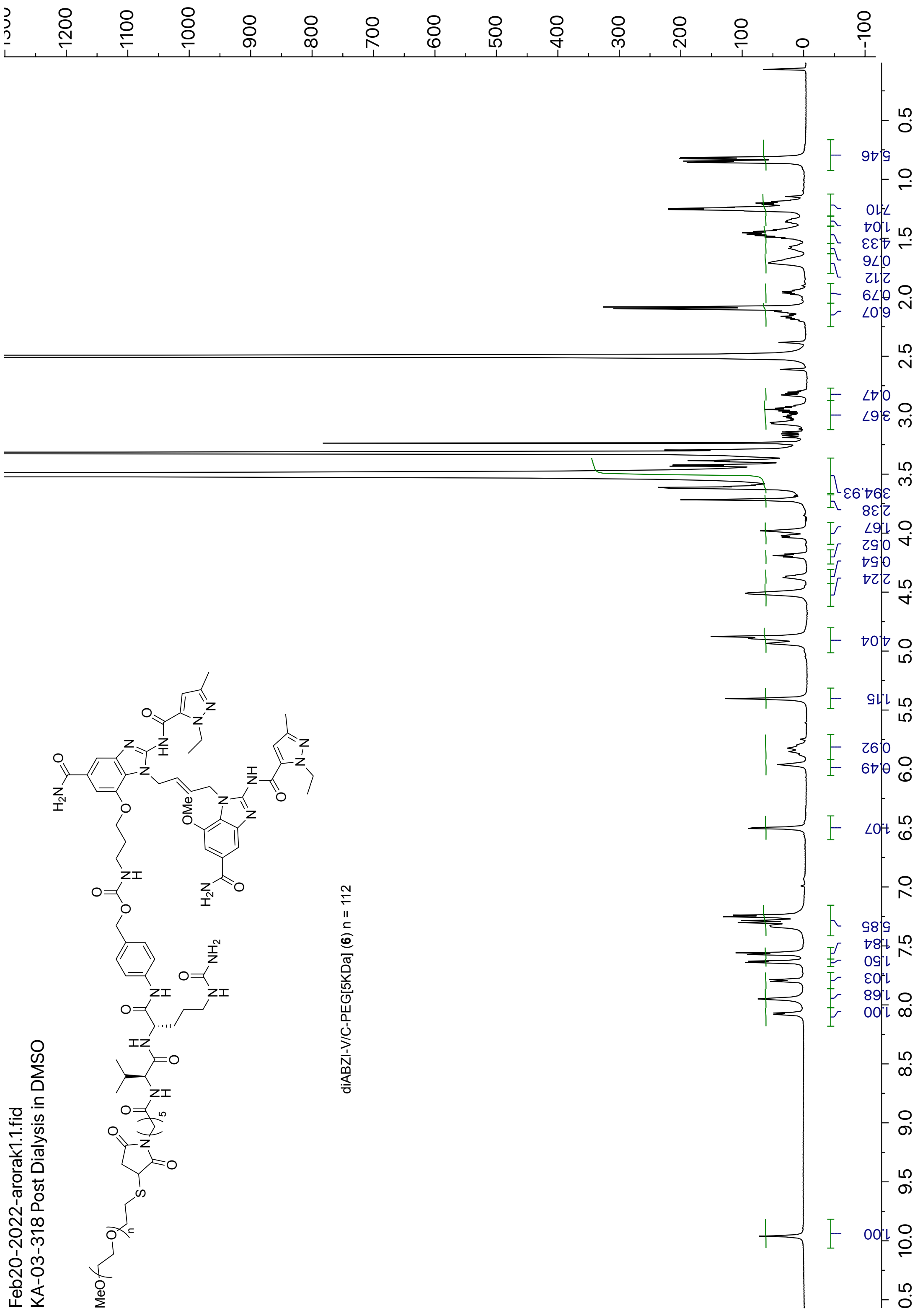


**Figure S13**: $^{1}$H NMR of diABZI-V/C-PEG[5KDa] (**6**) in DMSO

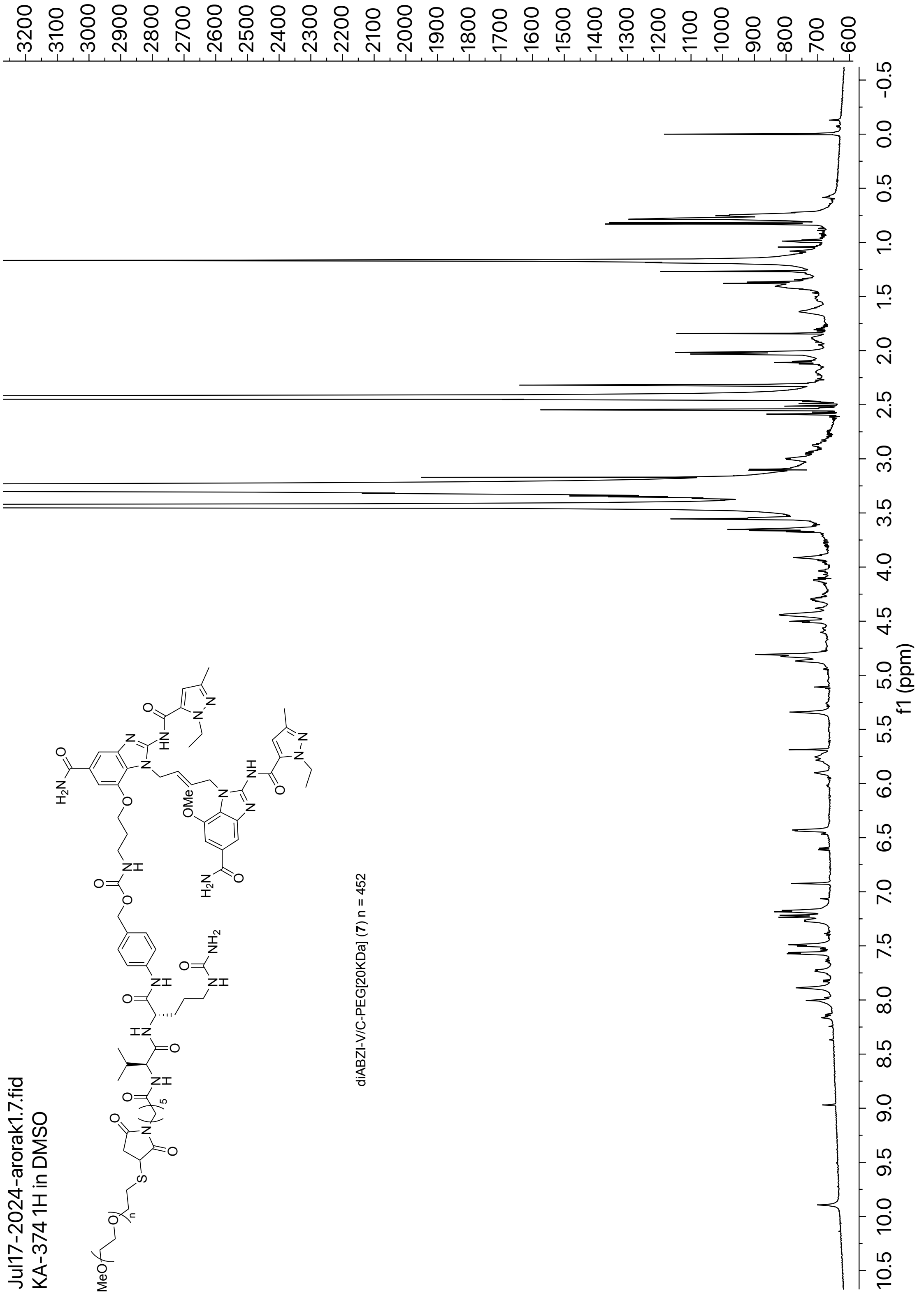


**Figure S14**: $^{1}$H NMR of diABZI-V/C-PEG[20KDa] (**7**) in DMSO

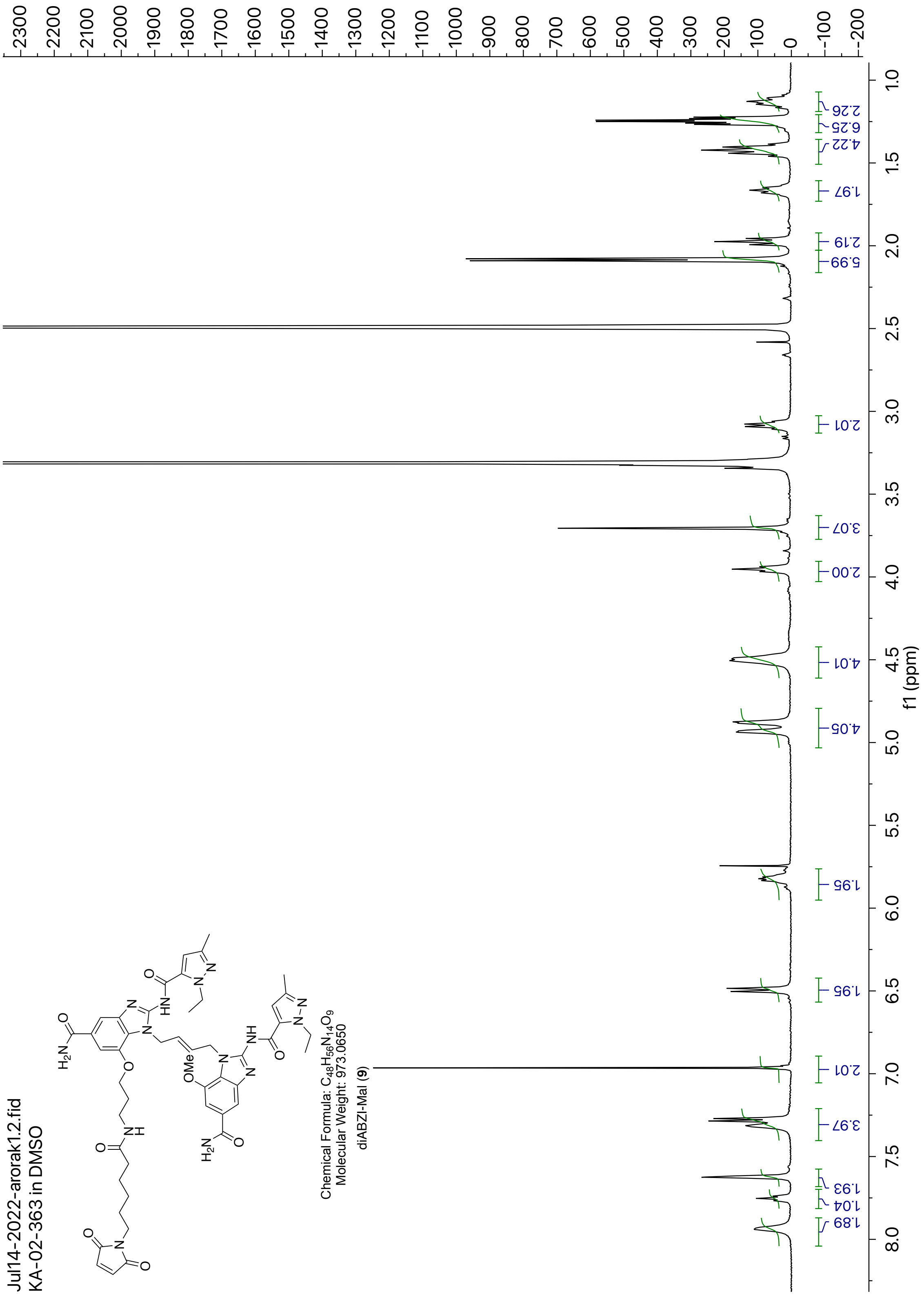


**Figure S15**: [1]H NMR of diABZI-Mal (**9**) in DMSO

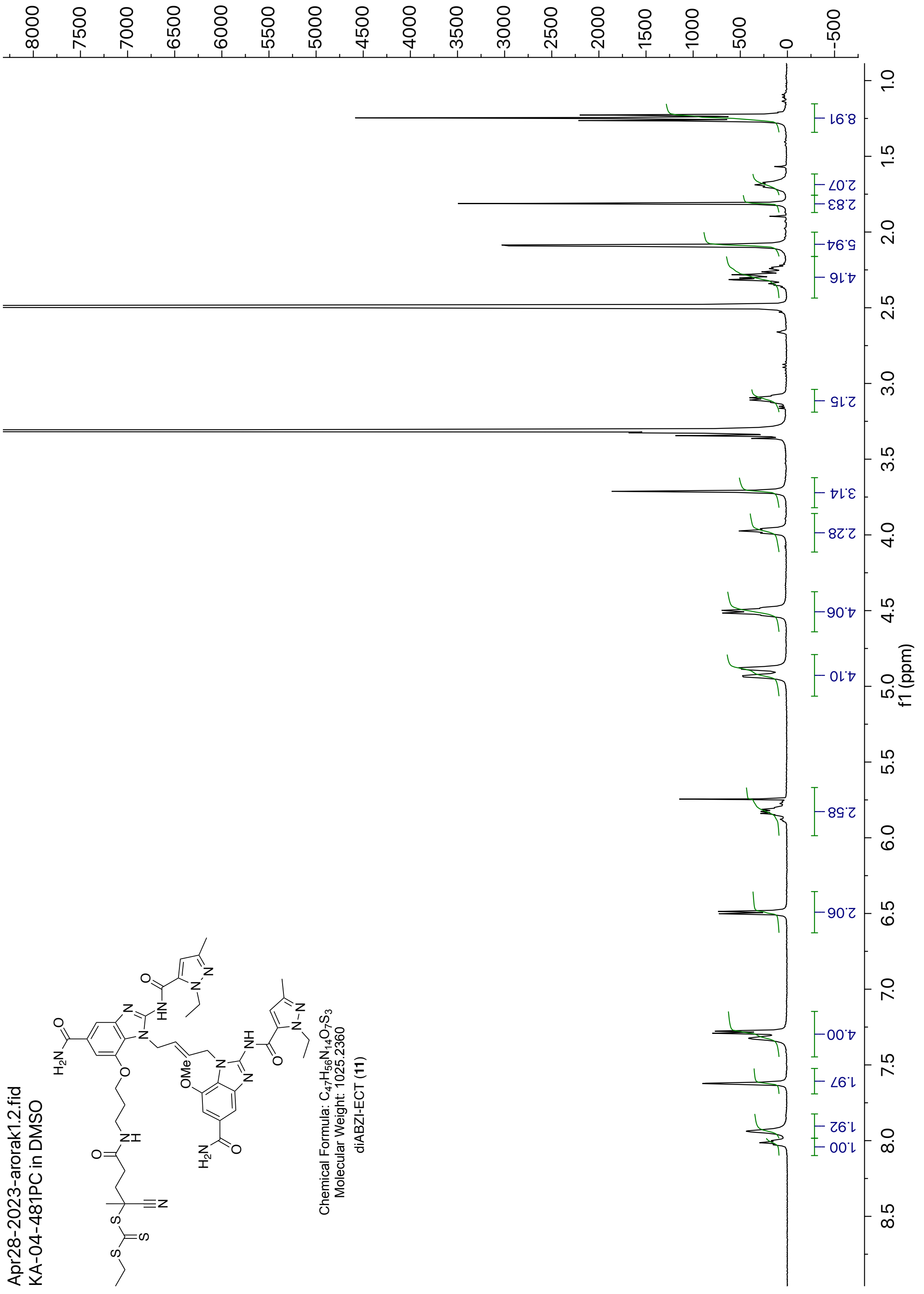


**Figure S16**: $^{1}$H NMR of diABZI-ECT (**11**) in DMSO

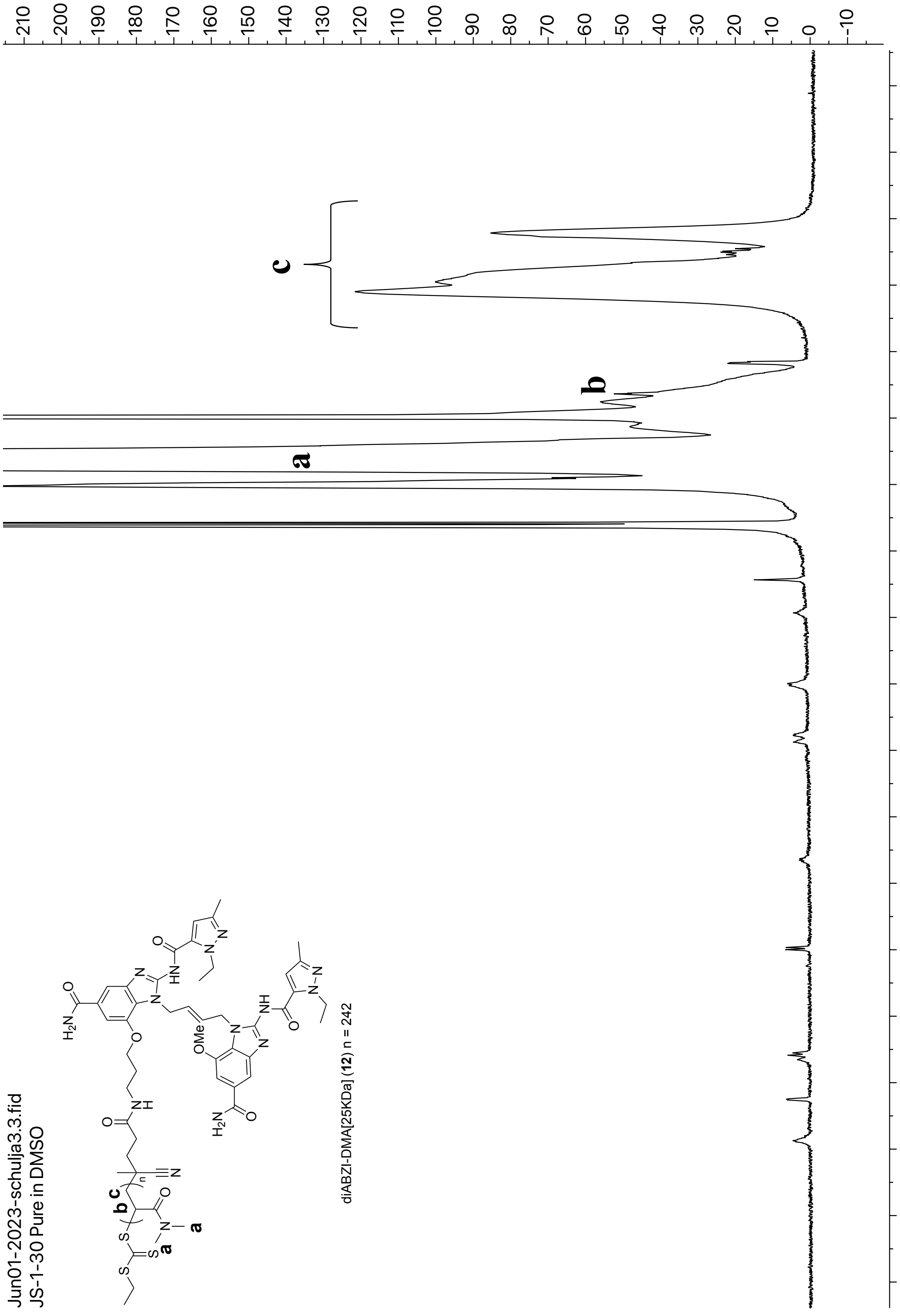


**Figure S17**: $^1$H NMR of diABZI-DMA[25KDa] (**12**) in DMSO

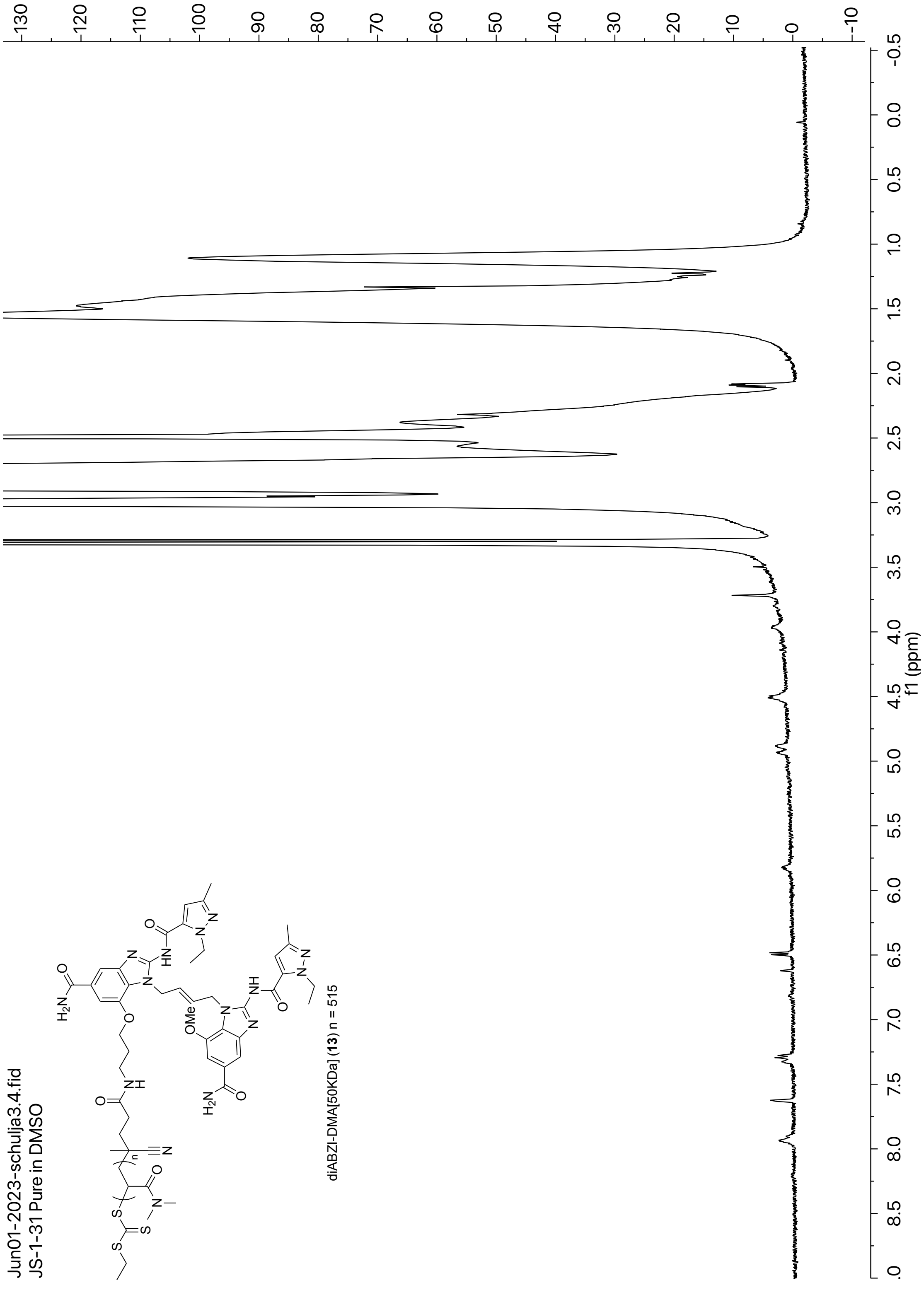


**Figure S18**: $^{1}$H NMR of diABZI-DMA[50KDa] (**13**) in DMSO

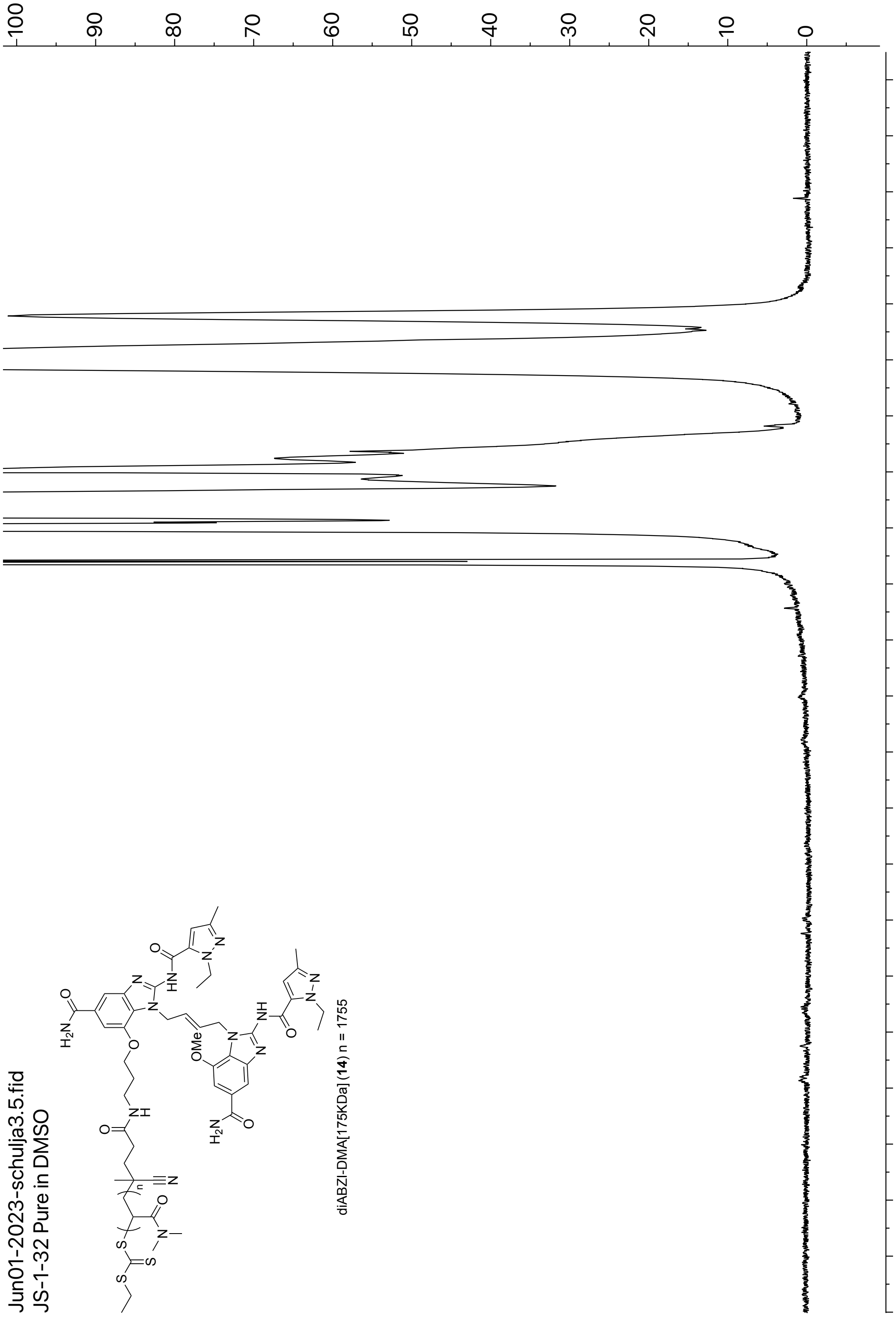


**Figure S19**: $^{1}$H NMR of diABZI-DMA[175KDa] (**14**) in DMSO

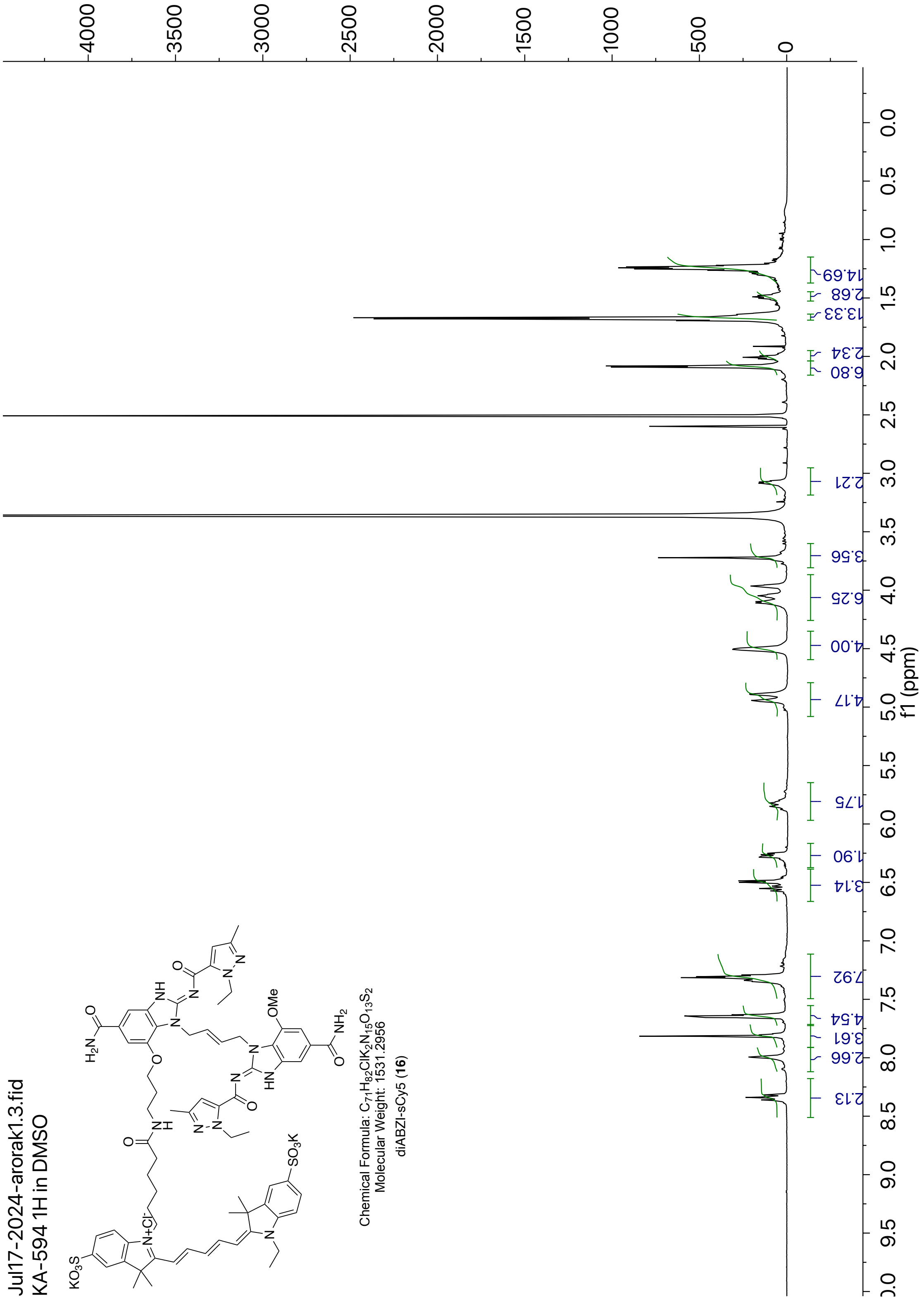


**Figure S20**: [1]H NMR of diABZI-sCy5 (**16**) in DMSO

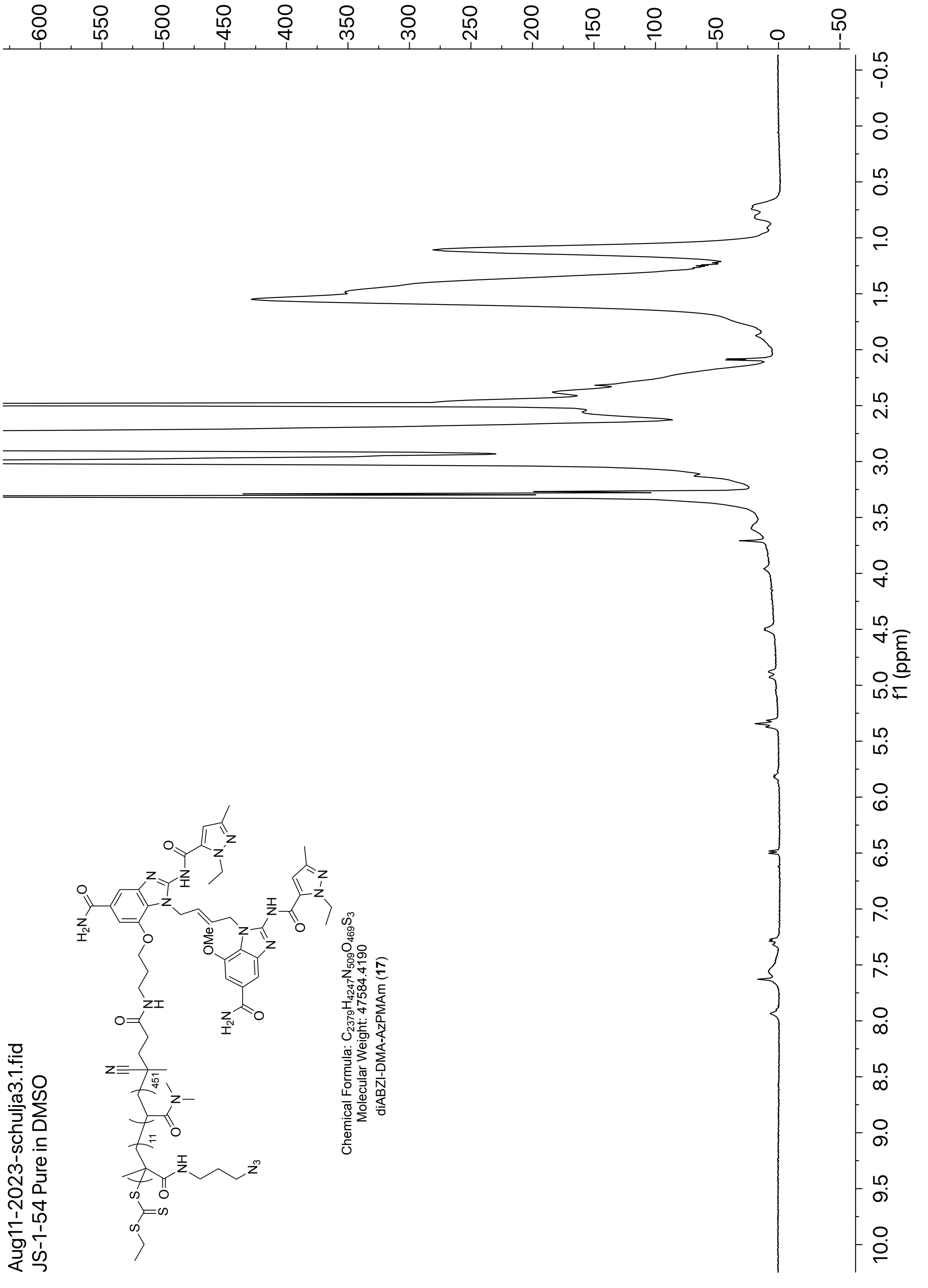


**Figure S21**: [1]H NMR of diABZI-DMA-AzPMAm (**17**) in DMSO

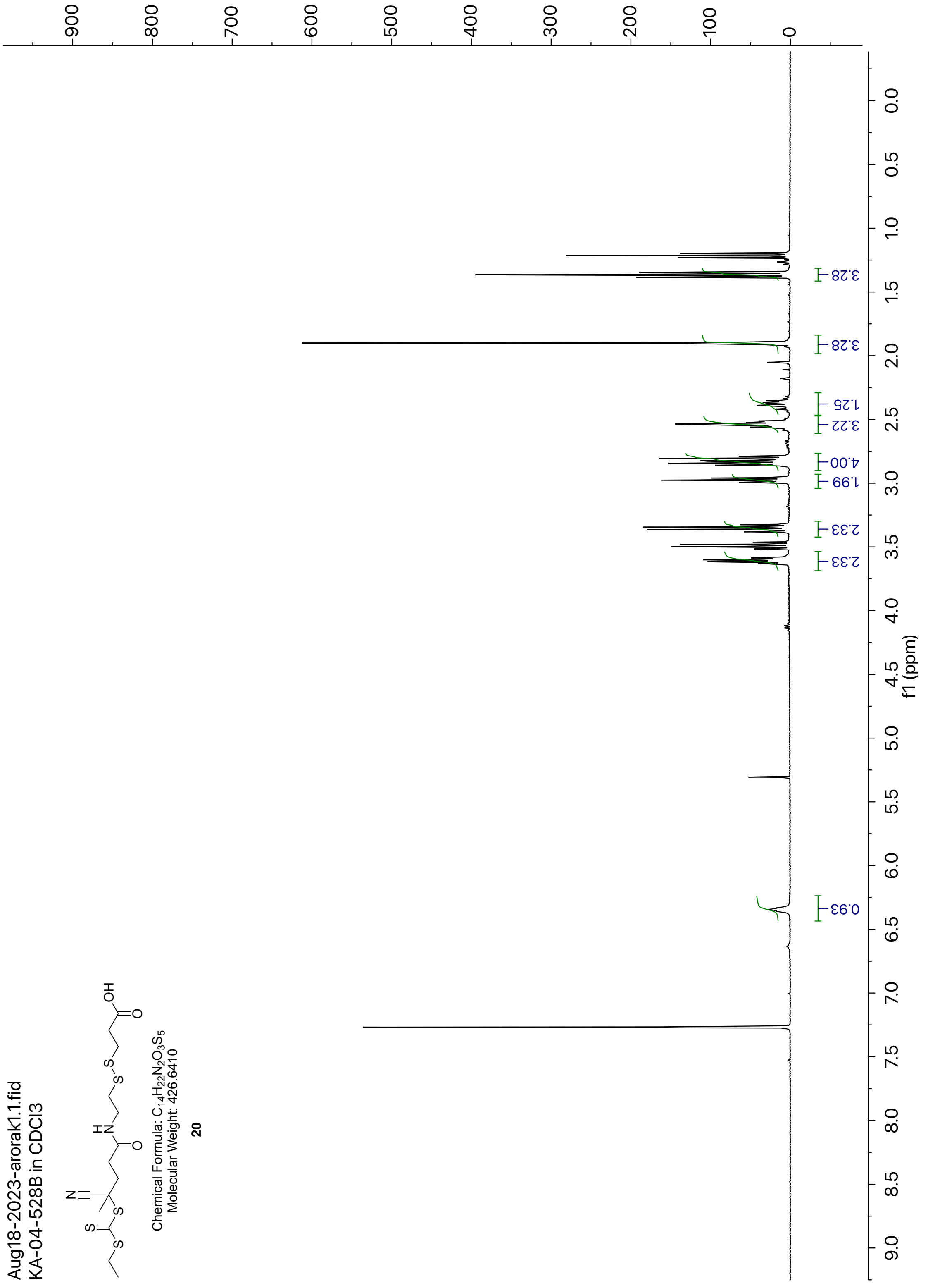


**Figure S22**: $^{1}$H NMR of compound (**20**) in $CDCl_3$

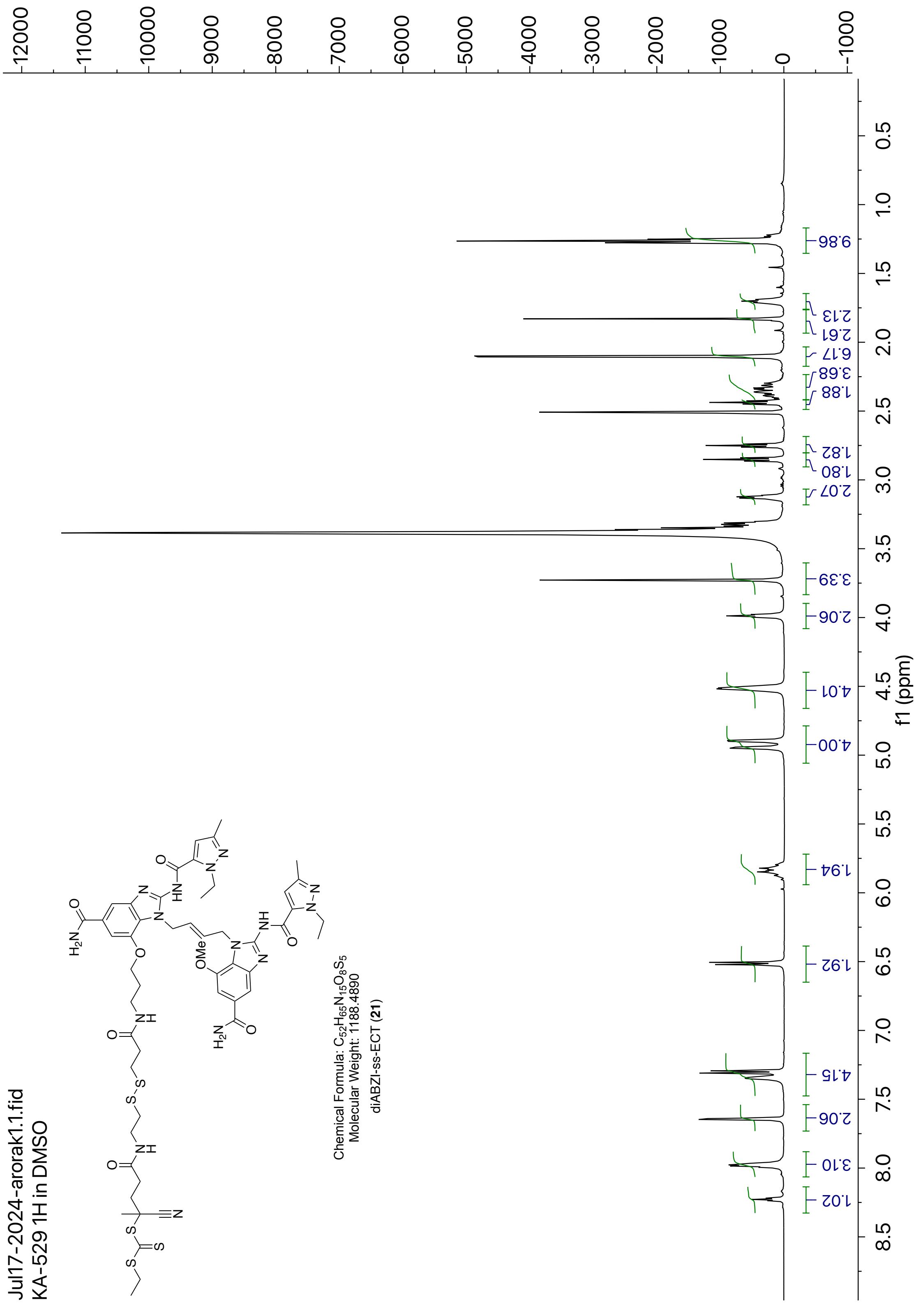


**Figure S23**: [1]H NMR of diABZI-ss-ECT (**21**) in DMSO

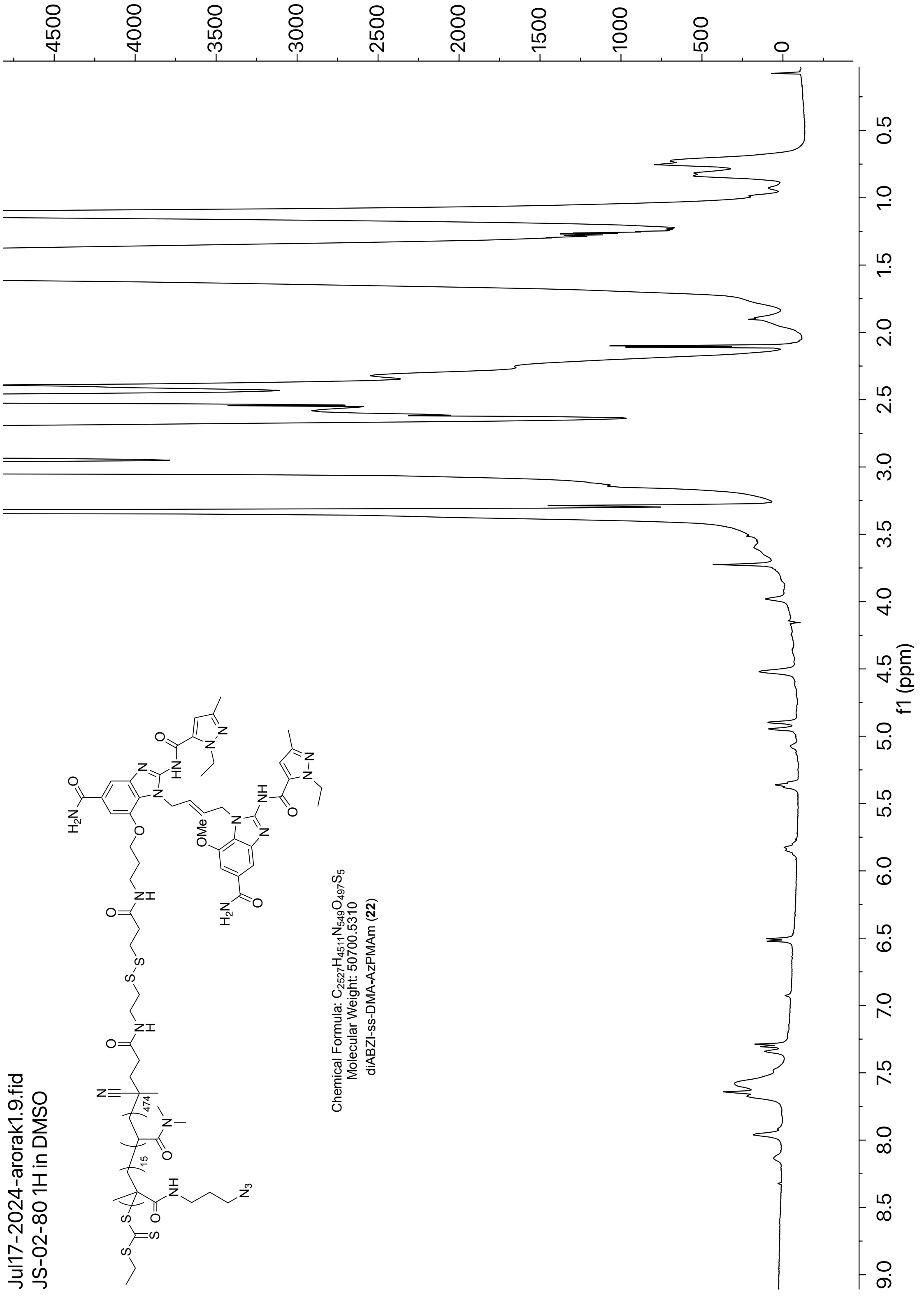


**Figure S24**: $^1$H NMR of compound **22** in DMSO

**Part F: $^{13}$C NMR spectra for new compounds**

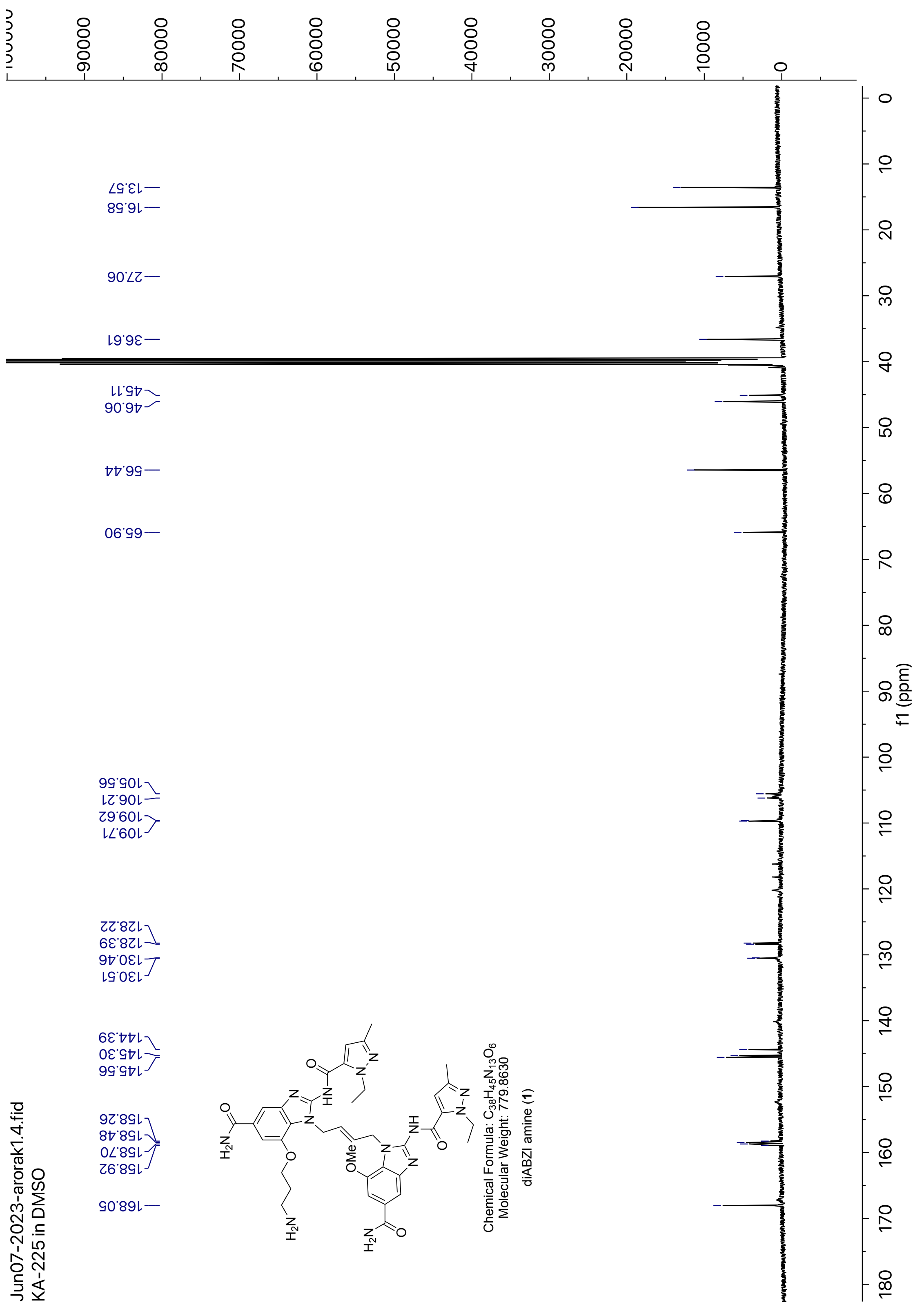


**Figure S25**: $^{13}$C NMR of diABZI amine (**1**) in DMSO

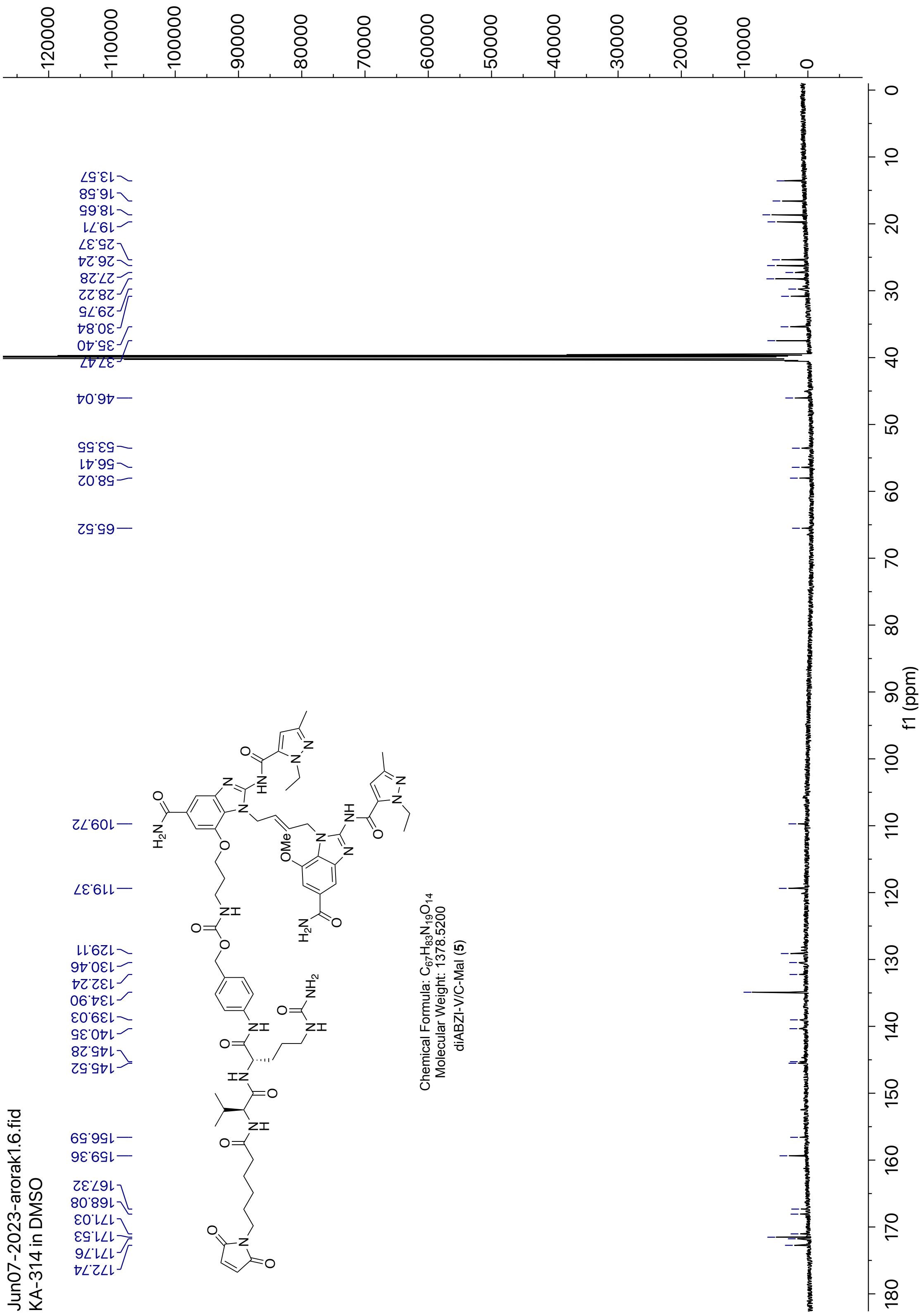


**Figure S26**: $^{13}$C NMR of diABZI-V/C-Mal (**5**) in DMSO

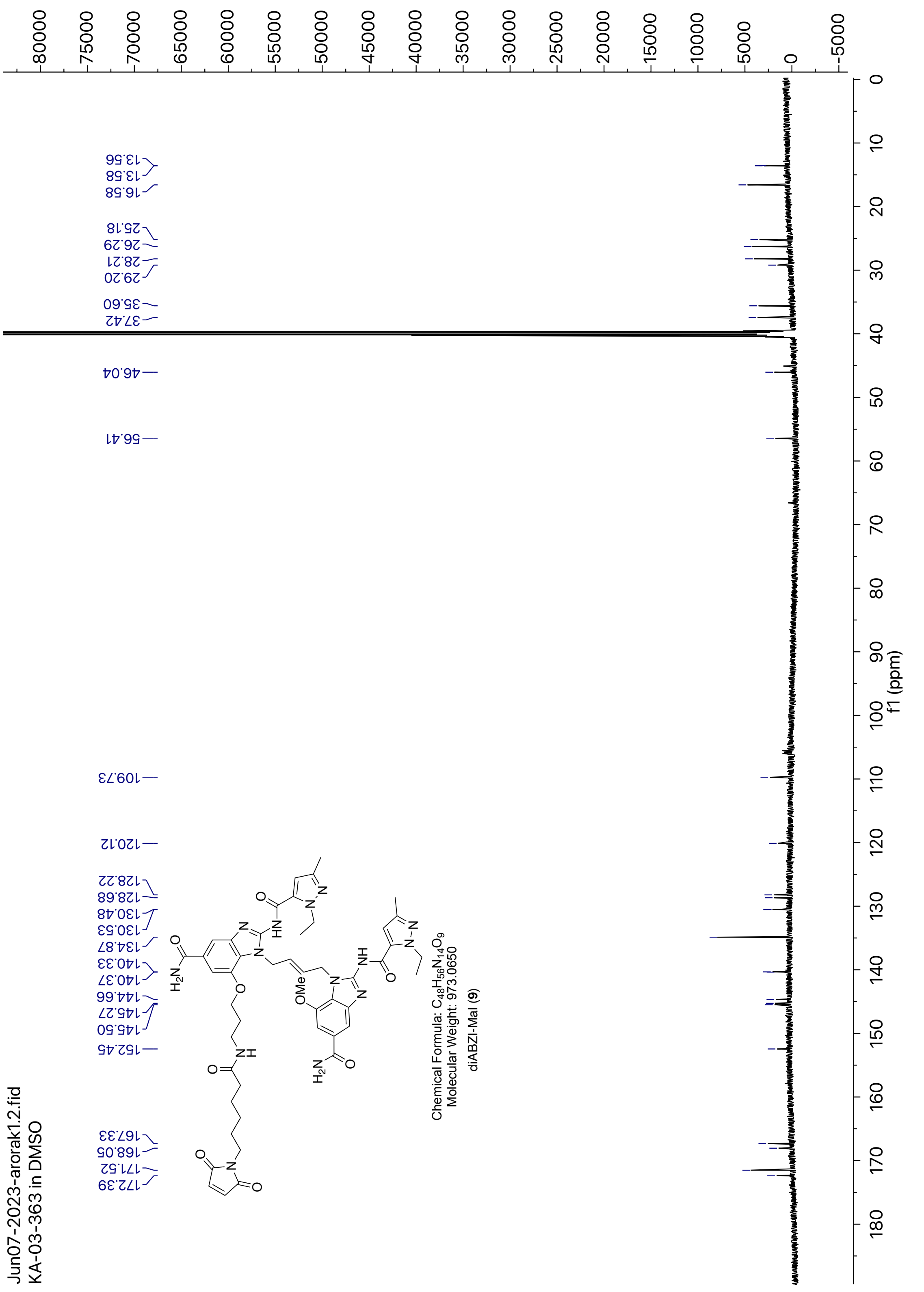


**Figure S27**: $^{13}$C NMR of diABZI-Mal (**9**) in DMSO

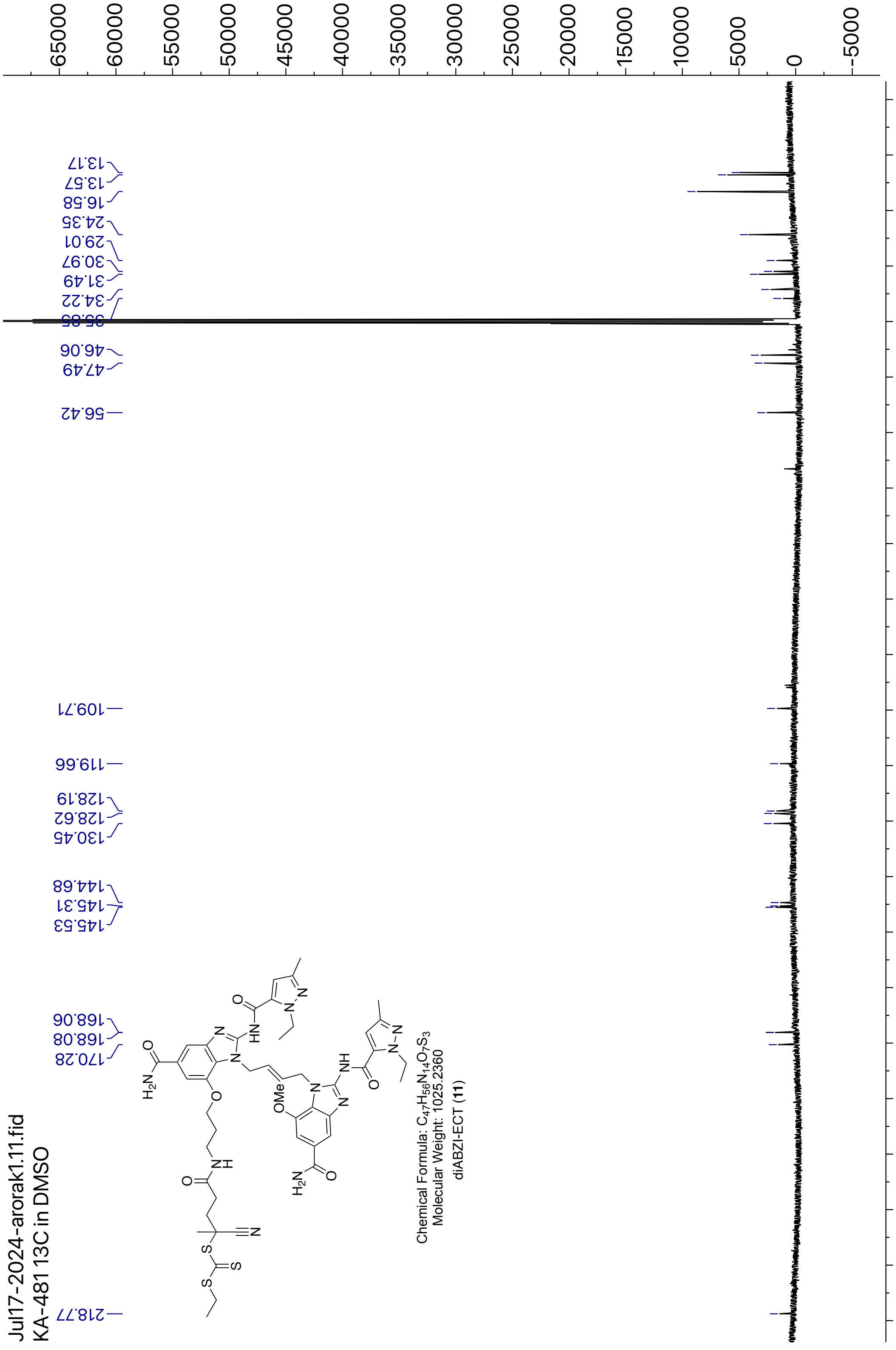


**Figure S28**: $^{13}$C NMR of diABZI-ECT (**11**) in DMSO

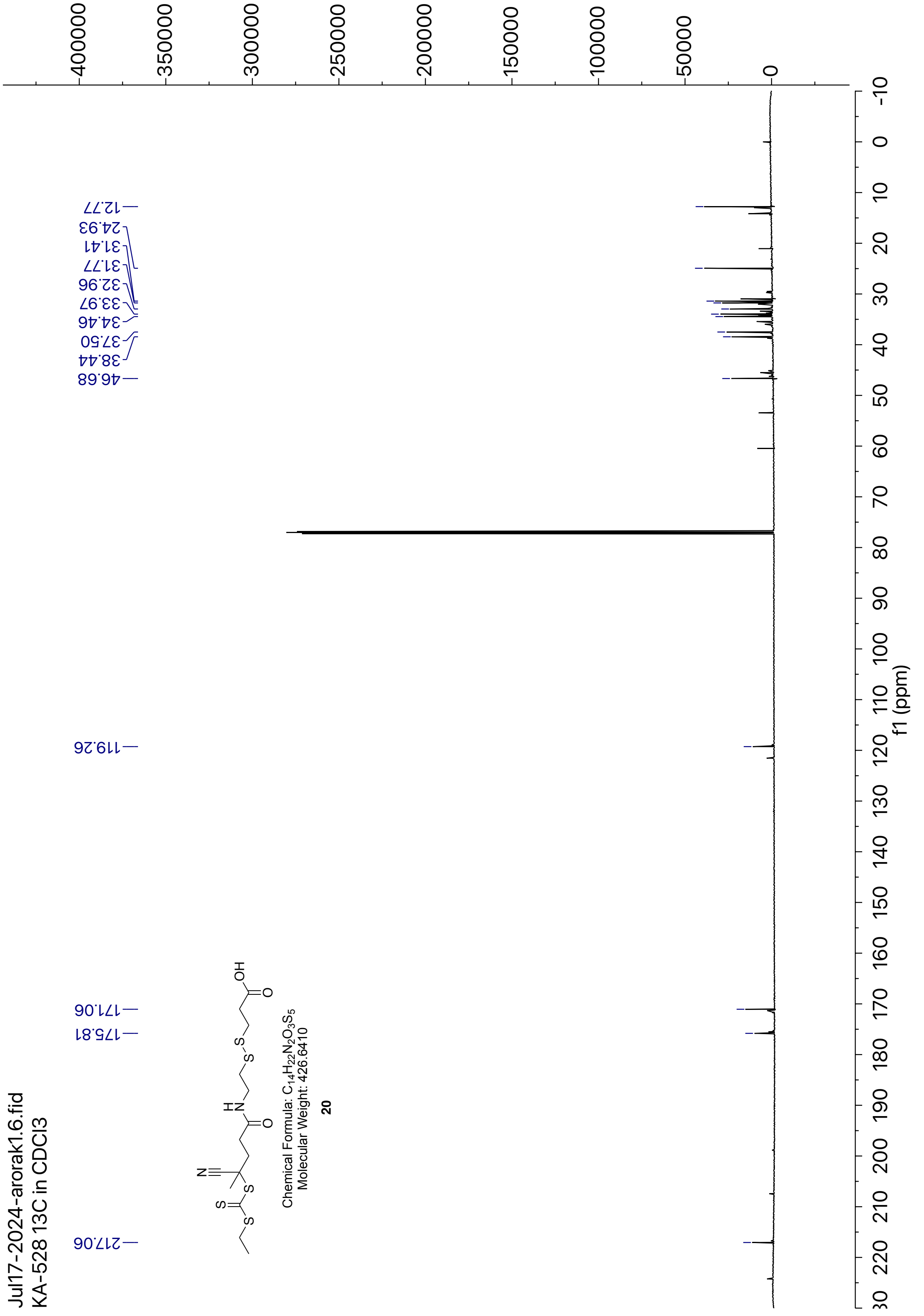


**Figure S29**: $^{13}$C NMR of compound (**20**) in DMSO

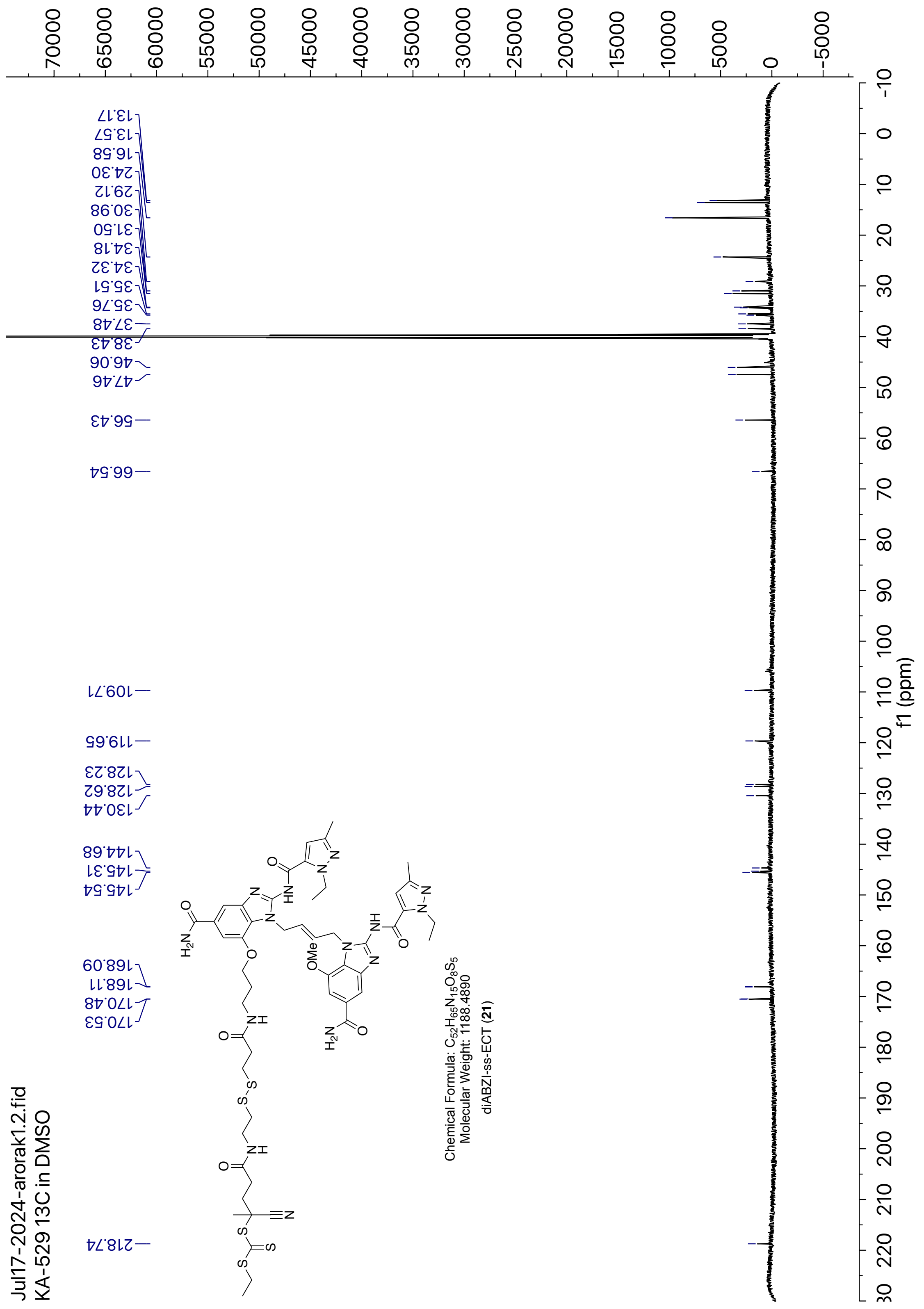


**Figure S30**: $^{13}$C NMR of diABZI-SS-ECT (**21**) in DMSO

### Part G: HRMS spectra for new compounds

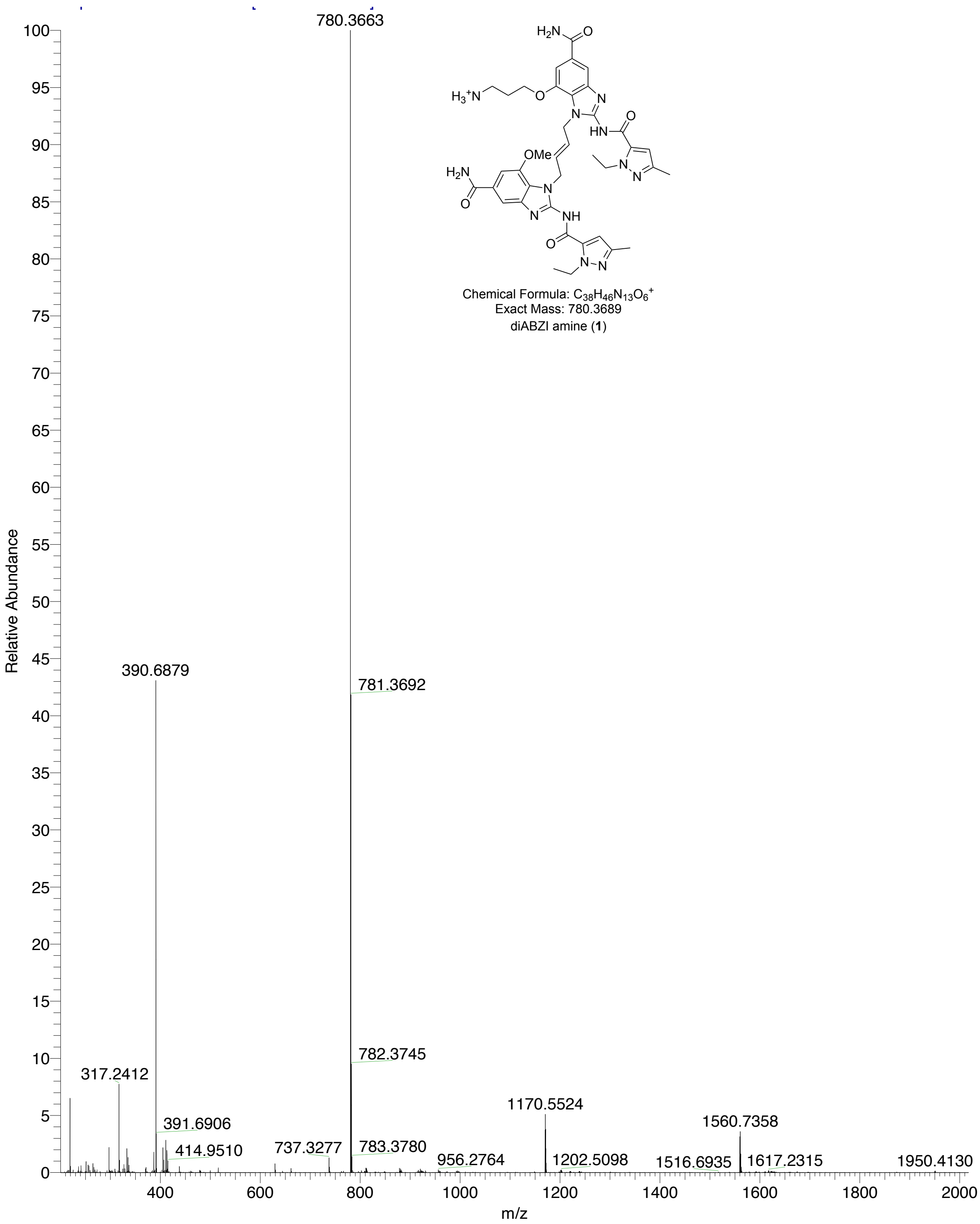


**Figure S31**: Mass spectrum of diABZI amine (**1**)

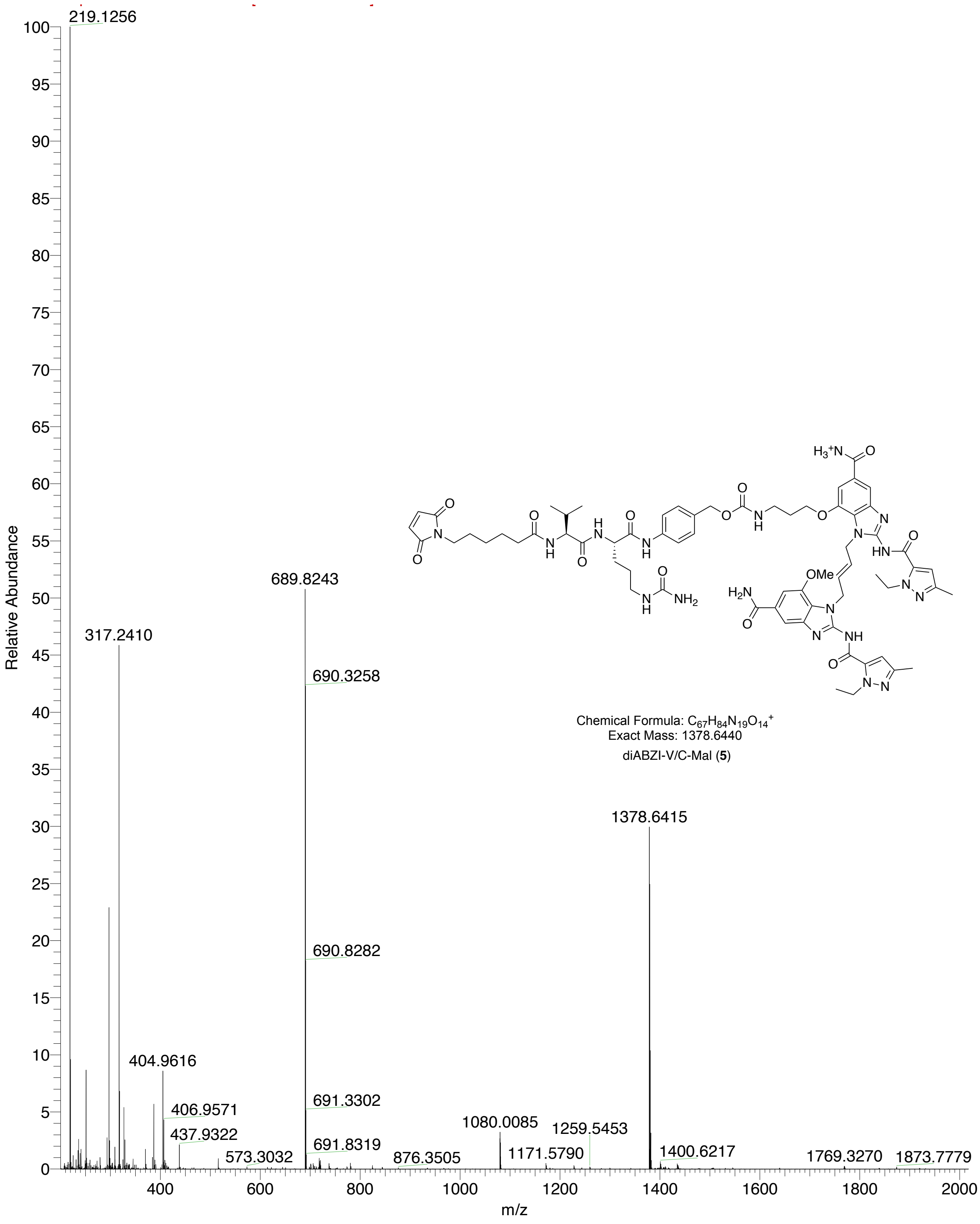


**Figure S32**: Mass spectrum of diABZI-V/C-Mal (**5**)

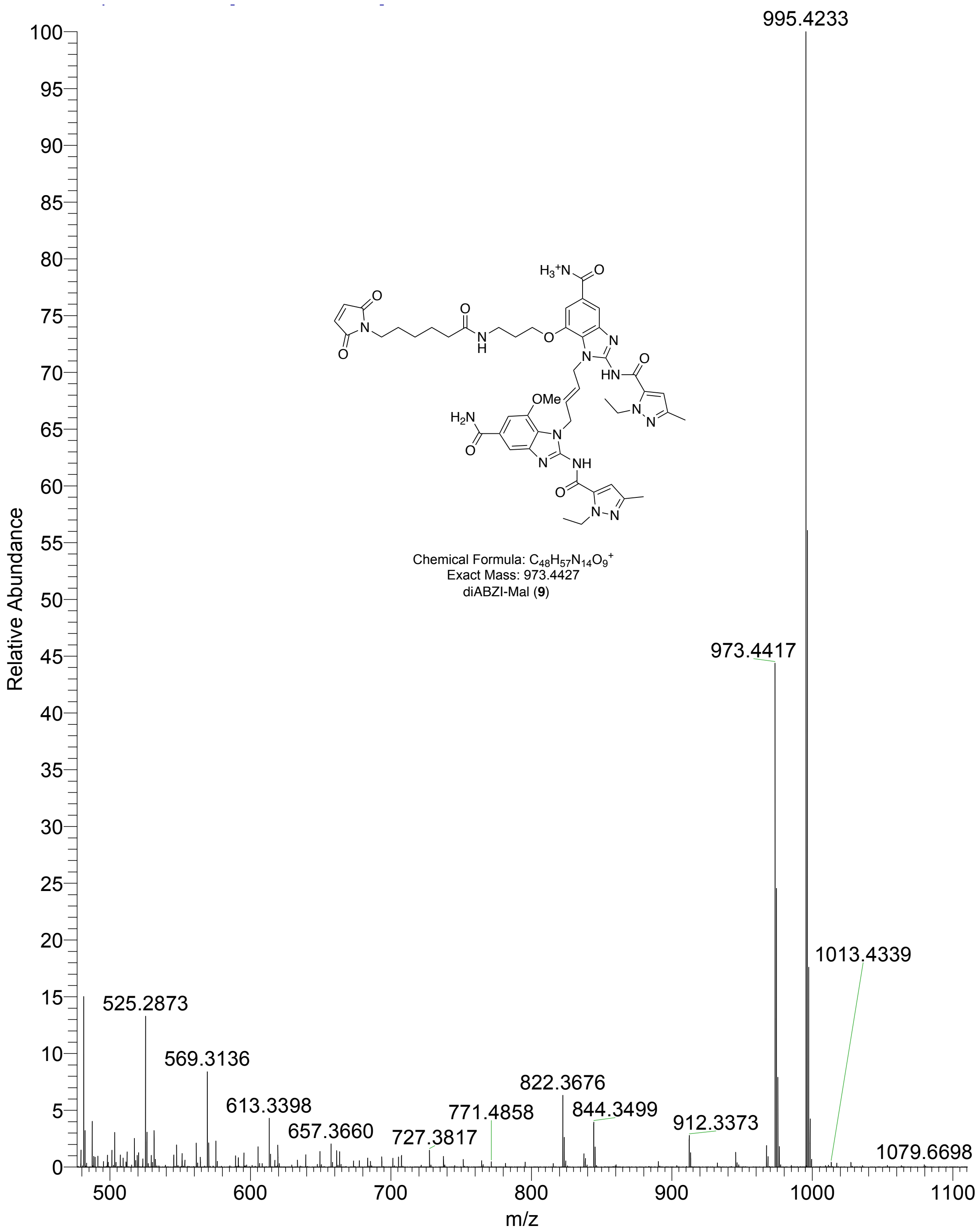


**Figure S33**: Mass spectrum of diABZI-Mal (**9**)

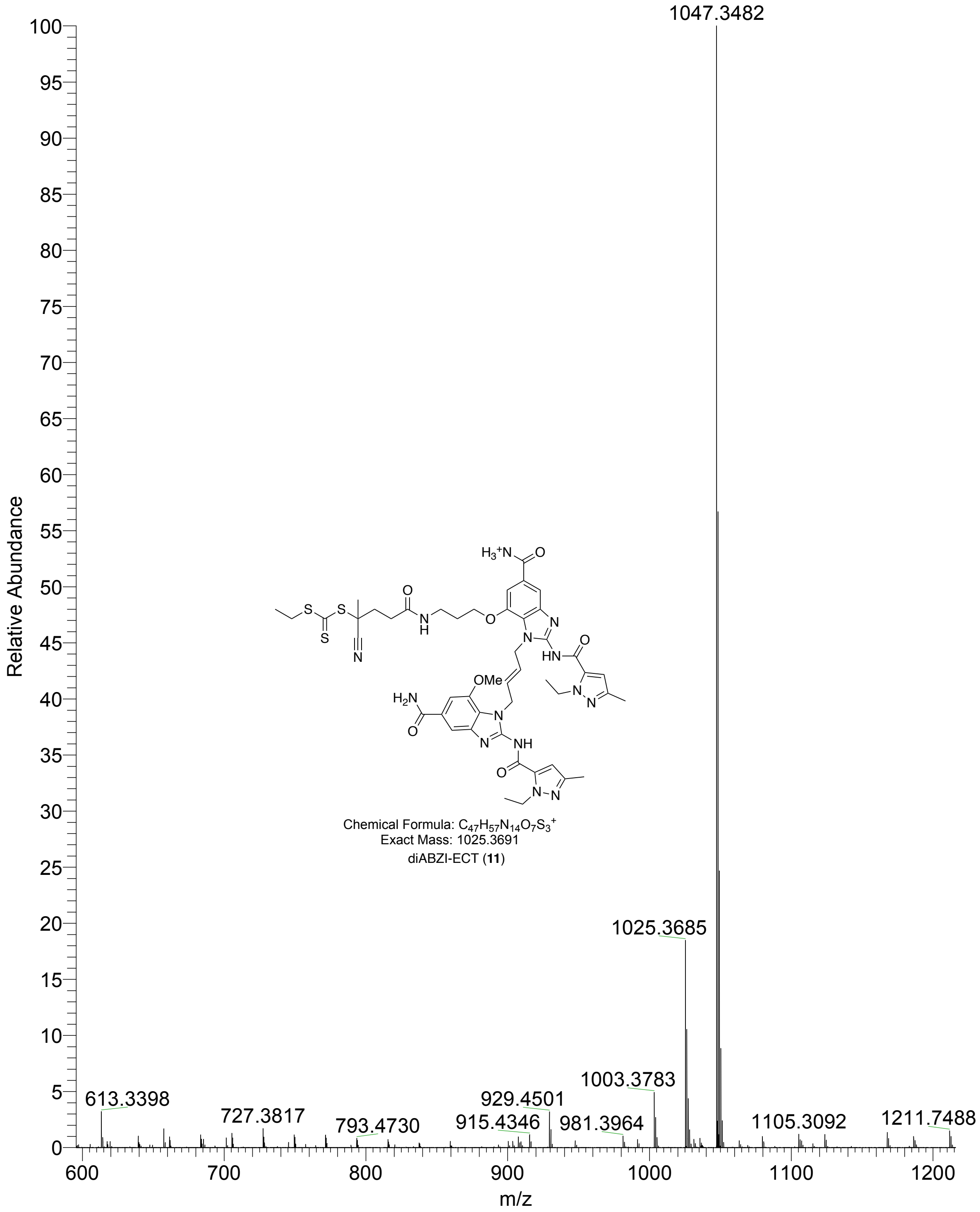


**Figure S34**: Mass spectrum of diABZI-ECT (**11**)

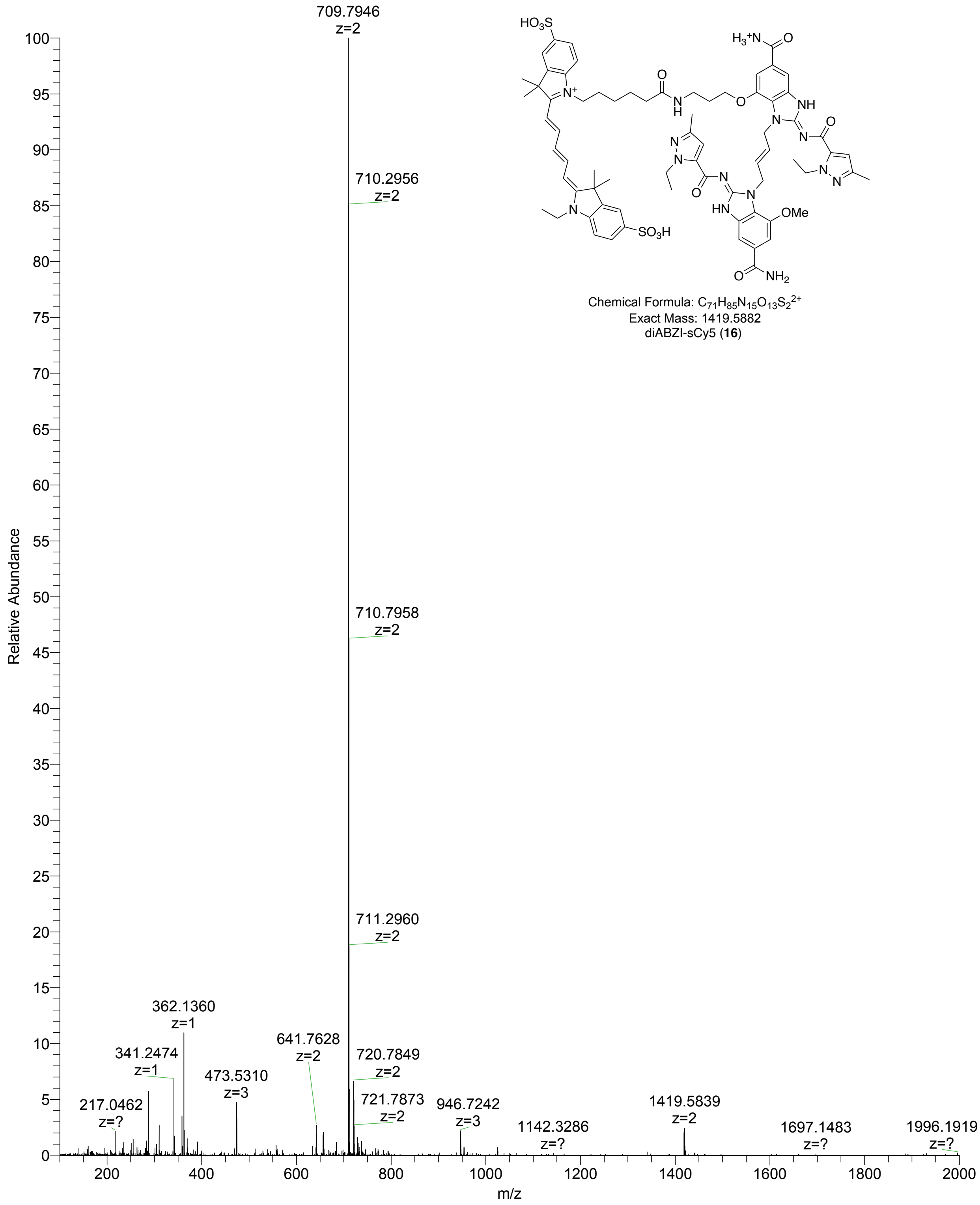


**Figure S35**: Mass spectrum of diABZI-sCy5 (**16**)

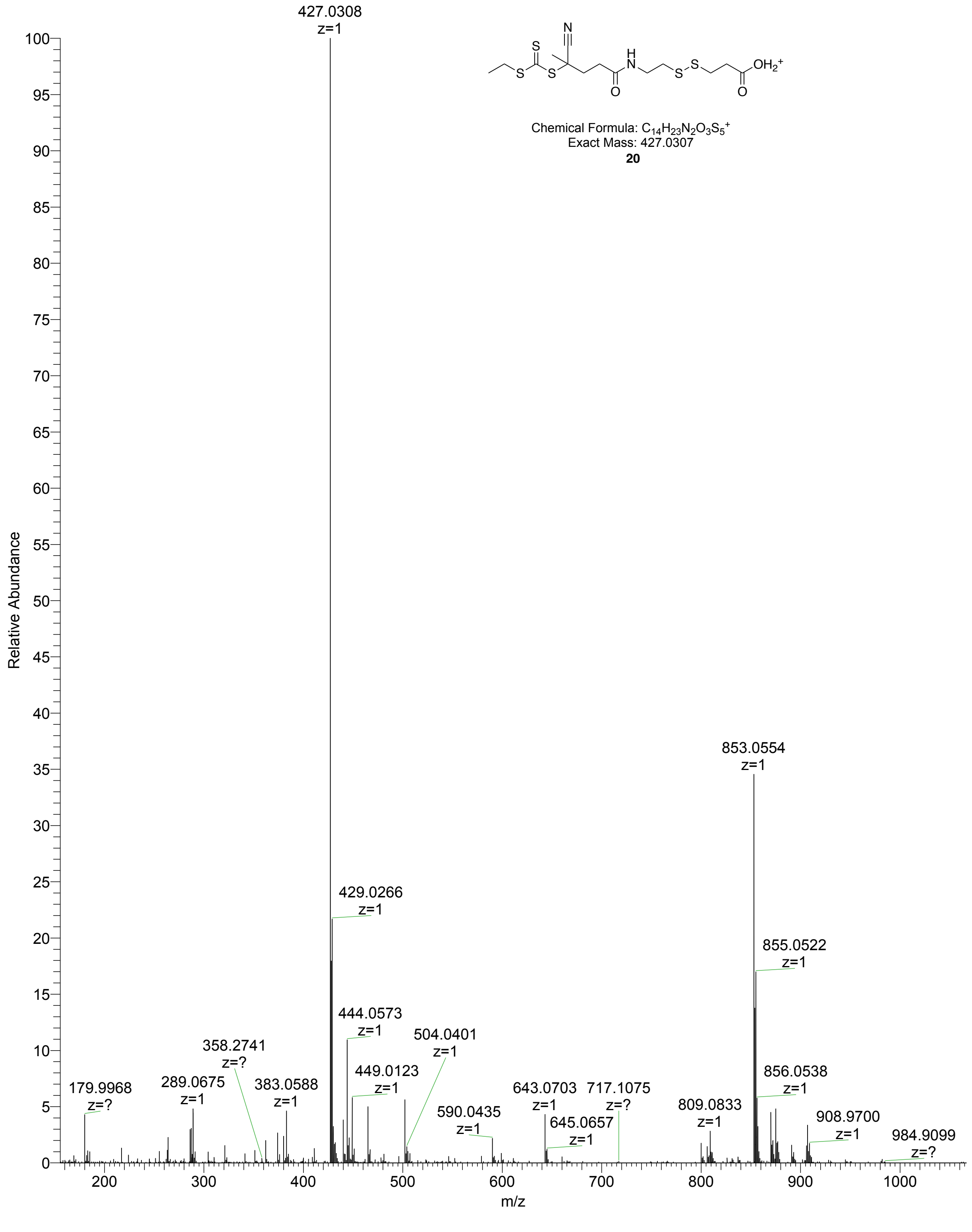


**Figure S36**: Mass spectrum of compound **20**

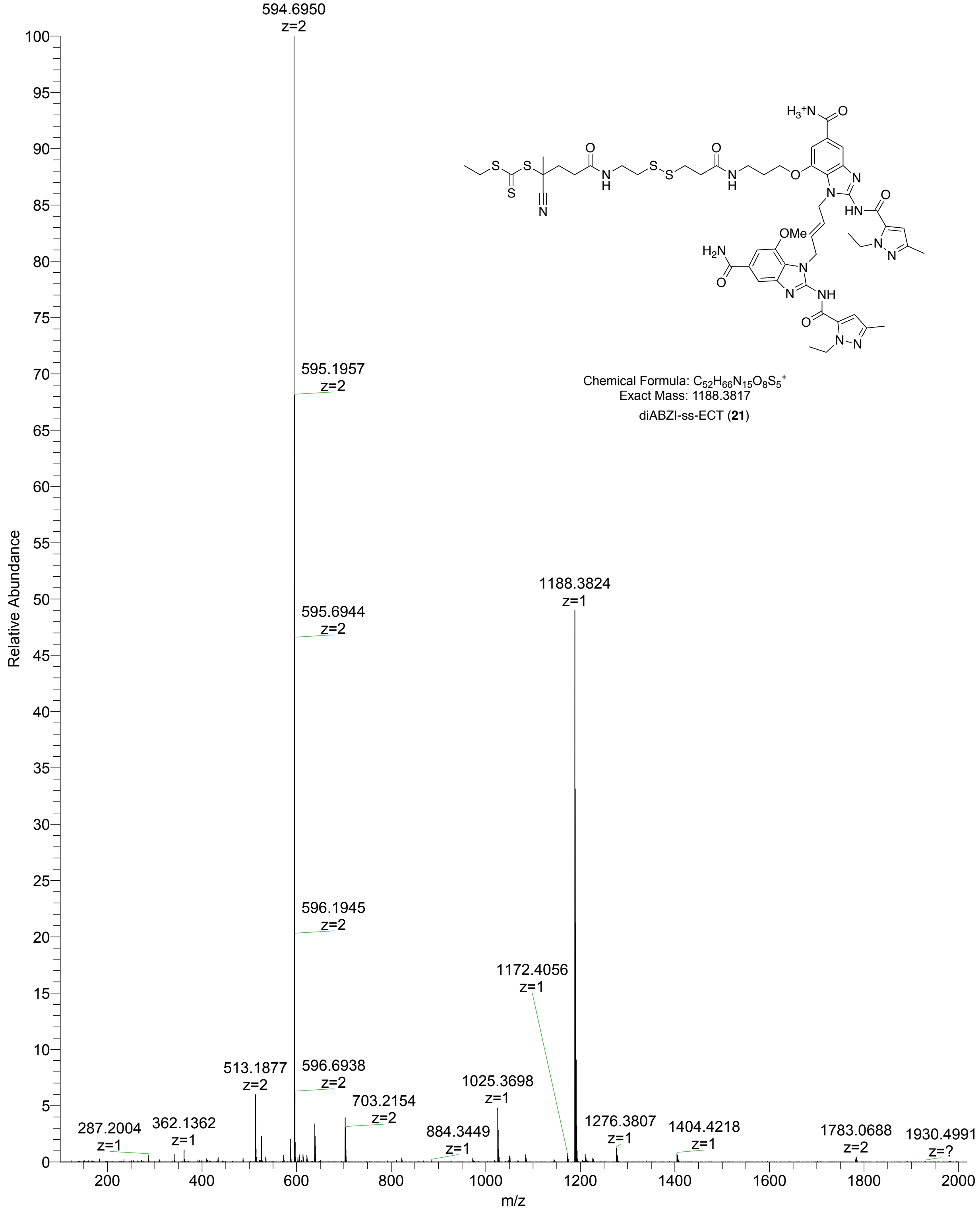


**Figure S37**: Mass spectrum of diABZI-SS-ECT (**21**)

### Part H: GPC/LS of Polymers

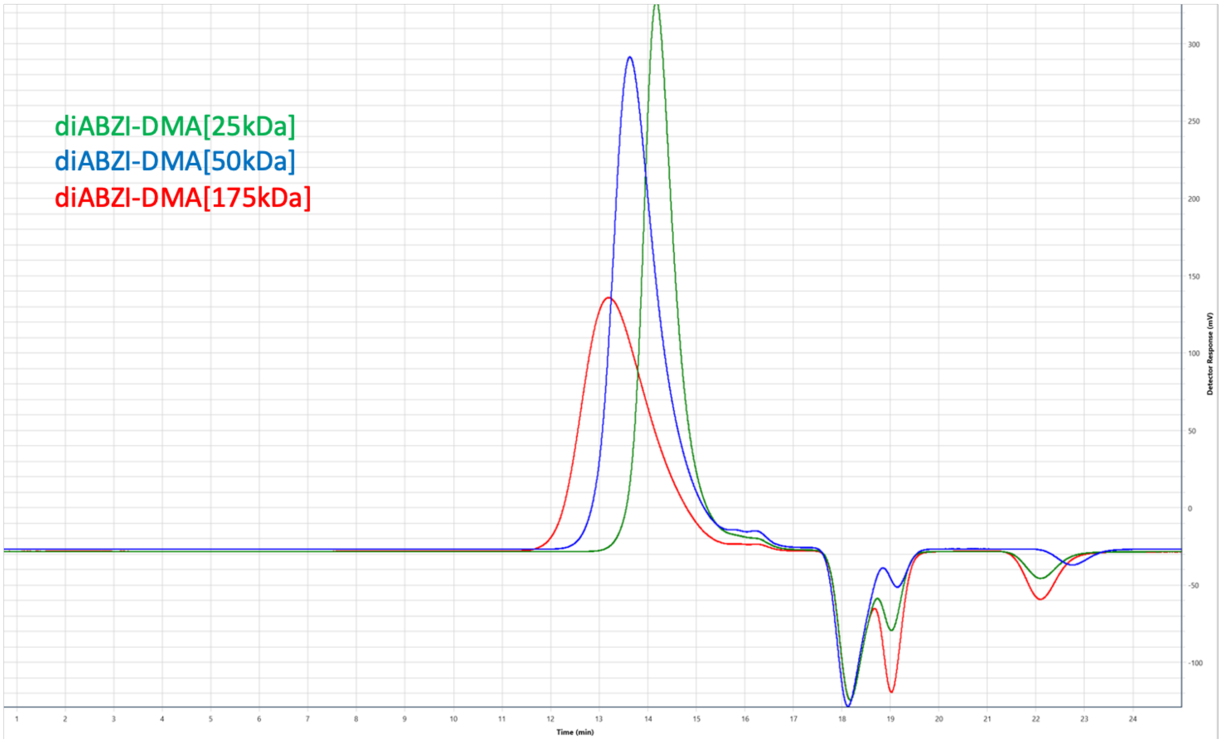


**Figure S38:** GPC/LS Analysis of diABZI-DMA[25kDa, 50kDa, and 175kDa]

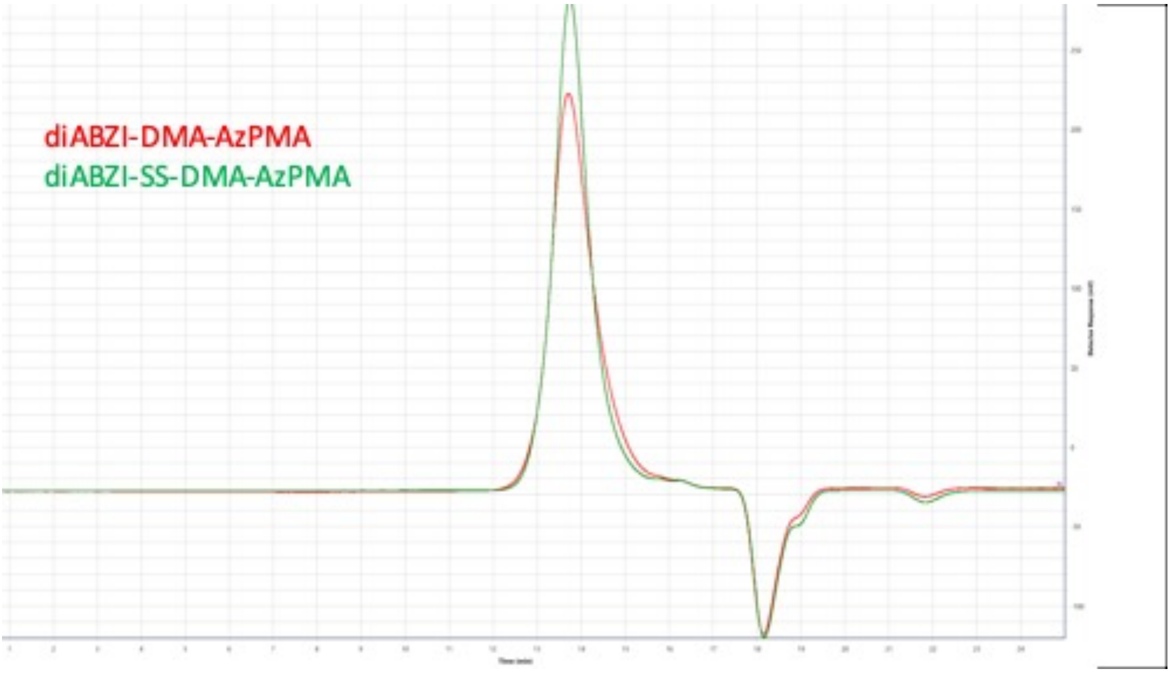


**Figure S39:** GPC/LS Analysis of diABZI-DMA[25kDa, 50kDa, and 175kDa]

**Part I: References**

(1) Sheehy, T.L.; Kwiatkowski, A.J.; Arora, K.; Kimmel, B. R.; Schulman, J.A.; Gibson-Corley, K. N.; Wilson, J.T.; STING-Activating Polymer-Drug Conjugates for Cancer Immunotherapy. *bioRxiv* **2024**, 2024.2003.2023.585817. DOI: 10.1101/2024.03.23.585817.
(2) Blanchard, S.; Coats, J. Process for the preparation of peptide drug linker compounds. WO2019108797, 2019.
(3) van Dongen, S. F. M.; Clerx, J.; Norgaard, K.; Bloemberg, T. G.; Cornelissen, J. J. L. M.; Trakselis, M. A.; Nelson, S. W.; Benkovic, S. J.; Rowan, A. E.; Nolte, R. J. M. A clamp-like biohybrid catalyst for DNA oxidation. *Nat. Chem.* **2013**, *5* (11), 945-951. DOI: 10.1038/nchem.1752.
(4) Qiu, J.; Meng, F.; Wang, M.; Huang, J.; Wang, C.; Li, X.; Yang, G.; Hua, Z.; Chen, T. Recyclable DMAP-Functionalized polymeric nanoreactors for highly efficient acylation of alcohols in aqueous systems. *Polymer* **2021**, *222*, 123660. DOI: 10.1016/j.polymer.2021.123660.
(5) Goor, O. J. G. M.; Keizer, H. M.; Bruinen, A. L.; Schmitz, M. G. J.; Versteegen, R. M.; Janssen, H. M.; Heeren, R. M. A.; Dankers, P. Y. W. Efficient Functionalization of Additives at Supramolecular Material Surfaces. *Adv. Mater. (Weinheim, Ger.)* **2017**, *29* (5), n/a. DOI: 10.1002/adma.201604652. Luzuriaga, M. A.; Welch, R. P.; Dharmarwardana, M.; Benjamin, C. E.; Li, S.; Shahrivarkevishahi, A.; Popal, S.; Tuong, L. H.; Creswell, C. T.; Gassensmith, J. J. Enhanced Stability and Controlled Delivery of MOF-Encapsulated Vaccines and Their Immunogenic Response In Vivo. *ACS Appl. Mater. Interfaces* **2019**, *11* (10), 9740-9746. DOI: 10.1021/acsami.8b20504.
(6) Yang, F.; Li, Z.; Li, S.; Wang, J.; Zheng, X. Compound biotin-tripolyethylene glycol-S-S-dibenzazepine cyclooctyne, its preparation method and application. CN114106012, 2022.
